# The genomic landscapes of desert birds form over multiple time scales

**DOI:** 10.1101/2022.03.07.483329

**Authors:** Kaiya Provost, Stephanie Yun Shue, Meghan Forcellati, Brian Tilston Smith

## Abstract

Spatial models show that genetic differentiation between populations can be explained by factors ranging from geographic distance to environmental resistance across the landscape. However, genomes exhibit a landscape of differentiation, which could indicate that multiple spatial models better explain divergence in different portions of the genome. We test whether alternative geographic predictors of intraspecific differentiation vary across the genome in ten bird species that co-occur in Sonoran and Chihuahuan Deserts of North America. Using population-level genomic data, we characterized the genomic landscapes across species and modeled five predictors that represented historical and contemporary mechanisms. The characteristics of genomic landscapes differed across the ten species, influenced by varying levels of population structuring and admixture between deserts. General dissimilarity matrix modeling indicated that the best-fit models differed from the whole genome and partitions along the genome. Almost all of the historical and contemporary mechanisms were important in explaining genetic distance, but particularly historical and contemporary environment, while contemporary abundance, position of the barrier to gene flow, and distance explained relatively less. Individual species have significantly different patterns of genomic variation. These results illustrate that the genomic landscape of differentiation was influenced by alternative geographic factors operating on different portions of the genome.

## Introduction

Levels of nucleotide diversity and the degree of differentiation both vary across genomes (e.g., Ellegren et al., 2012; Li and Ralph, 2019). These so-called genomic landscapes are produced by variable processes including ones intrinsic to the genome (meiotic recombination, mutation) and those extrinsic (introgression, selection, and drift). Fluctuating levels of genetic diversity across the genome have been shown to be associated with recombination rate indicating that linked selection reduces variation (Burri et al., 2015, Martin et al., 2019, Johri et al., 2020). Likewise mutation rates and coalescent times are all known to covary with levels of differentiation across the genome (Nosil and Schluter, 2011; Benzer, 1961; Hodgkinson and Eyre-Walker, 2011). In contrast to intrinsic processes which are primarily mediated by genomic properties, extrinsic processes are mediated through interactions with the adaptive and demographic factors operating across space. The locations of speciation genes are found to be associated with genomic differentiation (Nosil and Schluter, 2011; Benzer, 1961; Hodgkinson and Eyre-Walker, 2011). Despite evidence of the patterns and processes driving a heterogeneous genomic landscape (e.g., Li and Ralph, 2019, Wang et al., 2020), studies examining the geographic predictors of genetic differentiation often use only single summary statistics to represent the entirety of the genome, for example using a single F_ST_ value for comparing whole populations. Clarifying the relationship between the heterogeneity of the genomic landscape and geographic predictors of differentiation will elucidate how intraspecific variation arises in the complex physical landscape.

The spatial processes attributed to population differentiation operate over historical through contemporary time scales; herein, we focus on five as examples. An atemporal manifestation of historical isolation, such as isolation by barrier(s) (IBB; *sensu* Mayr, 1942) can occur, where population differentiation is best predicted by a landscape feature. Over shallower evolutionary scales, non-random mating with individuals in closer geographic proximity can cause genetic differentiation by isolation by distance (IBD; Wright, 1943). IBD has been shown to impact taxa at both small (e.g., Aguillon et al., 2017) and large geographic scales (e.g., Rethelford, 2004). Geographic distances alone may not be the best predictors of differentiation because adaptation to local climatic conditions causes selection to generate structuring across environmental gradients, which is known as isolation by environment (IBE; Wang and Bradburd, 2014, Myers et al., 2019, Berg et al., 2015; Zamudio et al., 2016). These two factors have been shown to work concurrently with one another in many groups (Sexton et al., 2014). Because local environmental conditions change rapidly, for example due to species turnover or succession (Phillips, 1996, Nuvoloni et al., 2016), associations between differentiation and environment are likely more recent phenomena than historical associations. The increased availability of ecological data for many organisms, such as census data, allows for testing even shallower associations with genetic structuring across the landscape. Contemporary demographic data can be used to test whether genetic differences are associated with abundance troughs that restrict gene flow (Barton and Hewitt, 1981; Hewitt, 1989; Barrowclough et al., 2005; referred to herein as “IBA” for brevity). Though it is often assumed that abundance and niche occupancy are correlated due to the link with suitable habitat (Holt, 2009), this is not necessarily borne out (Waldock et al., 2021) and as such we estimate these factors separately. Local population size is also known to be a strong driver of genetic structure, especially when compounded with environmental change determining local suitability (Weckworth et al., 2013). Finally, population history is often linked to Pleistocene glacial cycles that shifted and fragmented distributions. An association of genome-wide structuring linked to population fragmentation can be tested under a scenario where genetic distances are modeled against paleo-climatic suitability (Vasconcellos et al., 2019; Moreira et al., 2020; referred to herein as “IBH” for brevity).

While the focus of these models is often on genetic variation, they can also be applied to phenotypic variation (e.g., Moreira et al., 2020). Phenotypic variation is often the product of many loci with little effect (Zeng, 1994). As such, looking directly at phenotype can help reveal whether a particular process is associated with trait variance. Examining the genomic landscape in the context of these alternative geographic models will provide evidence for how factors of varying temporal resolutions influence the peaks and valleys of differentiation. To investigate how landscape features impact genotypic and phenotypic variation across space, we use an archetypical assemblage of co-distributed birds distributed across the Sonoran and Chihuahuan Deserts of the southwestern USA and northern Mexico.

Here we characterize the genomic landscapes of birds occurring across the Sonoran and Chihuahuan Deserts and test the relative effect of alternative geographic models in predicting patterns of intraspecific differentiation. To do this, we integrate population-level whole-genome resequencing, specimen-based morphometrics, and comparative sampling across ten co-distributed species that occur across the deserts. We hypothesize that the best-predictors of genetic diversity will vary across species and different partitions of the data, reflecting the multiple extrinsic factors that structure variation across the genomic landscape (Supplementary Figure 1). Alternatively, species could show homogeneous patterns either by the same geographic modeling predicting differentiation in windows across the whole genome or by species exhibiting congruent genomic landscapes shaped by the same geographic barrier. We further evaluate whether summary statistics, reflective of alternative evolutionary processes, could explain alternative geographic predictors of genomic landscapes. This comparative framework will provide resolution to the extent at which peaks and valleys of the genomic landscape correspond to historical through contemporary factors.

## Results

### Summary of Genomic Data

We sequenced the genomes of 221 individuals across 10 focal species of passerine birds distributed in the Sonoran and Chihuahuan Deserts (Figure 1). Individuals varied in their coverage across the genome. We created three datasets to address this variation in downstream analyses: a complete dataset of all individuals, a dataset where individuals with greater than 75% missing base pairs were removed, and a dataset where individuals with greater than 50% missing base pairs were removed; we call these the 100%, 75%, and 50% missing data partitions, respectively. We found that the three missing data partitions did not vary substantially with respect to coverage at non-missing sites or number of SNPs. As such, here we describe the results for the complete dataset (for the 75% and 50% missing data partitions, see Supplementary Information). We recovered sequences with a mean coverage of 2.9 per individual (range 0.4–8.8), 6–25 million reads per individual, and 5–28 million SNPs per species. Mean X coverage within species ranged from 2.1x– 4.2x, with *Phainopepla nitens* having the lowest coverage and *Melozone fusca* the highest. The average missing data per species ranged from 48–64%. Across individuals, missing data ranged from 13–93% with a mean of 53% (Table 1).

**Figure 1:**
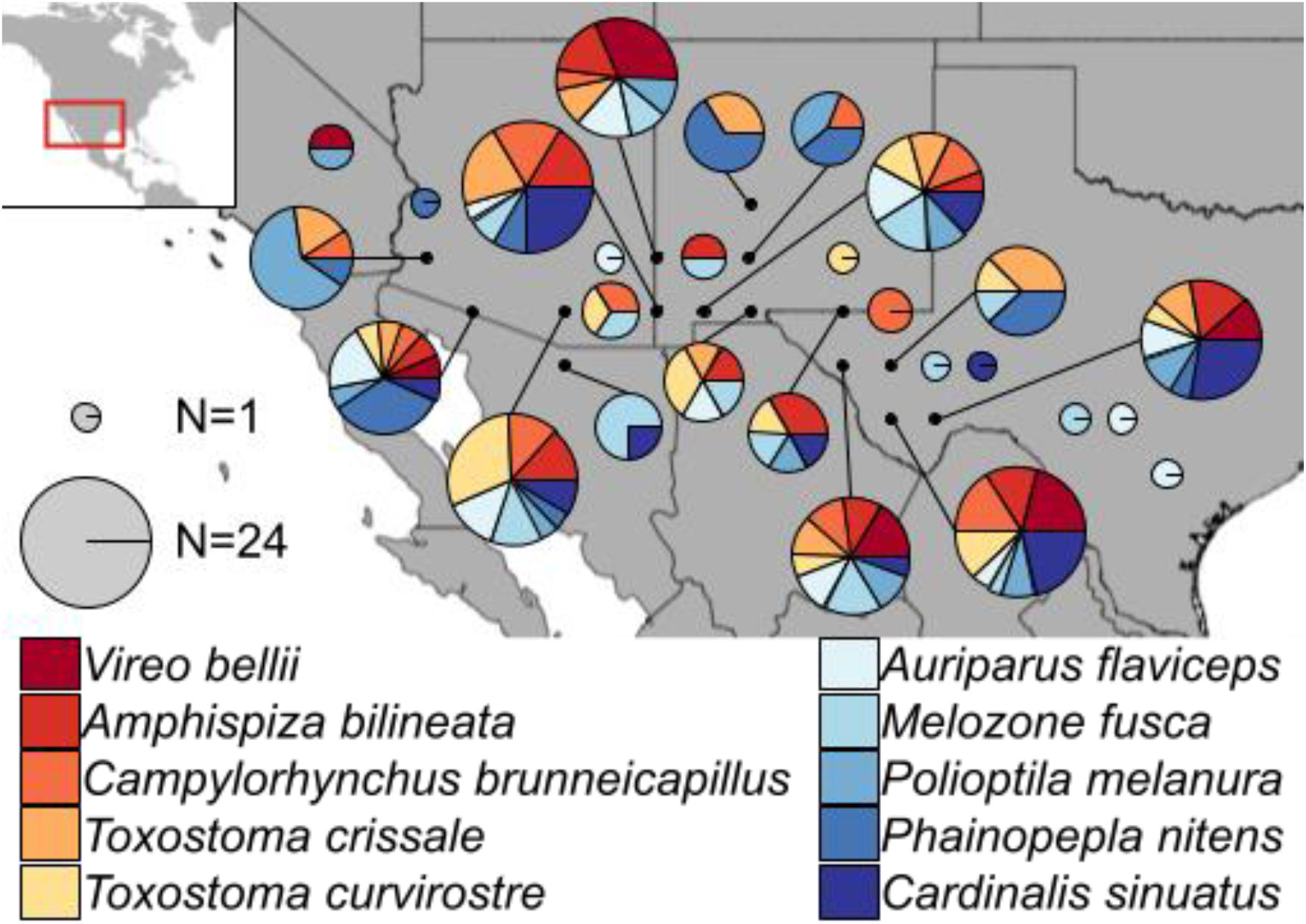
Sampling map of study across southwestern North America. Localities are given with black points (with latitudes/longitudes of specimens rounded to nearest degree). Pie charts show the number (radius of pie chart) and species identity (color of slices) of specimens used from that area. Large pie charts are linked to their locality with a black line.

**Table 1:**
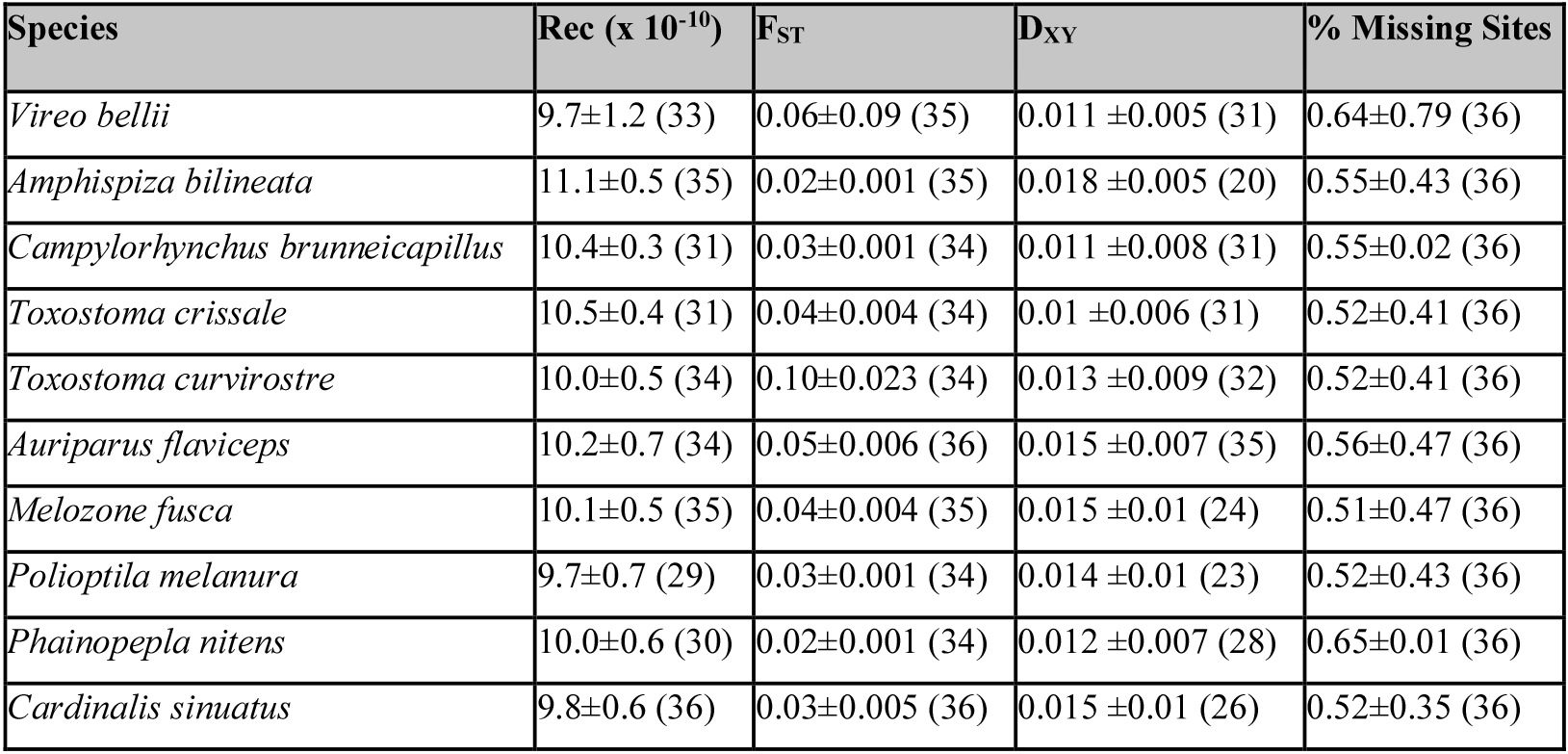
Chromosome-wise values for the recombination rate, F_ST_, D_XY_, and proportion of missing data per each species. Values given as mean±standard deviation (number of chromosomes). These are calculated by weighting all chromosome means equally; for size-weighted values see Supplementary Table 1. Note that the number of chromosomes was based on the pseudo-chromosomes we generated, with a maximum of 36. “Rec”=population recombination rate, or rho. Values are given for the complete dataset; for the 50% and 75% values, see Supplementary Table 1.

### Recombination Rate

Mean recombination rates for the entire genome estimated using ReLERNN (Adrion et al., 2020) ranged from 8.9–12.8 × 10^−10^ c/bp (where c is the probability of a crossover) across species. Correlations between species in mean recombination across chromosomes range from -0.57 to 0.53 (mean±SD 0.02±0.25). Correlations between species in mean recombination at the same genomic positions ranged from -0.33 to 0.43 (mean±SD -0.01±0.22). Recombination rate was not associated with log corrected chromosome size (p=0.82).

### Lostruct outliers and F_ST_ outliers

We divided the genome into three kinds of partitions. First, we analyzed chromosomes independently. Second, we identified high F_ST_ outliers (by calculating the z-score of F_ST_ values across the genome within species and retaining only those more than 5 standard deviations above the mean) and analyzed those. Finally, we performed a multidimensional scaling (MSDS) analysis the using R package lostruct version 0.0.0.9000 (Li and Ralph, 2019), which subdivided genomes into four partitions, three outliers (LS1, LS2, LS3) and one non-outlier partition (Figure 2; Supplementary Figure 2). Note that outlier groupings are not analogous across taxa. On average across all species 85.3% of labeled values were non-outliers, and ∼4.88% each were LS1, LS2, and LS3.

**Figure 2:**
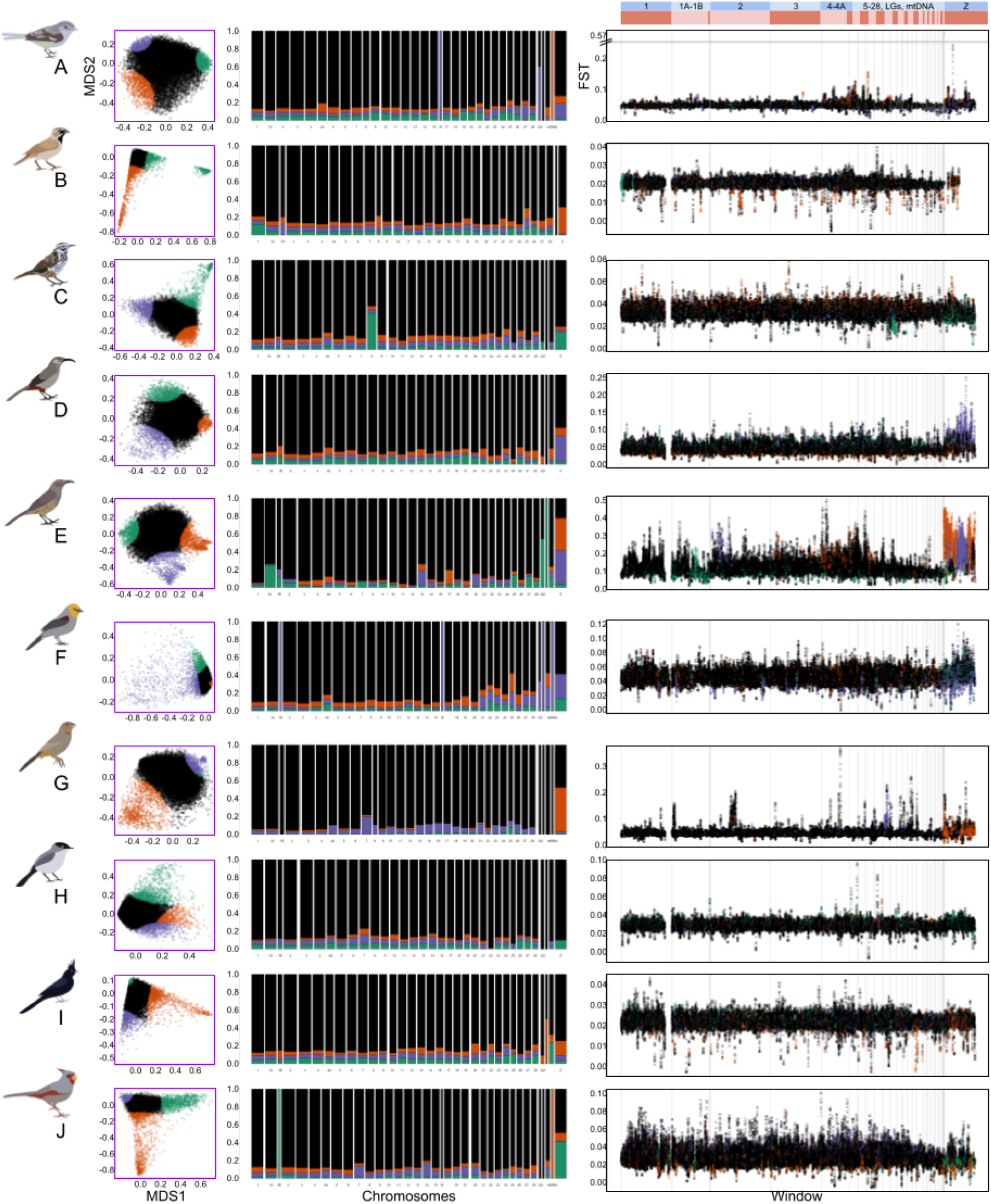
Lostruct partitions vary across species and across chromosomes. Species are as follows: A) *Vireo bellii*, B) *Amphispiza bilineata*, C) *Campylorhynchus brunneicapillus*, D) *Toxostoma crissale*, E) *Toxostoma curvirostre*, F) *Auriparus flaviceps*, G) *Melozone fusca*, H) *Polioptila melanura*, I) *Phainopepla nitens*, J) *Cardinalis sinuatus*. Left column: Multidimensional scaling coordinate 1 (x-axis) vs 2 (y-axis) for each species, with outlier points highlighted in orange, green, and purple as different partitions, and non-outlier points in black. Middle column: proportion of chromosomes assigned to LS1 (orange), LS2 (green), LS3 (purple), and non-outlier (black) lostruct partitions. Width of bars approximately proportional to length of each chromosome. Right column: F_ST_ values for windows across the genome, colored by lostruct partition (orange, green, purple, black). Each window represents one 100,000 base pair wide section of the genome, with subsequent windows overlapping by 10,000 base pairs Note that F_ST_ values are not on the same scale for all taxa. Chromosomes separated by gray lines, with legend at the top. Species images are not to scale.

The number of highly differentiated regions in the genome varied between species. F_ST_ outlier analysis across datasets with different levels of missing data found largely congruent results with respect to how many outliers were present across taxa (see Supplementary Information for 75% and 50% datasets). The number of high F_ST_ outliers for the complete dataset ranged from 28– 758 across species (with the total number of windows analyzed per species ranging from 100,733– 113,555). The outlier lostruct partitions identified above (LS1, LS2, LS3) vary in the proportion of the F_ST_ outliers examined (for the complete dataset), ranging from 0.0%–3.4% (mean 0.2%) for peaks. Though not significant, there appears to be a trend where species with generally higher F_ST_ have more high-F_ST_ outliers identified.

### Population differentiation

Signatures of population structure varied in our ten species. Population differentiation in species ranged from being highly structured among deserts in four species (*T. curvirostre, V. bellii, A. flaviceps*, and *P. melanura*), showing a gradient of structuring with admixture in three (*T. crissale, M. fusca*, and *Cardinalis sinuatus*), or unstructured in the remaining taxa (*A. bilineata, C. brunneicapillus, P. nitens*; Supplementary Figure 3). F_ST_ values for the species within these three groups varied accordingly: highly structured=0.03–0.10; gradient=0.03–0.04; and unstructured=0.02–0.03. Population differentiation estimated from the chromosomal partitions were generally concordant with genome-level patterns, but smaller chromosomes and/or those with fewer SNPs showed different patterns (Figure 3, Figure 4, Supplementary Figure 4).

**Figure 3:**
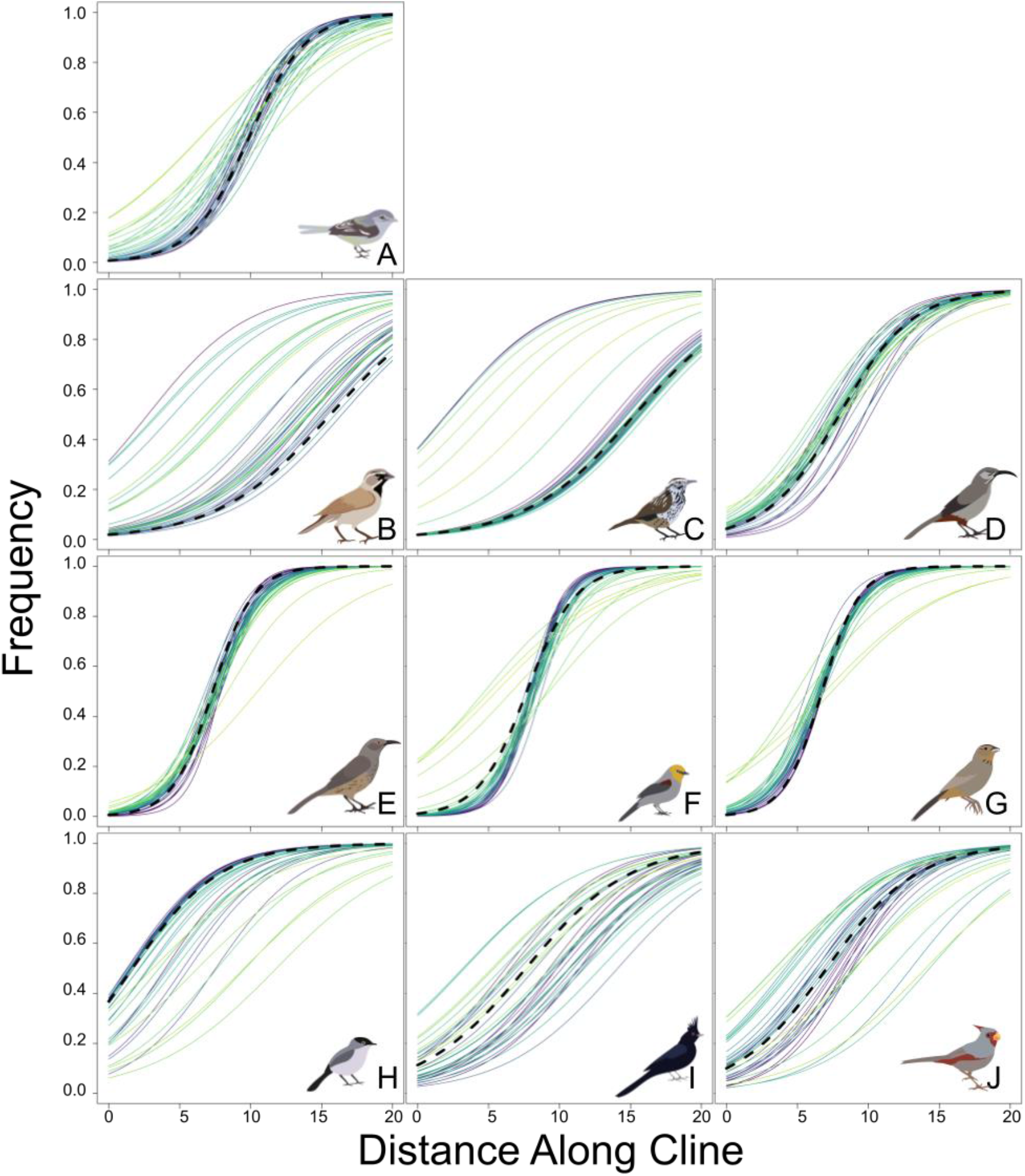
Cline width and center location vary across species and across chromosomes. X-axis shows distance (in degrees longitude) along the sampled area. Y-axis shows the projected cline from population assignments of 0 to 1 in each taxon (panel) and each chromosome (colored lines). Genomes are given by thick dashed black lines. Species are as follows: A) *Vireo bellii*, B) *Amphispiza bilineata*, C) *Campylorhynchus brunneicapillus*, D) *Toxostoma crissale*, E) *Toxostoma curvirostre*, F) *Auriparus flaviceps*, G) *Melozone fusca*, H) *Polioptila melanura*, I) *Phainopepla nitens*, J) *Cardinalis sinuatus*. Species images are not to scale.

**Figure 4:**
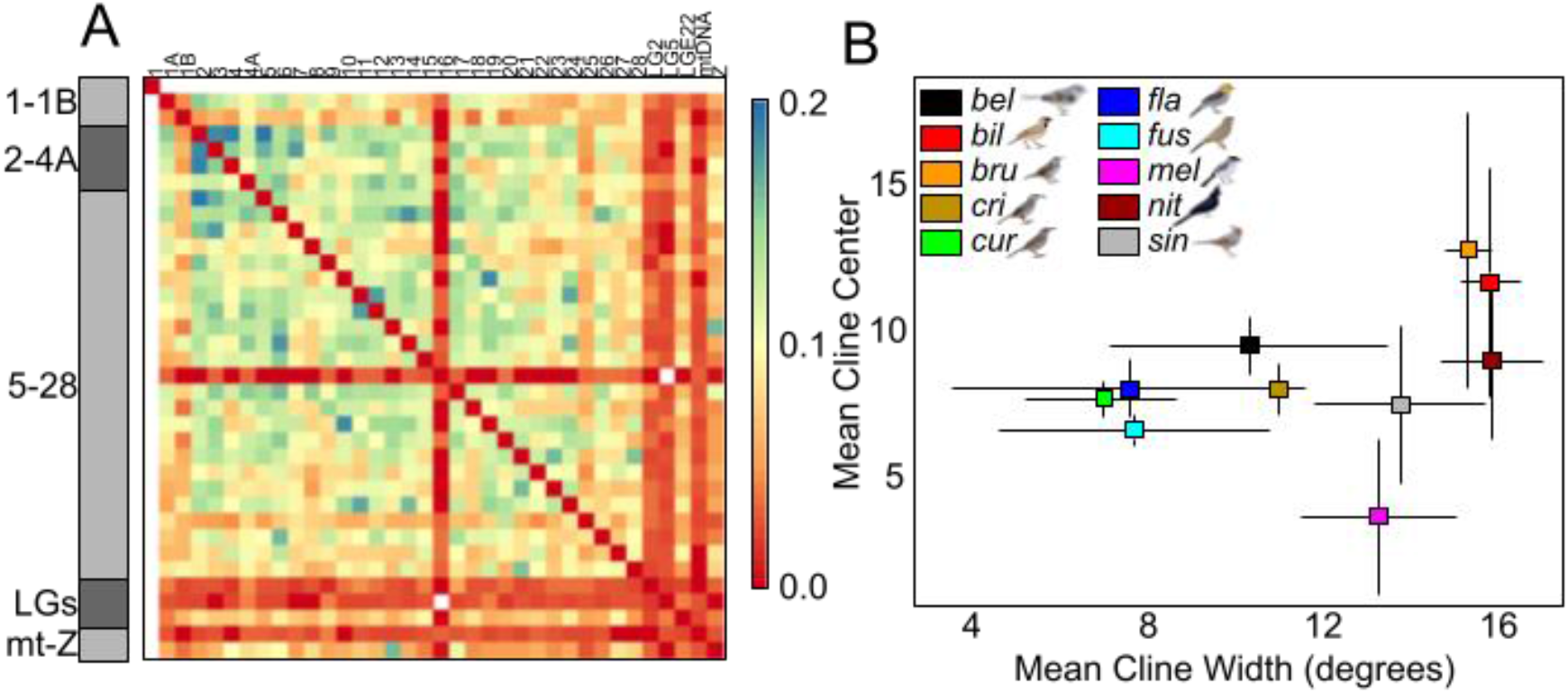
Species varied both in genetic structure across chromosomes and population structure across the Sonoran and Chihuahuan Deserts. A) Standard deviations of normalized Robinson-Foulds distances, averaged across species; see Supplementary Figure 4. Warmer colors indicate lower standard deviations and less variation across taxa. Chromosomes are arranged in alphanumeric order; y-axis shows blocks of chromosomes (gray) for legibility, x-axis shows individual chromosomes. B) Mean cline width in degrees vs. mean cline center across chromosomes for each species; see Figure 3. Lines from each point show standard deviations. Species names are shortened for legibility (“bel”=*Vireo bellii*, “bil”=*Amphispiza bilineata*, “bru”=*Campylorhynchus brunneicapillus*, “cri”=*Toxostoma crissale*, “cur”=*Toxostoma curvirostre*, “fla”=*Auriparus flaviceps*, “fus”=*Melozone fusca*, “mel”=*Polioptila melanura*, “nit”=*Phainopepla nitens*, “sin”=*Cardinalis sinuatus*). Colors indicate individual species.

Species varied in how wide their clines of genetic relatedness were between chromosomes. Mean cline width ranged from 6.9–15.9° longitude, where the total area encompassed by each species was ∼18° longitude (with zero on the cline defined as 116.1°W longitude; Supplementary Table 2; Figure 3; Figure 4; Supplementary Figure 1). Cline width increases as chromosome size decreases (p=1.4×10^−6^, adjusted R^2^=0.06), though this varies across species (range p=7.7×10^−7^– 0.43, range adjusted R^2^=-0.01–0.51). Mean cline center location ranges from 3.6° along the cline (∼112°W) to 12.7° along the cline (∼103°W). We found that there were negative correlations between the degree of population structure (measured by F_ST_; see Supplementary Information) and both mean cline width and the standard deviation of cline center locations, which was expected based on how clines are calculated. Species with higher F_ST_ between populations had narrower clines and less variation among partitions in the locations of their clines (Supplementary Figure 5). Cline width is also significantly, but weakly, associated with recombination rate (p=0.0023, adjusted R^2^=0.02)

### Phenotypic variation across the Cochise Filter Barrier

There were no clear, desert-specific patterns in morphological variation across the Cochise Filter Barrier (N=234), with morphological changes ranging from subtle to significantly different. In our principal components analysis, the first three principal components (PC1, PC2, PC3) explained 74%, 12%, and 6% of the variation in morphology and corresponded approximately to overall body size, bill size/shape, and wing size/shape, respectively (Supplementary Table 3, Supplementary Table 4; Supplementary Figure 9). We found significant differences across the Cochise Filter Barrier in six species in at least one analysis (Figure 6; see Supplementary Information for more details). Between deserts, *T. crissale* and *C. sinuatus* differed in body size and bill shape. *Vireo bellii* and *M. fusca* differed in bill shape. *Polioptila melanura* and *A. flaviceps* differed in body size. No species showed significant differences in wing shape.

### Climatic suitability and abundance across the Cochise Filter Barrier

During the Last Glacial Maximum, the most suitable areas for all taxa were projected to be further south than the most suitable areas during the present and mid-Holocene. Regions that are predicted to be suitable through all three periods are often reduced compared to current distributions (Supplementary Figure 8; Supplementary Figure 10). We calculated abundance for each species using the Breeding Bird Survey (Pardieck et al., 2019). Abundance was correlated with predicted climatic suitability across all taxa, with adjusted R^2^ values of fit lines (log-scaled) ranging from 0.42–0.62 (Figure 4, Supplementary Figure 6, Supplementary Figure 7).

### Phenotypic and genotypic datasets are idiosyncratic with respect to landscape features

We used generalized dissimilarity matrix (GDM) models to determine which geographic features best described variation in different partitions of genetic and phenotypic data. We had 515 combinations of species and partitions (out of a total possible of 540). For univariate models, performance of generalized dissimilarity matrix models was generally consistent whether looking at univariate, bivariate, or trivariate data partitions (see Supplementary Information). 2,899/3,090 univariate models converged successfully with an overall 94% convergence. Of those 515 datasets tested, 18.0% selected IBE as the best factor explaining variation, 17.5% selected IBB, 17.2% selected IBA, 9.1% selected IBD, 18.8% selected IBH, and the remainder were ambiguous, with multiple models equally explaining variation. Within the ambiguous models, of which there were 98, the best models often included IBE (99.0% of models), IBH (81.6%), and IBD (72.5%); in contrast, the best models rarely included IBA (4.1%) or IBB (2.0%). Across all the GDMs, percent deviance explained by the best model was variable, ranging from 0.1% to 81.9%. The mean±SD percent deviance explained for these runs was 12.7%±13.6%. Percent deviance explained for the whole genome was lower on average, ranging from 0.1%–29.2% (mean±SD 10.8%±10.4%). F_ST_ outliers, both high and low, tended to have lower percent deviances explained, ranging from 0.1%– 21.9% (mean±SD 6.5%±6.5%). Lostruct outliers ranged from 0.5%–32.2% (mean±SD 8.1%±7.3%). Percent deviance explained had the most extreme range in morphology, from 0.3% to 81.9% (mean±SD 16.6%±20.8%). The percent deviance explained for all datasets varied across taxa, with means ranging from 3.2% (*M. fusca*) to 20.3% (*A. bilineata*) and standard deviations ranging from 8.7%–16.4%.

For the models examining signals across the whole genomes, three species had IBB as the most important predictor, one had IBE, two had IBH, one had IBA, and three had mixed support. (Figure 5; Supplementary Figure 11). IBD was the least common predictor across chromosomes (5.2%), while all other predictors were of approximately equal frequency (19.6% IBH, 19.0% IBE, 18.2% IBB, 17.0% IBA, and 20.8% mixed support for multiple models). Within the mixed models, IBE was included 100% of the time, IBH was included 77.7% of the time, IBD was included 73.6% of the time, and IBE and IBB were each included 2.3% of the time.

**Figure 5:**
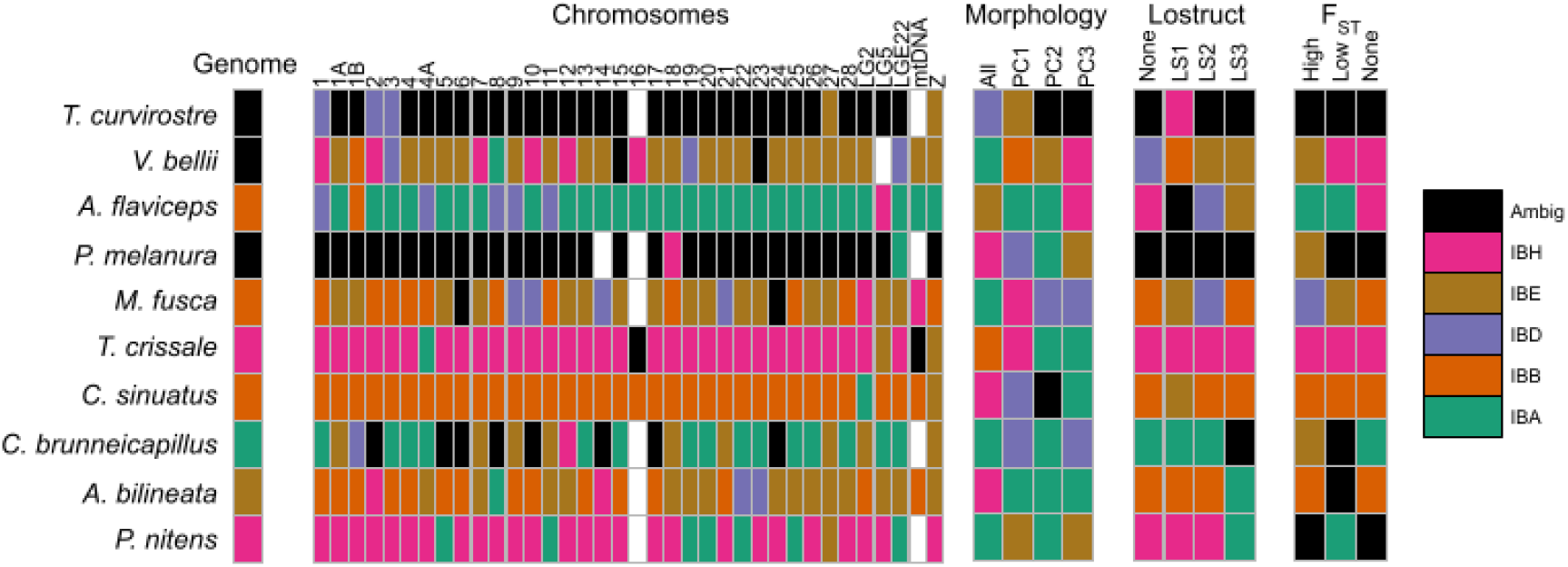
Generalized Dissimilarity Modeling revealed heterogeneous associations between genomic and phenotypic differentiation and alternative geographic hypothesis. Shown are the best performing GDM across all univariate, bivariate, and trivariate models. Species are along the y-axis and arranged from most to least differentiated across the Cochise Filter Barrier. Individual partitions are along the x-axis (whole genome, individual chromosomes, morphology, lostruct partitions, F_ST_ outliers). “Genome” refers to a partition where all genomic information was assessed at once. Color indicates the best model. The alternative models were as follows: isolation by abundance (IBA), isolation by barrier (IBB), isolation by distance (IBD), isolation by environment (IBE), and isolation by history (IBH). “Ambig” is shorthand for ambiguous partitions where multiple models equally best explain the data. White boxes represent models that failed to converge or did not have corresponding datasets. For more partitions of data see Supplementary Figure 2 and Supplementary Figure 11.

**Figure 6:**
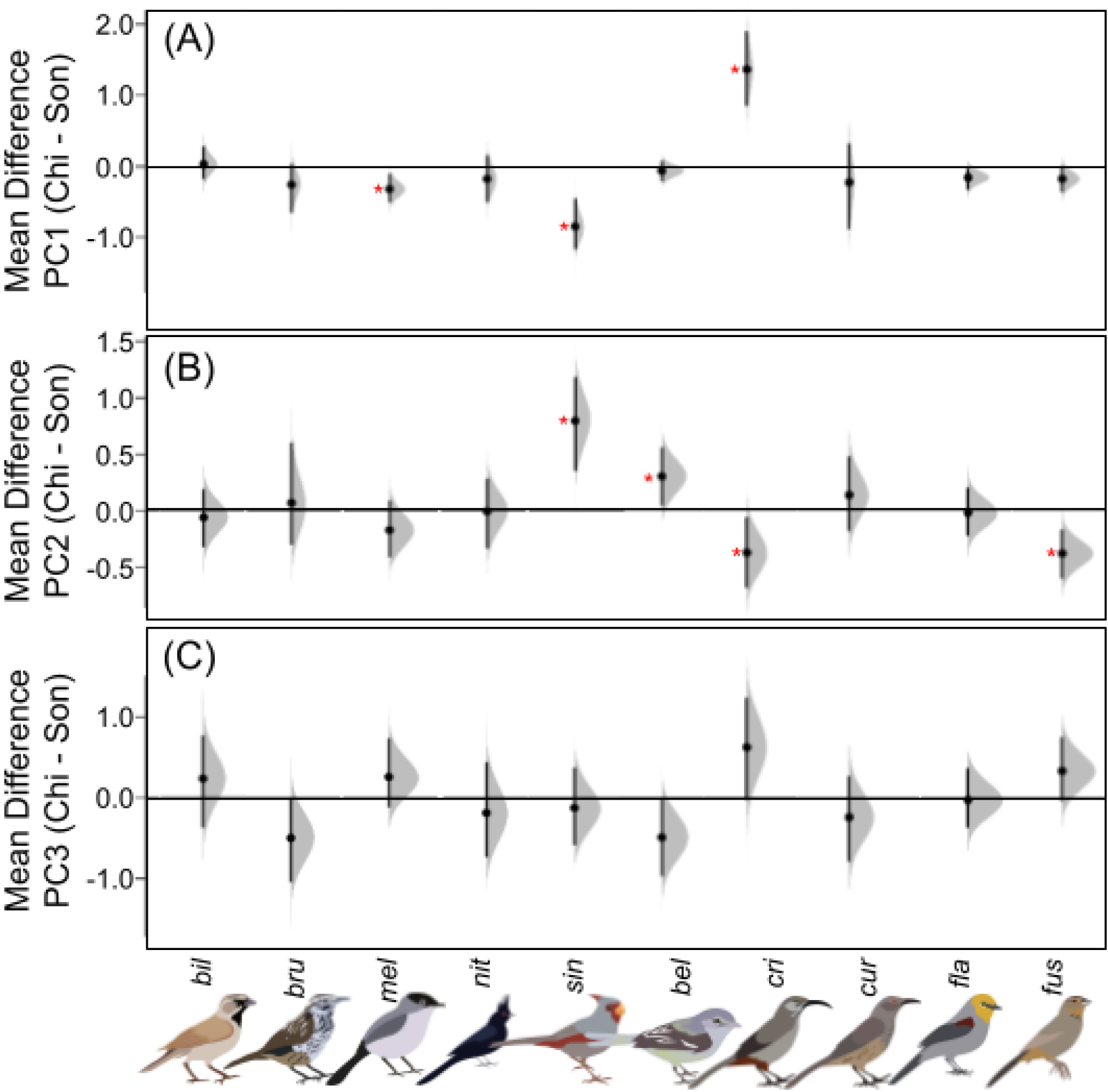
Distribution of unpaired mean differences between Sonoran and Chihuahuan Desert individuals for each species from DABEST analysis for morphological PC1 (A), PC2 (B), and PC3 (C). Black horizontal line is at zero, black points and vertical lines show mean and confidence intervals for each distribution in gray. Comparisons that do not cross zero are considered significant in DABEST tests, indicated with red asterisk. On the X axis are each species with images (scale does not reflect size differences) with species names are shortened for legibility (“bel”=*Vireo bellii*, “bil”=*Amphispiza bilineata*, “bru”=*Campylorhynchus brunneicapillus*, “cri”=*Toxostoma crissale*, “cur”=*Toxostoma curvirostre*, “fla”=*Auriparus flaviceps*, “fus”=*Melozone fusca*, “mel”=*Polioptila melanura*, “nit”=*Phainopepla nitens*, “sin”=*Cardinalis sinuatus*).

For the lostruct partitions, the outlier partitions (LS1, LS2, LS3) had 4/30 with IBA as the best model, 6/30 IBB, 2/30 IBD, 5/30 IBE, 6/30 IBH, and 7/30 as ambiguous. Among the ambiguous models, all of them showed IBE as important and nearly all showed IBH, IBD, or both as important Most species showed at least some overlap in which model best explained partitions: for example, *A. bilineata* and *C. sinuatus* all have at least two lostruct partitions best explained by IBB.

For the non-outlier partitions (LS0), the best model chosen was the same as the best model explaining whole-genome variation in all but three species (*V. bellii, A. flaviceps, and A. bilineata*) and that of one of the outlier partitions in all but two species (*V. bellii, A. flaviceps*). Notably, for *P. melanura* all three outlier partitions, the genome, and the non-outlier lostruct partitions are explained by multiple models (specifically, IBD, IBE, and IBH for all). Likewise, for *T. crissale*, all of these were explained by IBH.

For the genomic regions with F_ST_ outliers, the best predictors across species were generally congruent between different outlier partitions and the whole genome. In all species but *A. bilineata*, the non-F_ST_-outliers had the exact same best predictors as that of the whole genome (or in cases where multiple models were equally good predictors, one was a subset of the other). High-F_ST_ outliers showed different best predictors than the genome in *C. brunneicapillus, A. bilineata, A. flaviceps*, and *M. fusca*. Low-F_ST_ outliers showed different best predictors than the genome in *C. brunneicapillus, A. flaviceps, M. fusca*, and *P. nitens*.

There was little congruence across the best landscape predictor of morphological data within species; however, the best performing model across these three datasets was most frequently IBA (37.5%), IBD (17.5%), and IBH (17.5%), with relatively fewer models with IBE (12.5%), IBB (7.5%), IBB or a mixture of models (7.5%, with approximately even amounts of IBA, IBD, IBE, and IBH making up the mixture). 3/30 of the PCs matched overall morphology in terms of best predictors (including mixtures of models). Additionally, 10/30 individual PCs did match each other when they did not match the genome: PC1 and PC2 in four species, PC1 and PC3 in two species, and PC2 and PC3 in four species. Notably, all PCs in *A. bilineata* were best explained by IBA despite its overall morphology being best explained by IBH. While the distribution of best models for overall morphology, PC1, and PC3 were not significantly different than expected, for PC2 this was nearly significant (χ^2^=6.8, p=0.079, df=3, simulated p=0.11)

Overall morphological variation was best explained by IBA in 4/10 species, IBH in 3/10, and 1/10 each for IBB, IBD, and IBE. In contrast, PC1 (body size) showed a more even distribution between all models (1/10 IBE, 2/10 IBA, 3/10 IBD, 2/10 IBB, 2/10 IBH). PC2 (bill shape) was best explained in 6/10 of species by IBA, 1/10 each by IBE and IBD, and 2/10 with a mixture of results (combinations of IBA, IBD, IBH, and IBE). Lastly, PC3 (wing shape) was best explained in 3/10 of species by IBA, 2/10 each by IBE, IBD, and IBH, and 1/10 of species had ambiguous results (IBA, IBE, and IBH).

### Data characteristics of best-fit models

Genomic summary statistics were associated with which geographic patterns best predicted variation within species. Cline width per chromosome was significantly different relative to the predictors (p = 1.85e-5), being wider between IBB models and IBD or mixed models, between IBH models and IBD or mixed models, and between IBE and IBD models. Cline centers also significantly differed, with chromosomes supporting mixed models having much more eastern cline centers than chromosomes supporting IBA, IBB, IBE, or IBH models. Centers were also significantly more eastern for chromosomes predicted by IBA models than by IBH models (p=8.86e-10). Chromosomes with lower recombination were significantly more likely to be explained by mixed models than by IBA or IBE models (p = 0.0147). Chromosomes explained by mixed models also had higher estimated F_ST_ than those explained by IBA, IBB, or IBH models (p = 4.2×10^−5^). Chromosomes with IBH as the best model had lower D_XY_ than those with IBB or IBE as best models. Chromosomes with less missing data were more likely to show mixed support for models compared to IBA, IBE, or IBH models, and more likely to show IBB over IBA or IBE models. Species with higher mean contact zone suitability were more likely to have IBB as the best model compared to all other models, and species with lower contact zone suitability were more likely to have IBH as the best model compared to all other models. Likewise, species with highly variable habitat suitability were more likely to have IBH as the best model. Not significant at all was chromosome length across predictors. Tajima’s D was significantly different across chromosomes with different models (p = 0.0432), but Tukey’s honestly significant difference tests showed that none of the individual comparisons were significant.

Species differed more than expected with respect to what geographic models best explain their genotypes and phenotypes. Best-predictors vary across individual species (χ^2^=816.8, p∼0.0, df=45, simulated p<0.0005) and with respect to whether or not species have phylogeographic structure across the Cochise Filter Barrier (χ^2^=188.6, p∼0.0, df=10, simulated p<0.0005). However, best-predictors did not vary with respect to individual genotypic or phenotypic partitions (χ^2^=238.3, p=0.88, df=265, simulated p=0.88).

## Discussion

We tested modes of population structuring in birds distributed across a biogeographic filter barrier, where we found that genomic landscapes were best-explained by different geographic models across partitions at multiple scales. The disparity in predictors of intraspecific differentiation among the whole genome versus windows and between windows extends the view that evolutionary inferences are dependent on which portions of the genome are examined in a spatial framework. Despite this, individual species behave more consistently than expected across all of their corresponding genomic and phenotypic partitions. The heterogeneity in model fit between taxa partitions was consistent with the expectation that various evolutionary processes contribute to the peaks and valleys of the genomic landscape. By applying this framework across an assemblage of birds that evolved across a common, dynamic region we showed that at the community-scale, predictors of genomic structure remain idiosyncratic across the community, which may reflect taxa at different stages of the population histories and responses to a barrier that mediates gene flow.

### Extrinsic drivers of the genomic landscape

Our modeling showed that environmental distance was a common predictor of levels of intraspecific differentiation, but this pattern was species-dependent. Contemporary environment was the single most important or one of the most important factors in nearly 40% of partitions, followed closely by the paleoclimate environment (Supplementary Figure 2). Genome-wide patterns of differentiation across the Cochise Filter Barrier are partially shaped by environmental adaptation as observed in non-avian taxa distributed across the barrier (Myers et al., 2019). Environmental adaptation is often recovered in taxa who respond to environmental gradients via altered phenotypes (Branch et al., 2017, Dubec-Messier et al., 2018), genotypes (Berg et al., 2015,

Manthey and Moyle, 2015), or both (Ribeiro et al., 2019). Despite the importance of environment on the genotype and phenotype in these birds, the fact that patterns are highly species specific instead suggests that individual taxa have unique responses to those environments. Although the focal taxa are co-distributed, we showed how environmental suitability, their general morphologies, and abundances across space varied among species, which may help explain why best-fit models differed. As such, these species-specific factors likely explain isolation by environment was the best explanatory variable for many, but not all, of the species we investigated. Individual partitions of the genome also varied with respect to how much environmental variation played a role. At one extreme, environmental variation appears to have a strong impact on the sex chromosomes. Environment was the most (or one of the most) important factor on the Z chromosome for 6/10 species, including species with population structure, a gradient, and panmixia. This is likely because the chromosome evolves faster than sites under selection for adaptation to local environmental conditions. Sex chromosomes are known to diverge faster than autosomes due to their differences in effective population size (Mank et al., 2010), importance in sexual selection (Kirkpatrick, 2017), and the presence of speciation genes (Sæther et al., 2007). Given the evidence for environmental variation predicting genetic differentiation on the Z chromosome, this would suggest that any speciation genes present in these taxa may be involved in adaptation to the environment. The autosomes, in contrast to the sex chromosomes, vary in how important environment is, from some chromosomes with environment only being one of multiple factors (i.e., chromosome 1) to autosomes that are majority driven by environment (i.e., chromosome 27).

The environment was the most important driver for species with genetic structure, with 35.3% of partitions in structured species having the environment as the best model. The most intuitive explanation for this was that population structuring in these taxa was facilitated by natural selection to the environmental gradient across the barrier. There was some evidence that this could have happened across other taxa that occur across the Cochise Filter Barrier, as IBE was the best predictor of genome-wide divergence in a community of snakes distributed across the barrier (Myers et al., 2019). However, we must stress that while this explanation was the most intuitive and aligns with predictions, there are numerous processes that can produce IBE (Wang and Bradburd, 2014), and it is possible that divergence led to adaptation to these environments secondarily, rather than the reverse, or the patterns are being influenced by some factors that we did not quantify. Nevertheless, at present our results are consistent with the importance of ecologically mediated population differentiation, or isolation by environment, in structuring communities across the deserts of North America.

### Contemporary versus historical predictors of genomic differentiation

Our finding that the best-fit models varied across species was consistent with the expectations that species idiosyncratically respond, over a range of time scales, to the Cochise Filter Barrier. The spatial patterns we examined vary temporally, with Pleistocene environmental changes being a historical process, while geographic distances, abundances, and environmental variation reflecting more contemporary processes. Historical signatures of Pleistocene isolation are commonly recovered patterns for the Cochise Filter Barrier (Provost et al., 2021) and other communities (Shafer et al., 2010; Ralson et al., 2021), and our data showed that isolation in glacial refugia best explained genome-wide differentiation in two of our species, one that showed a gradient of phylogeographic relatedness and one that was unstructured. Within chromosomes, there are two additional species where one of multiple, equally-well-fit predictors is historical isolation. The lack of signal in the other six species, particularly the ones with phylogeographic structure across the barrier, could be due to erosion of historical signals as the Cochise Filter Barrier filters taxa and changes the contemporary patterns of gene flow. Alternatively, our proxy for IBH (resistance over projected Pleistocene habitat suitability) may be a poor model for actual historical isolation. For example, paleoenvironmental gradients may no longer be as readily detectable. Nevertheless, this lack of support for paleoenvironmental factors, and thus glacial refugia, suggests that these processes may not leave strong detectable signals in the genomes of most of these desert birds.

In contrast, current environments best explain a large amount of genetic and morphological variation, suggesting that phenomena operating on more recent timescales influenced contemporary patterns across the landscape. If some of the taxa herein are going through incipient speciation, then these contemporary factors should be most potent. Our identification of species abundances as a relatively important predictor of genetic divergence aligns well with landscape genetic studies that use proxies for the effects of contemporary phenomenon and ecological factors on genetic variation (Burney and Brumfield, 2009; Paz et al., 2015). For example, urbanization, which fragments and reduces population sizes, is well known to impact rates of gene flow and drift, acting as a strong barrier of gene flow since the 20th century (Miles et al., 2019). Our use of available abundance data across large spatial scales shows a more direct relationship between varying abundances across the landscape with levels of differentiation. Further, while both historical and contemporary processes are influencing taxa across this biogeographic barrier, environmental patterns in particular irrespective of timing seem more influential.

### Relationship between best-models and window summary-stats

In contrast to the extrinsic drivers of the genomic landscape that we have focused on here, there were few clear associations between partition characteristics and support for a particular model. For example, we found that regions with low predicted recombination rate were more likely to show multiple models as equally important. At the phylogeographic-scale, low recombination regions of the genome have been shown to be more likely to reflect population structure (Martin et al., 2019, Li et al., 2019, Manthey et al., 2021). The avian recombination rate landscape is thought to be conserved across taxa, even though exact genomic locations of divergence across taxa are not (Singhal et al., 2015, Turbek et al., 2021). Correlations in recombination rates at the same genomic position in other species are greater than 0.37 across chromosomes and always positive (Turbek et al., 2021), even across large phylogenetic distances. The ten desert birds we investigated, which range in divergence time from ∼9 to ∼55 million years between taxa (Harris et al., 2018; Kumar et al., 2017; Barker et al., 2015; Mason and Burns, 2013; Price et al., 2014; Pasquet et al., 2014; Hooper and Price, 2017; Mitchell et al., 2016; Gibb et al., 2015), have correlations in recombination rates at the same genomic position that were often smaller in magnitude and negative. This could reflect a real pattern, where the recombination landscapes are only conserved within more closely related species; our closest taxa, the two non-sister *Toxostoma*, do have the highest correlation in recombination rates across windows and are in the top 25% of the distribution in correlations. However, the differences found could have been caused by coverage depth, differences in the recombination rate estimators used, missing data allowance, or the fact that software that estimates recombination rates do not currently exist that can handle genotype likelihood data. In addition, genetic partitions with higher F_ST_ were more likely to show multiple best models as being important. We expect regions with high differentiation to instead be associated with the presence of the barrier if the barrier reflects actual divergence. However, this was not the case. We suggest that this reflects the gradient in differentiation across species in the community, both in the degree to which divergence has happened, the genomic locations of any differentiation, and the timing of divergence.

We explored the signal in our data by examining multiple ways of partitioning genomic windows, using different thresholds of missing data, and evaluating how data attributes influenced model support. We found that genetic partitions with more missing data were more likely to have ambiguous results. Genetic summary methods like PCA are impacted by missing data, particularly when they are imputed, which can cause individuals with disproportionately high levels of missing data to appear like they are admixed between populations (Yi and Latch, 2021). It is likely that the reverse is true, where individuals with disproportionately low levels of missing data should fall out as their own populations more readily. Here we expect individuals with exceptionally low coverage should behave similarly. For example, in some of our species (namely *Vireo bellii, Auriparus flaviceps, Polioptila melanura*) the individuals with highest missing data clustered as their own population before detecting any other spatial patterning. We ameliorated this by dropping individuals with too much missing data in some of our datasets. Overall, we did not find qualitative differences in population assignments, but it did generally inflate our fixation values and deflate our genetic diversity values. This is sensible, as reducing the number of individuals should both increase the likelihood of fixation due to sampling error as well as decrease the overall amount of nucleotide diversity.

The clines of population differentiation across space that we measured were narrower in longer chromosomes. One explanation for this is that larger chromosomes are more dense with respect to polymorphisms across the deserts (Supplementary Figure 24), therefore having more information content with respect to clines. However, we propose that this is mediated by recombination variation across the genome. Chromosome length is frequently negatively correlated with recombination rate, where generally, the recombination rates are lower on larger chromosomes due to the necessity of crossovers to ensure successful meiosis (Tigano et al., 2022). This is a common occurrence in many taxonomic groups (Kaback et al., 1992; Jensen-Seaman et al., 2004; Pessia et al., 2012; Farre et al., 2013, Kawakami et al., 2014, Haenel et al., 2018 Tigano et al., 2022). Lowered recombination rate would be less likely to break up genetic variants within the genome in the event of gene flow between two populations. Further, SNP diversity is positively correlated with recombination, possibly due to mutagenesis at those sites (Lercher and Hurst, 2002; Arbeithuber et al., 2015) Regions of low recombination are known to facilitate genomic changes such as selective sweeps (e.g., Burri et al., 2015; Bourgeois et al., 2019). However, in our dataset recombination rate was not associated with the size of the chromosome. Post-hoc, we broke down this relationship into structured and unstructured species, where we found that species with structure or a gradient showed no association, while species that were panmictic exhibited the assumed negative relationship. Our within-species recombination estimating method is known to be sensitive to historical demographic events (Adrion et al., 2020); as such, the presence of population structure herein may have caused the estimates to deviate from expected patterns. As such, we suspect that recombination landscape differences associated with chromosome length are contributing to the differences in these clinal patterns.

### Morphological versus genetic associations

We found that in most taxa, genotypic and phenotypic variation within species, and even different aspects of morphological phenotype within species, were not associated with the same landscape factors, in contrast to high congruence within species in different genotypic datasets. Phenotypes were better explained by abundance, whereas genotypes were better explained by the contemporary and historical environment. Discordance between genetic and phenotypic predictors of spatial variation have been observed in other systems, where phenotypic variation was better explained by the environment (Moreira et al., 2020). This discordance could be due to polygenic traits, where genotype-phenotype associations may be mediated by multiple loci of small effect working in concert, either by changing protein structure or regulation (Yusuf et al., 2020, Knief et al., 2017, Duntsch et al., 2020, Aguillon et al., 2021). However, for some phenotypes like plumage color, single genes of large effect have been implicated which should strengthen correlations between genotype and phenotype, at least for those loci (Sin et al., 2020; Toews et al., 2016). For desert birds in particular, phenotypic variation in metabolism (as well as in microbiomes) has been linked to genes that vary with the environment (Ribeiro et al., 2019). In our study, as with genetic differentiation, the extent of phenotypic structuring varied across species, with bill and body size being significantly different between deserts in a few taxa, but somewhat surprisingly, environmental variation did not usually explain morphological differences. For example, adaptations in bill morphology are frequently observed, such as in Song Sparrows on the Channel Islands that have higher bill surface area in hotter climates (Gamboa et al., 2021). The lack of a tight correlation between environment and phenotype in our study were likely reflective of the shallowness of the evolutionary divergences and the subtlety of the environmental gradient across deserts. The two *Toxostoma* species in our study have previously shown contrasting patterns with respect to climate on beak morphology: *T. crissale* has larger bills in drier habitats, which may aid in cooling while conserving water, while *T. curvirostre* showed a pattern contrary to thermoregulatory predictions with larger bills in cooler climates (Probst et al., 2021), suggesting even in closely related species climate may not have the same role on morphological variation. Even though phenotypic data partitions often did not have the same explanatory factor with respect to the general dissimilarity modeling, there was a correlation between population structure in the genome (and chromosomes to a lesser extent) and phenotypic variation across these ten birds, in that taxa lacking morphological change also lacked genetic variation overall.

### Fitness effects of the Cochise Filter Barrier

We found multiple species that have relatively sharp clines across the Cochise Filter Barrier, typically the taxa that also show population structure. These clines may represent areas that are hybrid zones, potentially under selection against the two populations coming back into contact. Our sampling throughout that transition zone is quite extensive, with the exception of *V. bellii*. In three species (*T. crissale, T. curvirostre, M. fusca*) there are one or two individuals close to the transition zone between the deserts that have intermediate assignments between populations according to our NGSadmix analysis. For *T. curvirostre* in particular, this is close to where hybrid individuals have already been suggested to exist (e.g., Zink and Blackwell-Rago, 2000). Further, one species (*P. melanura*) has individuals close to this transition zone, though only when three populations are assigned rather than two. Multiple individuals of two species (*A. bilineata, C. sinuatus*) also come out as being admixed, but distributed throughout the range of the species. It is likely that the Cochise Filter Barrier is thus causing fitness effects, especially in those taxa that have few individuals admixed in the transition zone. Further investigation with more explicit determination of hybrid status in these species is likely warranted.

### Conclusion

By quantifying patterns in genotypic and phenotypic variation in communities distributed across a barrier to gene flow, we found that multiple co-occurring processes occur that impact genomic and phenotypic divergence within taxa. Environmental gradients were among the most important associations in predicting genetic and phenotypic variation, but the best-fit model was highly associated with species-specific patterns. These findings underscore the importance of accounting for heterogeneity in the genome, phenome, and diversification mechanisms acting across time and space to have the most comprehensive picture of geographic structuring in species. This will allow for an assessment of whether biotic and abiotic geographic variation, which act as proxies for neutral and adaptive processes, consistently predict variation across phenotypes and genotypes that are evolving under the same conditions. Without a holistic understanding at each of these levels of organization, as well as the addition of future work that concurrently estimates selection at the organismal and the nucleotide levels, the actual mechanisms that shape communities will remain obscured. Overall, this work displays the necessity of integrating geographic predictors of population divergence, differentiation across the genomic landscape, and phenotypic variation in understanding the multiple different mechanisms that have produced the population histories we see across contemporary communities of birds in North America.

## Methods and Materials

### Study system

The Sonoran and Chihuahuan Deserts contain environmental and landscape variation that make them suitable for testing if any of the five discussed geographic models (IBA, IBB, IBD, IBE, and IBH) structure intraspecific variation in taxa. Across the two deserts and the transition zone between them, there is variation in precipitation, elevation, temperature, and vegetation that could result in local adaptation and isolation by environment. (Shreve, 1942; Reynolds et al., 2004). Pleistocene glacial cycles repeatedly separated and connected, such that some taxa experienced dramatic range shifts (Smith et al., 2011; Zink, 2014), which could have isolated taxa in each desert. Further, there is a well-studied biogeographic barrier separating the deserts, the Cochise Filter Barrier, which is an environmental disjunction that demarcates the transition between the Sonoran and Chihuahuan Deserts of southwestern USA and northern Mexico. The barrier is thought to have begun forming during the Oligo-Miocene and completed during the Plio-Pleistocene (Morafka, 1977, Van Devender, 1990; Van Devender et al., 1984, Holmgren et al., 2007, Spencer, 1996) and has formed a community ranging from highly differentiated taxa to unstructured populations (Provost et al., 2021). Demographic troughs caused by geographically varying population abundances could impact the frequency of gene flow across the landscape and the degree of genetic connectivity across the deserts.

### Genetic sequencing and genome processing

We performed whole-genome-resequencing for 10 species of birds from the Sonoran and Chihuahuan Deserts, obtaining genetic samples from new expeditions and loans from natural history museums (*Cardinalis sinuatus*; *Toxostoma crissale, Toxostoma curvirostre*; *Amphispiza bilineata, Melozone fusca*; *Polioptila melanura; Phainopepla nitens*; *Auriparus flaviceps*; *Campylorhynchus brunneicapillus*; *Vireo bellii*; Supplementary Table 5; Supplementary Figure 15). These species reflect different songbird morphotypes and ecologies in the deserts (e.g., large-to small-bodied, insectivorous to granivorous, migratory to resident). Three of these species (*V. bellii, T. curvirostre, M. fusca*) have shown evidence of structure across the Cochise Filter Barrier, while an additional three (*P. melanura, A. flaviceps, C. brunneicapillus*) have shown evidence of no structure (Zink et al., 2001; Rojas-Soto et al., 2007; Teutimez, 2012; Klicka et al., 2016, Smith et al., 2018). However, some of the taxa without structure at the Cochise Filter Barrier do have structure at other barriers (e.g., Vázquez-Miranda et al., 2022).

Using 221 individuals across our 10 focal species, we sequenced 8–14 individuals in both the Sonoran and Chihuahuan Deserts per species for a total of 18–25 samples per species. We extracted DNA using the MagAttract HMW DNA Kit (Qiagen); 33 of the samples were extracted using a Phenol-Chloroform protocol, but we switched to the former to improve extraction quality. Library preparation and sequencing was performed by RAPiD Genomics (Gainesville, FL) on an Illumina HiSeq X PE150. All individuals sent on the same plate were sequenced across N lanes, where N is the number of samples divided by 20. We sent six plates which ranged from 20–96 individuals (some plates also contained individuals from other projects).

We mapped raw reads of each species to their phylogenetic closest available reference genomes (Supplementary Table 6): notably, *A. bilineata* and *M. fusca* were mapped to the same genome, as were *C. brunneicapillus, T. crissale, T. curvirostre, P. melanura*, and *P. nitens* (see Supplementary Information). Before mapping, we created pseudo-chromosomal assemblies of these genomes using Satsuma version 3.1.0 (Grabherr et al., 2010) by aligning to the *Taeniopygia guttata* genome (GCF_000151805.1), retaining pseudo-chromosomes with the prefix “PseudoNC”. Hereafter, pseudo-chromosomes will be referred to as chromosomes.

We filtered our sequences with FastQ Screen version 0.14.0 (Wingett et al., 2018) to remove contamination by filtering out reads that mapped to PhiX and the following genomes: *Homo sapiens, Escherichia coli, Enterobacteriophage lambda*, and *Rhodobacter sphaeroides*. For more details on bioinformatics methods, see Supplementary Information. In brief, we did the following: From our raw reads, we used a pipeline that produced genotype likelihoods using ANGSD version 0.929 (Korneliussen et al., 2014). We converted cleaned FastQ files to BAM using bwa version 0.7.15 (Li and Durbin, 2009, Li and Durbin, 2010) and picard version 2.18.7-SNAPSHOT from the GATK pipeline (McKenna et al., 2010, DePristo et al., 2011, Van der Auwera et al., 2013). Next, we prepared the BAM files to be used in the ANGSD pipeline using samtools version 1.9-37 (Li et al., 2009; Li, 2011), bamUtil version 1.0.14 (Jun et al., 2015), and GATK version 3.8-1-0 (McKenna et al., 2010). This methodology creates genotype likelihoods to account for uncertainty for low-coverage sequences.

We investigated the impact of missing data (due to low coverage) on our analyses using three thresholds for retaining sites: a complete dataset, in which all individuals were retained irrespective of missing data; a 75% dataset, in which individuals were only retained if they had less than 75% missing sites; and a 50% dataset, in which individuals were only retained if they had less than 50% missing sites. These different datasets were used for a suite of downstream analyses to assess the sensitivity of the results to individuals with missing data.

### Evaluating population structure across the Cochise Filter Barrier

We characterized the degree of population structure across the whole genome and in individual chromosomes across the Cochise Filter Barrier in our focal species. First, we used a combination of PCAngsd in ANGSD (Meisner and Albrechtsen, 2018) and NGSadmix (Skotte et al., 2013), to assign individuals to K clusters and estimate admixture proportions for each individual. We chose K=2 to evaluate whether there was structure across the Cochise Filter Barrier (though we visualized K values from two to three). Because of differences in coverage among individuals, we performed this for the complete, 75%, and 50% missing data datasets, but found that these values were largely congruent across the datasets, and so we only use the complete dataset for describing population structure (Supplementary Figure 16, Supplementary Figure 17, Supplementary Figure 18). Second, we plotted PCAngsd individual population assignments over space using a cline analysis via the hzar version 0.2-5 R package (Derryberry et al., 2014) and custom scripts (modified from Burbrink et al., 2021). Analyses were conducted in R version 3.6.1 (R Core Team, 2019). We did this to quantitatively evaluate the differences in population structure across chromosomes and in the genome more broadly. We thus were able to calculate the location and width of clines for the entire genome and each chromosome.

Complementing our genome-wide analyses, we ran a local principal components analysis along the genome on the complete dataset using the R package lostruct version 0.0.0.9000 (Li and Ralph 2019). Different chromosomes showed different relationships between individuals with respect to predicted phylogeographic relatedness (see Supplementary Information). Because of this, we wanted to cluster regions of the genome together that showed similar relationships between individuals in case specific evolutionary processes were causing this pattern. The lostruct method performs principal component analysis on individual windows of the genome, then uses multidimensional scaling (MSDS) to summarize how similar the windows’ principal component analyses are when dividing the genome. To accommodate genotype likelihoods in the method, we calculated covariance matrices using PCAngsd to describe the relationships between individuals, then fed those covariance matrices into the lostruct code. We extracted three subsets of outliers for each species, which we designated LS1, LS2, and LS3, and compared it to the remainder of the genome, representing non-outliers.

### Genomic summary statistics

We characterized genetic variation across each species’ genome and partitions of the genome by calculating a suite of summary statistics and metrics. To quantify genetic differentiation within each species, we calculated pairwise genetic distances from the genotype likelihoods using NGSdist (Vieira et al., 2016), which served as the genetic distance matrices for our generalized dissimilarity matrix models (see below). Neighbor-joining trees were calculated from these matrices to contrast genealogies across the genome. Genealogies across the genome were visualized by calculating pairwise and normalized Robinson-Foulds (RF) distances between all pairs of trees per species (Robinson and Foulds, 1981). We also performed a sliding window D_XY_ analysis using the calcDxy R script included with ngsTools version 1.0.2 (Fumagalli et al., 2014), which gives site-wise D_XY_ values, and then averaged across windows. Windows were overlapping with a size of 100,000 base pairs and offset by 10,000 base pairs. Missing data were calculated using vcftools (Danecek et al., 2011). This was calculated per window, per chromosome, per genome, per site, and per individual.

Using ANGSD’s realSFS function, we performed a sliding window F_ST_ analysis by converting SAF output from ANGSD to a site frequency spectrum for both desert populations in each species. Detailed settings can be found in the supplementary information. We performed F_ST_ outlier analysis for our species using the calculated F_ST_ values. Z-scores for F_ST_ for each species were calculated using the formula ZF_ST_=(observedF_ST_-meanF_ST_)/SDF_ST_. We split the genome into two different partitions based on these z-scores: F_ST_ peaks, for values of F_ST_ greater than five standard deviations above the mean (z-score>5) and F_ST_ troughs for values of F_ST_ greater than five standard deviations below the mean (z-score<-5). We only report the F_ST_ peaks in the main manuscript: for F_ST_ troughs, see the supplementary information. We performed this outlier detection for the complete, 75%, and 50% missing datasets to assess if low coverage impacted our calls.

Recombination rates (in crossovers per base pair, c/bp) across the genome were estimated using the program ReLERNN (Adrion et al., 2020), assuming a mutation rate of 2.21×10^−9^ mutations per site per year (Nam et al., 2010) and a generation time of one year. This program combines simulation with a recurrent neural network to estimate the recombination rate on each chromosome in 100,000 bp windows. At present ReLERNN does not support genotype likelihoods, so we used SNPs in VCF format. We called SNPs using ANGSD with the following parameters: a p-value of 0.01; using the frequency as a prior; removing sites with a minor allele frequency below 0.05; a minimum mapping quality of 20; a minimum base quality score of 20; SNPs only called at a posterior probability greater than 0.95; minimum of four individuals with SNP.

### Morphological data

We quantified morphological variation in our 10 focal species to assess which of the geographic models best explain morphological variation across the landscape (see *Generalized Dissimilarity Matrix Models*). We measured 366 specimens (19–59 per species), excluding known females and known juveniles to account for any variation attributed to sex and age. Of those, 29 were also present in the genomic dataset, with 0–8 individuals per species.

We generated seven raw plus seven compound morphological measurements, which we designated as proxies for thermoregulation and dispersal, respectively (see Supplementary Information). We reduced the dimensionality of the 14 morphological measurements using a principal components analysis (PCA). We then calculated four distance matrices between individuals: one Euclidean distance matrix for all morphological variables, where we calculated the Euclidean distance between individuals among all raw and calculated measurements; and three Euclidean distance matrices for the first three principal components, PC1, PC2, and PC3. We assessed whether there were differences in morphological PCA space between the Sonoran and Chihuahuan Desert populations in each species using DABEST tests in the dabestr package version 0.3.0 (Figure 6; Supplementary Figure 19; Supplementary Figure 20; Ho et al., 2019). Note that this method does not give explicit significance values, instead it shows whether expected confidence intervals overlap zero (i.e., no difference between deserts) or not.

### Isolation across the landscape at different temporal resolutions

We calculated IBD matrices by calculating the Euclidean geographic distance between the latitude/longitude pair of each specimen in R. We used the WGS84 projection for all data. These variables were somewhat correlated with one another, though less so after accounting for geographic distance (Supplementary Figure 21).

To produce data for the IBH model, we calculated environmental resistances in the Last Glacial Maximum (LGM; ∼21,000 years ago) for each species. To do this, we created ecological niche models (ENMs) using 19 layers representing contemporary climate (WorldClim; Hijmans et al., 2005) at a resolution of 2.5 arcminutes. We used MaxEnt (Phillips et al., 2006), with ENMeval version 0.3.1 as a wrapper function for model selection (Muscarella et al., 2014). ENMeval optimizes MaxEnt models based on different sets of feature classes and regularization values (see Supplementary Information). The contemporary ENMs (see IBE section below) were then backprojected to the LGM using WorldClim paleoclimate data (Hijmans et al., 2005). We also backprojected to the Mid-Holocene, but contemporary and Mid-Holocene ENMs were highly correlated, so we excluded the Mid-Holocene values from downstream analyses. We then scaled the LGM suitability values to range between 0–1 and calculated resistances across the environment using the least cost path distance method in ResistanceGA version 4.0–14 (Peterman et al., 2014, Peterman, 2018). Regions of high resistance are predicted to reflect poor habitat and be costly to traverse through. The ENMs were thresholded to equal sensitivity-specificity values for visualization (Supplementary Figure 22).

We approximated IBB by assigning individuals based on their location relative to the Cochise Filter Barrier (see Supplementary Information). For proximity to the Cochise Filter Barrier, we assigned individuals to either Sonoran or Chihuahuan populations either based on the results of the K=2 clustering analysis, if there was structure across longitudes, or according to a cutoff of longitude if there was no structure. We chose 108 °W longitude as our cut off— individuals west of this point were deemed Sonoran, and individuals east of this point were deemed Chihuahuan (but see Provost et al., 2021). In some cases, species with genetic breaks had some uncertainty due to unsampled areas or admixed individuals—we labeled these individuals as being unclear with respect to their desert assignment. Georeferencing on some morphological specimens was poor, but all except two specimens (see Results) were identified at least to county level if not to a specific locality. When localities were given, we georeferenced the specimens to the nearest latitude/longitude. Otherwise, we assigned individuals to the centroid of their state or county.

We independently tested IBE by using two datasets: contemporary environmental distance and resistance. For the environmental distances, we used the 19 WorldClim bioclimatic layers (see IBH section). For the latitude/longitude location of each specimen used in both the morphological and genomic analysis, we extracted the values on those WorldClim layers and then calculated the Euclidean distances in environmental space between specimens. This gave us an estimate of how different the environments were at each specimen’s locality. For the environmental resistances, we created ENMs using the WorldClim layers, then added layers for soil properties, distance to water, terrain features, and vegetation, and occurrence data for the focal species (see Supplementary Information). We then calculated resistances and thresholded as described above.

To assess IBA, which had a temporal scale of the last 50 years, we obtained abundance information from the Breeding Bird Survey (Pardieck et al., 2019). This dataset consists of replicated transects where individual birds are counted across the whole of the United States. The methodology for counting is standardized and covers multiple decades of observations, with our dataset comprising data from 1966–2018. We downloaded raw data for all points, then subsetted our data to our ten focal species. We averaged the number of individuals across years (though some points only had a single year). We then interpolated across points using inverse distance weighted interpolation in the spatstat version 2.1-0 package in R (idp=5). The interpolations were converted to rasters with extents and resolutions matching those of the ENMs. We then calculated resistances such that regions of high abundance had low resistance, to generate an abundance distance matrix between individuals.

### Generalized dissimilarity matrix models

We assessed the relative effect of alternative geographic models on intraspecific variation in our focal species by building generalized dissimilarity matrix models (GDMs). As spatial layers representing our five models, we calculated geographic distances, abundance resistances, environmental distance and resistance, separation by barrier, and paleoenvironmental resistance between all individuals in each species. The models likely represent different temporal resolutions, from millions of years ago to the present-day configuration of the barrier. These predictors served as the input parameters for our GDMs and will be discussed in detail below. With our numerous response matrices (four morphological matrices, three genome matrices for each missing data cutoff, 35 matrices for chromosomes, five matrices for the lostruct partitions, and six matrices for the F_ST_ outliers with missing data cutoffs) and our six predictor matrices (with two for IBE: environmental distance, environmental resistance), we generated generalized dissimilarity matrix models using the gdm package version 1.3.11 in R (Manion et al., 2018). We tested which of IBA, IBB, IBD, IBE, IBH, or a combination best explained the variation in the response matrix (see below). Not all species had all chromosomes sequenced, and not all models converged: we have omitted those data. For each of the 45 response matrices per species, we built a univariate model where the genomic/chromosomal variable was predicted solely by one of the six predictor matrices. We also built models with combinations of two (bivariate) or three variables (trivariate), which we present in the Supplementary Information. Further, we present the GDM results for the chromosomes in the supplementary information. We compared the models based on the highest percent deviance explained.

To identify any overarching patterns with respect to which model of landscape evolution best explained genetic diversity (Supplementary Figure 23), we calculated four summary statistics for each chromosome, each lostruct and F_ST_ outlier partition, and the genome as a whole. We tested whether genomic summary statistics on each chromosome (F_ST_, D_XY_, missing data, recombination rate) were correlated with explained percent deviance with an analysis of variance (ANOVA) test and a Tukey’s honest significant difference test (Chambers et al., 1992, Miller, 1981, Yandell, 1997) using the stats v. 3.6.1 package in R. We did this for the complete dataset; for 75% and 50% missing data datasets, see Supplementary Information. We also calculated linear models comparing the proportion of each model to species-wide estimates of habitat suitability across the barrier. For all significance tests, we used an alpha value 0.05. However, due to multiple model testing for the GDM analyses, we applied a Bonferroni correction for simultaneous testing of six univariate models, with a final corrected alpha value of 0.0083 as our cutoff for all GDM tests (Bonferroni, 1936).

We evaluated whether the best-predictors of genomic landscapes varied across species and across partitions of the data using Chi-squared tests of significance, via the chisq.test function in the stats package in R. For each, the expected distributions assuming no differences between species, partitions, or structure were calculated and compared to the observed distributions. Chi-squared tests were performed both with and without Monte Carlo simulations (N=2000 simulations each repeated 1000 times).

## Acknowledgements

This work would not have been possible without generous specimen loans from DMNH (J. Woods), UWBM (R. Faucett, J. Klicka, S. Birks), UMMZ (J. Hinshaw, B. Benz), TCWC (G. Voelker), MSB (M. L. Campbell, M. Andersen, C. Witt, A. Johnson, J. McCullough), LSUMZ (D. Dittmann, F. Sheldon), CUMV (V. Rohwer, C. Dardia), AMNH (P. Sweet, P. Capainolo, B. Bird, T. Trombone). We are grateful to numerous State and Federal Collection Permit officers, and many BLM managers (T. Schnell, S. Cooke, M. McCabe, J. Atkinson, M. Daehler, S. Torrez, D. Tersey). Thanks to staff at Dalquest Desert Research Station (N. Horner) and Indio Mountains Research Station (J. Johnson). We thank M. Ingala for illustrating the birds used in many of our figures and for helpful feedback. Helpful input comes from the Smith Lab, S. Simpson, L. Musher, D. Fletcher, F. Burbrink, L. Alter, D. Kelly, I. Overcast, A. Xue, M. Hickerson, M. Blair, P. Galante, R. Harbert, E. Sterling, A. Xue, and E. Myers., the Underrepresented Genders in Museum Ornithology group, and the B. Carstens lab. We also thank two anonymous reviewers for their helpful feedback. This work was funded by the AMNH Frank M. Chapman Fund, American Ornithological Society, Society of Systematic Biologists, RGGS Sydney Anderson Travel Award, AMNH Linda H. Gormezano Fund, and AMNH RGGS Graduate Fellowship. BTS was supported by US NSF award DEB-1655736. KLP was supported by US NSF award DEB-2016189.

## Data Availability

These scripts used to perform these analyses are found at https://github.com/kaiyaprovost/GDM_paper/. All data used to perform analyses will be available on Dryad upon publication.

## Notes

### Competing Interest Statement

The authors have declared no competing interest.

### Summary of Updates

Peer review was done on the manuscript to make major changes. Most notably, calculation of genetic distances were converted from hard-called SNPs to the ngsDist protocol that can handle genotype likelihoods. As such we are more confident in these data. All figures, text, and supplement revised.

