## Supplemental Text for "The genomic landscapes of desert birds form over multiple time scales"

Supplementary Information for: **The genomic landscapes of desert birds form over multiple time scales**

### Supplementary Text

#### Genomic extraction, sequencing, and analysis details

Aligning individuals was done using the following references from GenBank: GCF_000691975.1, GCF_001522545.2, GCF_000385455.1, GCF_000277835.1, GCF_001447265.1, respectively.

#### Chromosomal and missing data supplementary results for summary statistics

Of the 221 individuals we sequenced, 199 were retained in the 75% missing dataset and 93 were retained in the 50% missing dataset. Mean X coverage for the 75% missing dataset was a mean of 3.3x per individual (0.4x–8.8x) with species means ranging from 2.3x–4.3x (*Phainopepla nitens* lowest and *Melozone fusca* highest). Mean X coverage for the 50% missing dataset had a mean of 4.1x per individual (0.6x–8.8x) with species means ranging from 2.7x–5.2x (*Phainopepla nitens* lowest and *Cardinalis sinuatus* highest).

Compared to the whole genome, chromosomes tended to vary with respect to their summary statistics. When all species are pooled, smaller chromosomes had higher average distances and were more discordant than larger ones (log number of scaffolds vs mean normalized RF distance R^2^=0.78, p=4.90x10^-13^). *Melzone fusca* had the lowest normalized RF distance across chromosomes (mean±standard deviation [SD] 0.724±0.200) whereas *Phainopepla nitens* had the highest distance (0.945±0.176; Supplementary Figure 4). *Campylorhynchus brunneicapillus* was the least variable across chromosomes (0.855±0.170) whereas *Melozone fusca* was the most variable.

Lostruct outliers were variable across chromosomes. In addition, the Z chromosome tended to have proportionately more loci fall into an outlier category (76.0% non-outliers, 4.10–11.2% range for LS1, LS2, LS3), usually one single category. Other chromosomes that show high proportions of outliers in at least one species included 1B, 4, 4A, 7, 8, 11, 22, 25, 27, 28, and LGE22. When visualized across the chromosomes, some species appear to have a strong association between F_ST_ and lostruct outlier status.

The mean D_XY_ for species varied from 0.0072–0.0159. D_XY_ was significantly correlated between the 100% and 75% and the 100% and 50% missing data partitions with a moderate R^2^ (p<0.0001 for both, range adjR^2^=0.35–0.36), but had a much higher magnitude of correlation between the 75% and 50% missing data partitions (p<0.0001, adjR^2^=0.87).

The mean F_ST_ per species varied from 0.020–0.119. F_ST_ was significantly correlated between the 100% and 75% missing data partitions with a high R^2^ (p<0.0001, adjR^2^=0.77). It was also significantly correlated between 100% and 50% (p<0.0001, adjR^2^=0.004) and 75% and 50% (p<0.0001, adjR^2^=0.02), but the magnitude of the correlation was highly reduced. Average F_ST_ across missing datasets is overall higher in the datasets with fewer individuals. Notably, the *Campylorhynchus brunneicapillus* and *Phainopepla nitens* F_ST_ values for the 50% missing data partition are approximately ~0.2, but they both have only one individual in the Chihuahuan desert, thus inflating fixation. Six species had low-F_ST_ outlier partitions identified in the complete dataset and F_ST_ low outliers ranged from 0–679, with fewer generally identified in species with higher F_ST_. Further, lostruct outliers generally contained 0.0–7.1% (mean 0.3%) of the F_ST_ troughs. Compared to the 100% missing data partition, the number of F_ST_ outliers was reduced in the 75% and 50% missing data partitions. These values for the 75% missing data partition were 33–753 high F_ST_ outliers and 0–674 F_ST_ low outliers, and for the 50% missing data partition was 0–425 high F_ST_ outliers and 0–709 low F_ST_ outliers. None of the F_ST_ low outliers were significantly different across models.

#### Cline analysis supplementary results

There is a negative correlation between F_ST_ and mean cline width such that species with more fixation between populations have narrower clines (R^2^=0.60, adjusted-R^2^=0.55, p=0.008; Supplementary Figure 5). Mean center location also shows a negative trend with F_ST_ but it is not significant (R^2^=0.03, adjusted-R^2^=-0.09, p=0.65). The standard deviation of cline width shows a positive trend with F_ST_, but it is also not significant (R^2^=0.11, adjusted-R^2^=0.00, p=0.34). The standard deviation of cline centers is negatively correlated with F_ST_, such that species with more fixation between populations also have less variation in the locations of their clines (R^2^=0.42, adjusted-R^2^=0.35, p=0.04). This supports the prediction that species with more structure should have narrower clines that are closely centered with respect to each other. Within-chromosome variation in mean cline width and cline center location was reduced compared to species (range width 10.57–16.77°, range center 7.72–10.99°), though smaller-sized chromosomes tended to have the largest widths. Cline width does not vary significantly across models.

#### Morphological methodology details

To quantify morphological variation in our ten focal species, we measured 367 adult male museum round specimens. Using digital calipers, we measured the lengths of the beak, flight feathers, and tarsus. We excluded females and juveniles. Many older museum skins lack information about age and sex—we assumed that unlabeled birds were adult males unless plumage suggested otherwise. We measured seven measurements: tarsus length, bill height, bill width, and bill length, tail length, length of longest primary to shoulder, and length of longest secondary to shoulder, using digital calipers with a precision of 0.01 mm. From these measurements, we calculated compound traits including Kipp’s Index (Kipp 1959), bill lateral surface area, bill volume, and bill surface area to volume ratio (see Greenberg et al. 2012). We also calculated base area, total surface area, and total surface area to volume, and the relative lengths of all metrics to body size. Each specimen was measured three times to estimate repeatability, then the measurements were averaged (Goodenough et al. 2010). We also supplemented this data with pre-existing specimen measurements from specimen tags, which were typically only measured once during specimen preparation. We used those raw measurements to calculate beak surface area and volume and the relative aspect ratio of the wing (Kipp’s Index, Kipp 1959).

We calculated all beak measures assuming that beaks were conical—this may be an underestimate in species of *Toxostoma* which have pronounced curves to their bills. We tested whether calculating these values before or after averaging measurements impacted our results. Both sets of measurements were highly correlated (adjusted R^2^ values range 95–99%) and our conclusions were not affected by which set we used, so we chose to use values after averaging.We scaled and centered all raw and calculated measurements, unscaled and without log transforms. We assessed differences within species in morphospace using DABEST (Ho et al. 2019) to visualize differences between populations.

We calculated the relative lengths of all morphological metrics to body size, using tarsus length as a proxy for body size (Rising and Somers 1989, Senar and Pascual 1997). We did this by performing a linear model, where we predicted the metric of choice from tarsus length, and then took the residuals to be the corrected relative length. We also log transformed our metrics, both the absolute and relative lengths. These metrics were not used in our subsequent analyses, but they did vary highly from the uncorrected metrics.

We calculated PCA values on the morphological measurements twice: once with those values imputed to the species mean (i.e., imputed dataset, N=369), and once with individuals missing measurements excluded (i.e., non-imputed dataset, N=234). With all other analyses using morphological data, we only used the non-imputed dataset.

#### Morphological results details

Bill Height, length, and width, and beak base area, loaded most highly onto PC2 (absolute value range 0.30–0.42), Kipp’s Index loaded most highly onto PC3 (absolute value 0.85), and the rest loaded most highly onto PC1 (absolute value range 0.26–0.30).

According to DABEST tests, *Toxostoma crissale* had larger PC1 values in the Chihuahuan Desert than the Sonoran in both imputed (n=16 Sonoran, 12 Chihuahuan) and non-imputed datasets (n=7 Sonoran, 13 Chihuahuan). In the imputed dataset only, *Polioptila melanura* had larger PC1 in the Chihuahuan Desert (n=14 Sonoran, 18 Chihuahuan) and both *Cardinalis sinuatus* (n=22 Sonoran, 9 Chihuahuan) and *Auriparus flaviceps* (n=21 Sonoran, 19 Chihuahuan) had smaller PC1 in the sonoran desert. T-tests agree that the *Toxostoma crissale* (p<0.004), *Auriparus flaviceps* (p<0.05), *Cardinalis sinuatus* (p<0.0002), and *Polioptila melanura* (p<0.005) comparisons were significant. Additionally, in the imputed dataset, *Melozone fusca* (n=19 Sonoran, 34 Chihuahuan) was nearly significantly different (p=0.067), with Chihuahuan desert individuals having lower PC1.

*Vireo bellii* had larger PC2 in the Chihuahuan Desert in both imputed (n=11 Sonoran, 23 Chihuahuan) and non-imputed datasets (n=11 Sonoran, 33 Chihuahuan). *Melozone fusca* had smaller PC2 in the Chihuahuan Desert in both imputed and non-imputed datasets (n=4 Sonoran, 9 Chihuahuan). Like with PC1, some species were only significantly different in the imputed datasets: *Cardinalis sinuatus* had a larger PC2 in the Chihuahuan desert while *Toxostoma crissale* had a smaller PC2 in the Chihuahuan desert. T-tests agree that the *Melozone fusca* (p<0.05), *Toxostoma crissale* (p<0.04), and *Cardinalis sinuatus* (p<0.003) comparisons were significantly different. While the imputed *Vireo bellii* comparison was also significant (p=0.048), the non-imputed dataset was not, though it was very nearly significant (p=0.056).

For no species was PC3 significantly different in the DABEST tests or T-tests, although *Vireo bellii* (both imputed and non-imputed) and *Toxostoma crissale* (only imputed) are nearly significantly different (pval range 0.074–0.088). The remaining taxa (*Amphispiza bilineata, Campylorhynchus brunneicapillus, Toxostoma curvirostre, Phainopepla nitens*) were never significantly different in morphology across the CFB, suggesting that morphological variation in most of these species is not associated with the barrier itself, and instead other factors may be causing morphological variation to develop.

Across the three morphological metrics, there was no correlation between F_ST_ and differences in PC1, PC2, or PC3, as calculated by the absolute value of the differences in morphology divided by the absolute sum of the values (range p=0.031–0.088, range R^2^=0.00–0.012). The six species identified as having morphological differences all have either phylogeographic structure or show a gradient (though, notably, *Toxostoma curvirostre* does not show morphological differentiation), suggesting that this differentiation in morphology reflects differentiation between incipient populations.

#### Ecological niche modeling

We obtained occurrence data for the focal species by downloading all eBird data in May 2017 (Sullivan et al. 2009). We then supplemented this data with data from GBIF, iNaturalist, and VertNet (gbif.org, inaturalist.org, vertnet.org). GBIF data was retrieved from the following DOIs on October 9th, 2017: <https://doi.org/10.15468/dl.0jvomy>, <https://doi.org/10.15468/dl.9w3468>, <https://doi.org/10.15468/dl.87sdjs>, <https://doi.org/10.15468/dl.a1otjs>, <https://doi.org/10.15468/dl.ejntps>, <https://doi.org/10.15468/dl.h8aj9l>, <https://doi.org/10.15468/dl.lw0qwz>, <https://doi.org/10.15468/dl.nc3113>, <https://doi.org/10.15468/dl.riixpl>, <https://doi.org/10.15468/dl.vn53xq>, <https://doi.org/10.15468/dl.yze7lh>.

Obtaining the occurrence data was done using custom R scripts that made use of the following packages: raster (Hijmans 2019), MASS (Venables and Ripley 2002), spocc (Chamberlain 2017), rgeos (Bivand and Rundel 2017), dplyr (Wickham et al. 2017), sp (Pebesma and Bivand 2005; Bivand et al. 2013), and dismo (Hijmans et al. 2017). We removed duplicate localities from these datasets per species. Outliers were detected and removed using a probability density function. After this, we thinned the data using spThin (Aiello-Lammens et al. 2015) to account for spatial autocorrelation in the data. Before thinning, we downloaded a total of 115,285 occurrence points across the ten species. After thinning to one point per 10 km, we ended up with a total of 6,024 points (range 369–929 per species).

Datasets included the 19 WorldClim rasters (i.e., BIOCLIM variables, see Booth et al 2014), soil acidity and soil texture (SoilGrids, Hengl et al. 2017), distance to water (calculated from Commission for Environmental Cooperation Lakes and Rivers 2009), elevation, roughness, relief, and slope (NASA SRTM, Farr et al. 2007), land cover (CEC, Homer et al. 2020), and vegetation type (LandFire, Ryan and Opperman 2013). Note that the LandFire dataset is only available for the USA. We converted all of the datasets to a grid cell size of 0.002083333 latitude by 0.002083333 longitude (7.5 arcseconds) and cropped them to 115–97 °W longitude and 26–37 °N latitude. This was used both for the background training extent and the final projection extent. We did analyses with three versions of this data: 1) a climate-only dataset, which includes the 19 bioclim variables only; 2) a North America dataset, which includes all layers except the LandFire data; and 3) a USA dataset, which includes all of the above.

With ENMeval, we used the linear and linear+quadratic feature classes, and regularization values of 0.5, 1, 1.5, 2, 2.5, 3, 3.5, and 4. Models were run for each combination of feature classes and regularization values, and the model with the lowest occurrence point omission rate and highest average test area under the curve (AUC) was chosen We used the entire extent of the layers as background points, partitioned with the block method, and thresholded our models when appropriate at the equal sensitivity-specificity threshold. ENMeval analysis on the best model showed that linear-quadratic models always won out over linear models (Supplementary Table 7). The AUC values were overall high: for training data they ranged from 0.87–0.97, while for the test data they ranged from 0.83–0.96. Omission rates varied from 7%–26%.

For all of the species, present and mid-Holocene ranges largely overlapped (Supplementary Figure 22) with three species (*Toxostoma crissale, Toxostoma curvirostre, Auriparus flaviceps*) being connected with a large amount of habitat across the CFB, four connected with a very small amount of habitat (*Vireo bellii, Amphispiza bilineata, Melozone fusca, Cardinalis sinuatus*), and the rest having multiple regions of suitability (*Campylorhynchus brunneicapillus, Polioptila melanura. Phainopepla nitens*). The number of stable regions of suitable habitat varied from 1–4 across species. One notable pattern was that for *Polioptila melanura*, refugia during the Last Glacial Maximum was predominantly concentrated in the western part of their range.

We found that the environmental landscape for this avifauna is complex and variable between species. Despite these taxa all being desert birds, we find large amounts of variation in how much of the region is suitable for them, their estimated distributions during the Last Glacial Maximum, and their abundance. We suspect that this pattern of heterogeneity is not unique to the CFB, but rather is present in many communities co-distributed across barriers. This discordance and heterogeneity may characterize the early stages of evolution and diversification in avian communities, where the factors impacting one group of taxa are not the same as the factors impacting another.

We quantified the amount of suitability across the Cochise Filter Barrier by constraining the habitat to a bounding box of 100 °W to 115 °W longitude and 30 °N to 35 °N latitude, and then taking the average suitability for that longitude value. We find that for all species there is a decrease in suitability at ~108.5 °W and ~106 °W longitude, with the area in between those being a relative peak in suitability. Outside of this region, species vary but generally the regions of the deserts proper have relatively high suitability in comparison to these regions of low suitability. Some taxa show a sharp decrease in the extreme eastern or extreme western parts of this range. Contemporary climatic suitability shows a sharp drop-off in the far western portions of all species’ ranges and a more protracted drop-off in the eastern portions (Supplementary Figure 8). Within the contact zone, mean scaled suitability ranges from 48% (*P. nitens*) to 71% (*C. sinuatus*), and the standard deviation ranges from 12% (*A. bilineata*) to 18% (*P. nitens*).

Environmental variables are correlated with each other across species (Supplementary Figure 21), with 80/150 of the variables having higher than 75% correlations, and always being positive. IBB is the least correlated amongst variables, followed by environmental distance (IBE); the remaining variables (environmental resistance [IBE], IBH, IBD, and IBA) are all roughly equally correlated. However, when geographic distance is accounted for by taking the residuals of environmental variables vs IBD, the degree of correlation decreases for most species with only 4/100 of the variables having higher than 75% correlations, and many being negative (note that all comparisons with IBD are excluded from the latter). The four correlated variables after accounting for IBD are always environmental resistance (IBE) wth IBH; both of these variables are resistance matrices calculated from ecological niche models. These are also highly correlated before accounting for IBD (correlation range 0.99-1.00) and are likewise highly correlated with IBD (correlation range 0.96-1.00).

#### Abundance data for IBA analyses

From breeding bird survey data, we extracted 27,154 data points for our 10 focal species (range 622–6,232 per species) across the years 1967–2018. This was a total of 687 routes, with a range of 93–850 data points per year. Many taxa are predicted to have a sharp drop-off in abundance in the region of the Cochise Filter Barrier.

#### Generalized dissimilarity matrix model inputs and details

In addition to the univariate models we described in the main text, we also ran bivariate models and trivariate models. The bivariate models accounted for structure, here defined as a distance matrix where individuals assigned to different sides of the CFB had a distance of 1, individuals who were assigned to the same side had a distance of 0, and individuals whose assignment was ambiguous (across datasets or due to high levels of admixture) had a distance of 0.5 to all other individuals. These assignments were based off of the K=2 analyses for all taxa. In the species that did not show longitudinal population structure, we assigned them to sides of the CFB based on where they fell relative to 108 °W longitude, approximately where the peak of contact zones is across the barrier (Provost et al. 2021). IBB was used as an effect in all bivariate and trivariate models such that we were only testing between the other predictors. The trivariate models additionally had IBD included as an effect. The univariate models, in contrast, used IBB as a predictor. We chose not to do trivariate models with multiple predictor matrices at once (as static effects not random effects) because the sample sizes in our data precluded us from using models with high complexity.

Univariate models had the formula as follows: genetic/phenotypic distance matrix ~ one of six predictor matrices (geographic distance, abundance resistance, environmental distance, environmental resistance, paleoclimate resistance, or structure distance). Bivariate models had this formula: genetic/phenotypic distance matrix ~ structure distance)+ one of the five remaining predictor matrices. Trivariate models instead had this formula: genetic/phenotypic distance matrix ~ structure distance + geographic distance + one of the four remaining predictor matrices.

GDMs are evaluated using three parameters: the null deviance, the model deviance, and the percent deviance explained. The null deviance was how much of the variation in the data is explained when no predictors are included (just fitting the intercept term). Model deviance becomes the variation explained when the predictors are added, and percent deviance explained is the difference between the two (scaled as a percentage of the null deviance).

#### Univariate, bivariate and trivariate model results

Across the 346/360 univariate chromosomal partitions that were successful, percent deviance explained by the univariate models ranged from 0.0% to 36.6% with a mean±SD of 10.0%±8.2%. All species had low-F_ST_ outlier partitions identified in the 100% missing dataset; all of those species had converged for the 100% and 75% datasets, but two did not for the 50%. All 10 species had high-F_ST_ partitions identified, and all converged across the 100%, 75%, and 50% datasets.

2,444/2,650 bivariate models converged successfully. In nearly all bivariate models, which contained IBB plus another predictor, IBB explained the highest percentage of the percent deviance. Across all of the data, 512 tests were successful in both univariate and bivariate models, and 78.3% of them have the same best model selected (if the 86 univariate models with IBB are excluded, this percentage is 94.1%). For the entire genome, *A. flaviceps* failed to converge for bivariate and trivariate models. 1/9 species had IBA as the most important predictor, 2/9 had IBH, 3/10 had IBE, and the remaining one had a mixture. There was some congruence across morphological partitions, with 23 of the partitions being shared within species. 360 total chromosomal partitions were analyzed, and 176 of those matched that of their respective species’ whole genome. Of all chromosomes, 107 showed IBE as the best predictor, 67 showed IBA, 70 showed IBH, 18 showed IBD, and the remaining 82 had mixed support. The percent deviance explained ranged from 0.08%–81.9% with a mean of 11.5%. As with univariate models, F_ST_, D_XY_, and missing data significantly differ across models with different best predictors.

1,911/2,120 trivariate models converged successfully. When we examined the trivariate model results, which contained IBB, IBD, and one other predictor, the best models explaining variation in the genome were 3/9 IBH, 2/9 IBE, 1/9 IBA, and the rest mixed support. 344 total chromosomal partitions were analyzed and 168 matched the genomes of their respective species. 128 chromosomes were explained by IBE, 75 by IBA, 72 by IBH, and the remainder by a mix of models. Across all datasets, 89.5% matched between bivariate and trivariate models (94.5% if IBD models are excluded) and 72.7% matched between univariate and trivariate models (94.3% if IBB and IBD models are excluded). Percent deviance tended to be comparable to univariate or bivariate models (range 0.09%–81.9%, mean 10.8%). Just like bivariate models, F_ST_, D_XY_, and missing data significantly differ across models with different best predictors.

The bivariate and trivariate models performed similarly to the univariate m models with respect to whether or not species, structure, or datasets were significantly different. and all were significant in both simulated and not-simulated tests (species-specific χ^2^>562.7, p<0.0005; partition-specific χ^2^<238.3, p>0.87; structure-specific χ^2^>95.5, p<0.0005).

#### Species-by-species analysis of GDM results for all models

*Vireo bellii:* Though Bell's Vireo shows structure across the CFB which would suggest IBB would be the best model, the best explanation for genomic variation in the species is instead a mixture of IBE and IBH. 22 chromosomes (including Z and mtDNA), all of the high-FST outliers, two of the low-FST outliers, two of the non-FST outliers, two lostruct partitions, and morphological PC2 (beak shape) are also explained by IBE. 6 chromosomes, the remaining low-FST outliers, and PC3 (wing shape) are explained by IBH. Some sections of the genome do show IBB as being the best model: chromosome 1B, one lostruct partition, and PC1 (body size). For the other models: overall morphology and one chromosome is best explained by IBA; three chromosomes and the non-outlier lostruct partitions are explained by IBD.

*Amphispiza bilineata:* The Black-throated Sparrow was unstructured across the CFB, though the overall genomic pattern is best supported by IBE, as are 15 chromosomes (including Z). Despite this, a higher proportion of partitions are best supported by IBB: 15 chromosomes (including mtDNA), the non-outlier and two outliers lostruct partitions, one high-FST partition, and a non-FST outlier partition. Notably, all morphological PCs are best explained by IBA, despite overall morphology being best explained by IBH.

*Campylorhynchus brunneicapillus:* The Cactus Wren’s overall genome is best supported by IBA for univariate models, despite showing panmixia across the CFB. 15 chromosomes, one high-FST partition, non-outlier FST partition, the non-outlier lostrcut partition, two outlier lostruct partitioons, overall morphology, and PC2 (beak shape) all also have IBA as the best model. The Z chromosome is best explained by IBE, while PC1 (body size) and PC3 (wing shape) are best explained by IBD.

*Toxostoma crissale:* The Crissal Thrasher shows a gradient in phylogeographic structure across the CFB and its genome is best explained by IBH in univariate models. Nearly all of its partitions are also explained by IBH. Notable partitions that are not include overall morphology and Z chromosome, which are explained by IBB; and PC2 (beak shape) and PC3 (wing shape), which are explained by IBA.

*Toxostoma curvirostre:* The Curve-billed Thrasher’s genome, in contrast to its congener, is best explained by a mixture of models: IBD, IBE, and IBH. This same mixture explains 28 additional partitions and an additional 9 partitions are explained by similar mixtures involving IBE and IBH. However, the Z chromosome and PC1 (body size) are best explained by IBE alone, and overall morphology is best explained by IBD.

*Auriparus flaviceps:* The Verdin shows structure in its phylogeography, with its genome having IBB as the best explanatory factor. Despite this, it is primarily explained by IBA, with 33 partitions (including the mtDNA and Z chromosomes). The only chromosome to also be explained by IBB is chromosome 1B. Overall morphology is best explained by IBE, and PC3 (wing shape) is best explained by IBH.

*Melozone fusca:* The Canyon Towhee shows a gradient in phylogeographic structure, though its genome shows IBB as the best model. 11 chromosomes, including the Z chromosome, are also best explained by IBB. However, 16 chromosomes are instead explained by IBE. The mtDNA, in contrast, as well as PC1 (body size) are explained by IBH. Overall morphology is best explained by IBA, and the other two morphological PCs are explained by IBD.

*Polioptila melanura:* The Black-tailed Gnatcatcher shows phylogeographic structure (for K=3 in particular), but its genome is best explained by a combination of IBD, IBE, and IBH. 31 of the chromosomes (including the Z) are also explained by a combination of some of these factors. Overall morphological differentiation is best explained by IBH, PC1 (body size) is best explained by IBD, PC2 (beak shape) is best explained by IBA, and PC3 (wing shape) is best explained by IBE.

*Phainopepla nitens:* Despite the Phainopepla showing panmixia, its genome is best explained by IBH, as are 26 of its chromosomes (including the Z). An additional 7 chromosomes as well as overall morphological differentiation and PC2 (beak shape) are explained by IBA. PC1 (body size) is explained by IBB, and PC3 (wing shape) is best explained by IBE.

*Cardinalis sinuatus:* The Pyrrhuloxia has a gradient in its phylogeographic structure and its genome has IBB as the best explanatory model, as well as 34 chromosomes (including mtDNA). The Z chromosome is best explained by IBE. Overall morphological differentiation is best explained by IBH, PC1 (body size) is best explained by IBD, PC2 (beak shape) is best explained by a combination of IBA, IBD, IBE, and IBH, and PC3 (wing shape) is best explained by IBA.

### Supplementary Figures

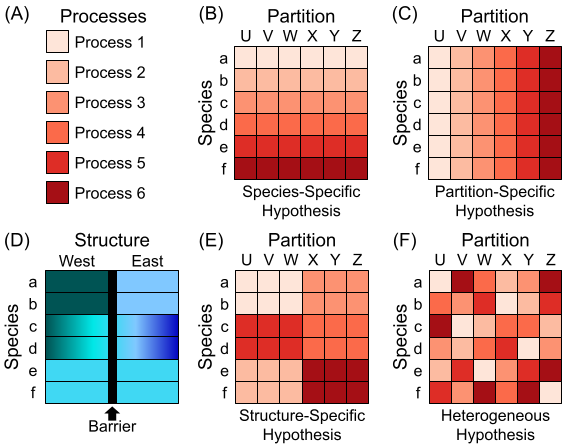

Supplementary Figure 1: Hypotheses of evolution investigated in this study for six processes (1-6), six species (a-f), and partitions of genotypic and phenotypic data (U-Z). (A) Six processes of evolution are investigated, with the processes being color-coded in shades of red. (D) Species are all distributed across a barrier that splits the species into western and eastern populations. Two species have clear phylogeographic structure at barrier (a, b); two species show a gradient from west to east (c, d); two species show no structure at barrier (e, f). Population structure is shown in shades of blue. (B) In the species-specific hypothesis, each species is impacted holistically by a single process across all partitions. (C) In the partition-specific hypothesis, each partition is impacted holistically by a single process across all partitions. (E) In the structure-specific hypothesis, the processes impacting species vary with population structure, with a gradient, or with no structure are each differentiated from each other. (F) In the heterogeneous hypothesis, there is no association between processes and species, partitions, or structure.

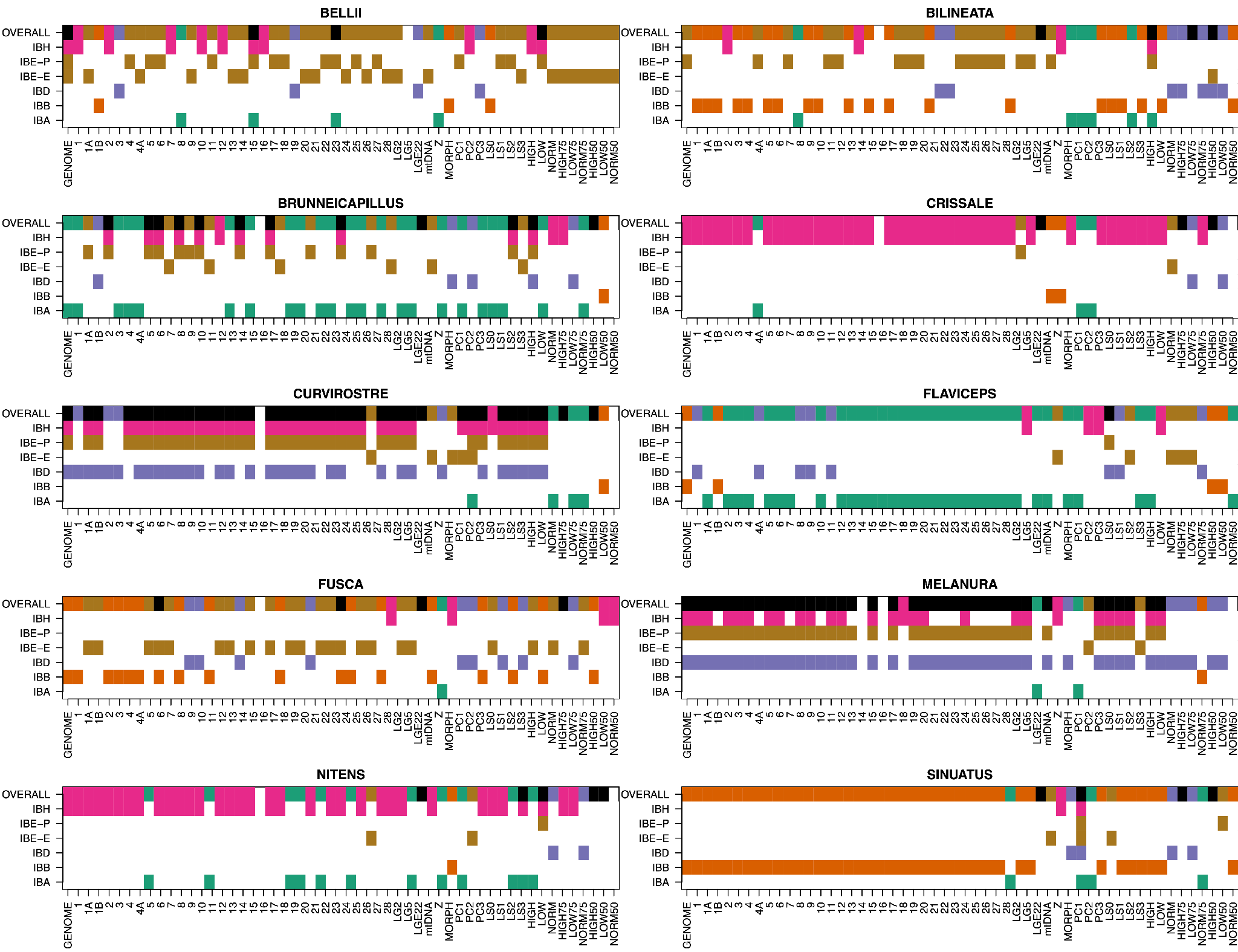

Supplementary Figure 2: Multiple spatial models contribute to patterns within partitions of genomic and phenotypic data. Each panel shows results for one species (species epithet above panel). Y-axis shows best performing spatial parameters in a GDM model, with full result along top (“OVERALL”) and each other line giving results for one individual geographic spatial model. The presence of color indicates that the spatial model in question is one of the best predictors. The alternative models were as follows: isolation by abundance (IBA), isolation by barrier (IBB), isolation by distance (IBD), isolation by environment for both predicted resistance (IBE-P) and raw environmental variables (IBE-E), and isolation by history (IBH). Colors are as in Figure 4; black in the overall models indicates a mixture is present.

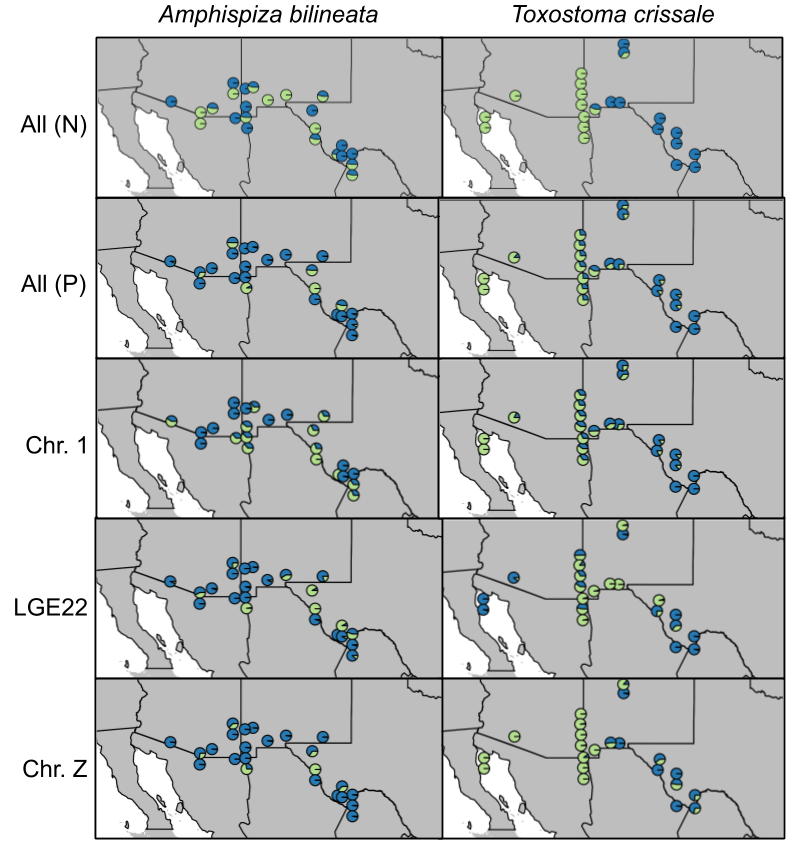

Supplementary Figure 3: comparison across genomes and chromosomes for two species, *Amphispiza bilineata*, one of our least differentiated species (left), and *Toxostoma crissale*, one of our most differentiated species (right). Panels from top to bottom indicate different partitions of the genome, from the whole genome (“All P” and “All N”) to individual chromosomes and scaffolds (“Chr. 1”, “LGE22”, “Chr. Z”). Analysis for “All P”, “Chr. 1”, “LGE22”, and “Chr. Z” used PCAngsd whereas analysis for “All N” used NGSadmix. Note that the LGE22 scaffold is much smaller than either of the chromosomes. Pie charts show individuals with colors giving the proportion of each cluster. Individuals at the same locality are jittered by latitude, though their longitude is kept the same.

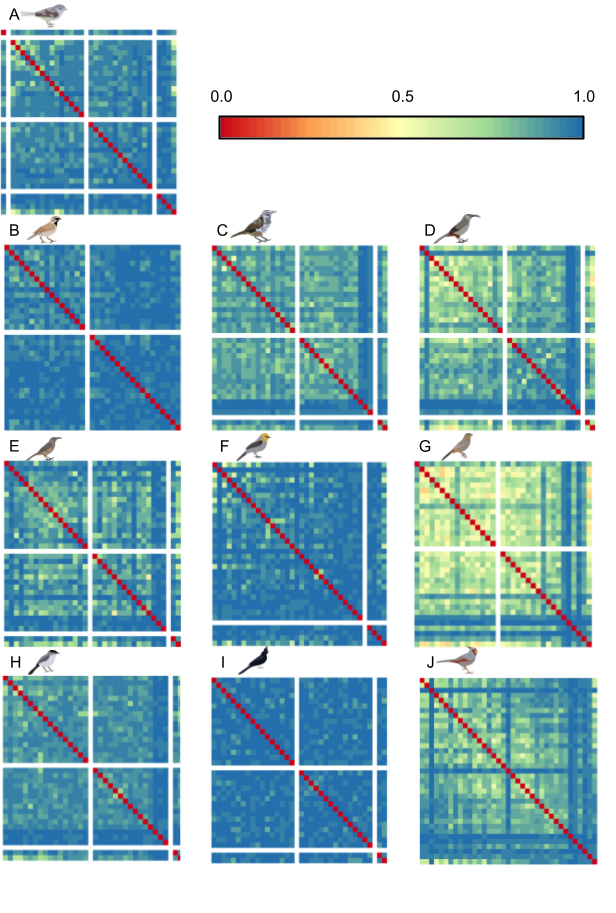

Supplementary Figure 4: heterogeneity in chromosomal Robinson-Foulds distances. Each row and column corresponds to one chromosome, with the right-most column and bottom-most row giving values for the entire genome. Cooler colors show comparisons with higher distances (more different) and warmer colors show comparisons with lower distances (more similar) with red indicating identical topologies. Rows and columns with all-white cells indicate that the corresponding chromosome was not available for that taxon (typically chromosomes 1A, 16, and LGE22). Chromosome order as in Figure 4A. A) *Vireo bellii*, B) *Amphispiza bilineata*, C) *Campylorhynchus brunneicapillus*, D) *Toxostoma crissale,* E) *Toxostoma curvirostre*, F) *Auriparus flaviceps*, G) *Melozone fusca*, H) *Polioptila melanura*, I) *Phainopepla nitens*, J) *Cardinalis sinuatus*.

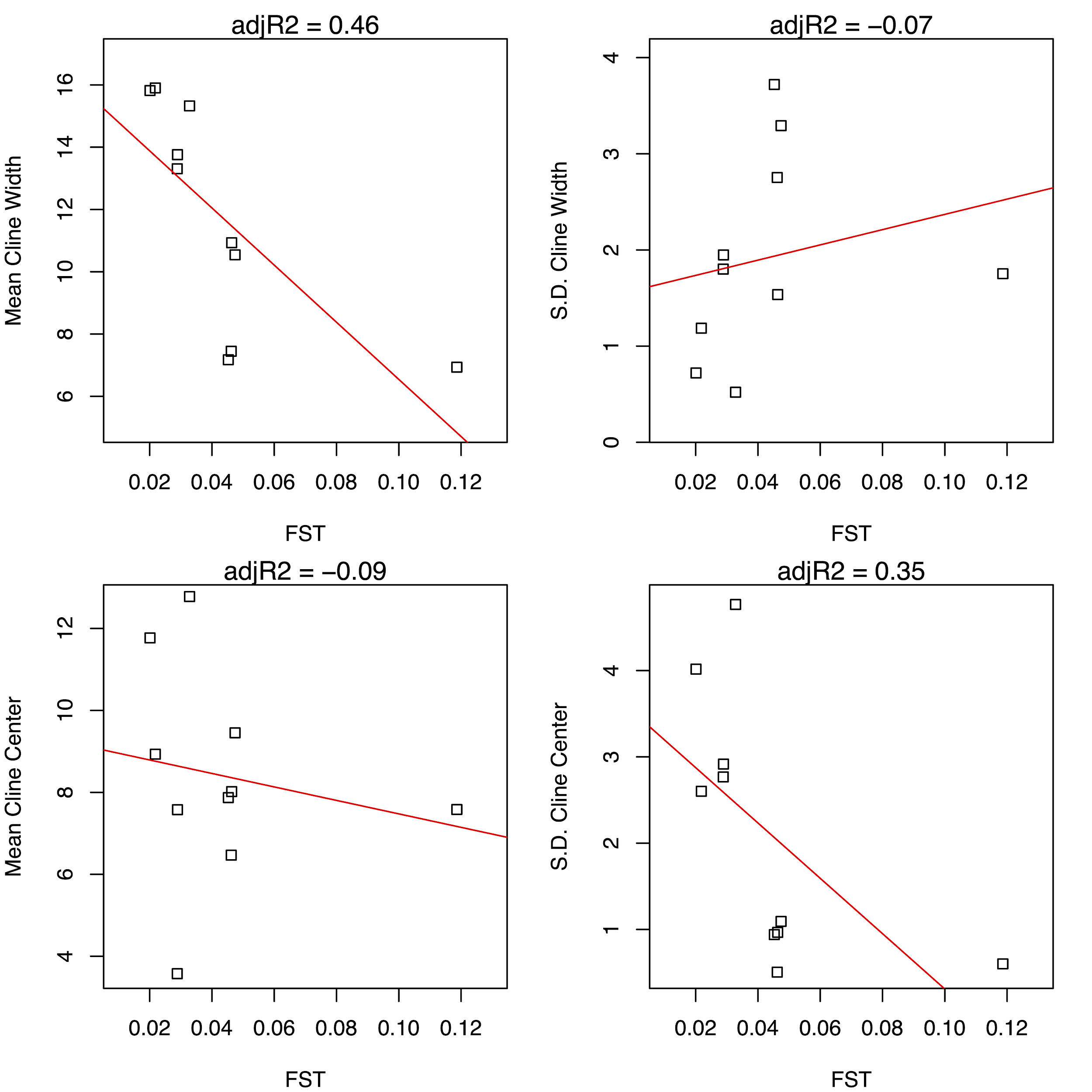

Supplementary Figure 5: Cline metrics (y-axis) vs F_ST_ (x-axis) averaged for each species. Top left: mean cline width. Top right: standard deviation (S.D.) cline width. Bottom left: mean cline center location. Bottom right: S.D. cline center location. Line through points show the line of best fit between measurements. Adjusted R^2^ values (adjR2) for lines of best fit are shown above each plot.

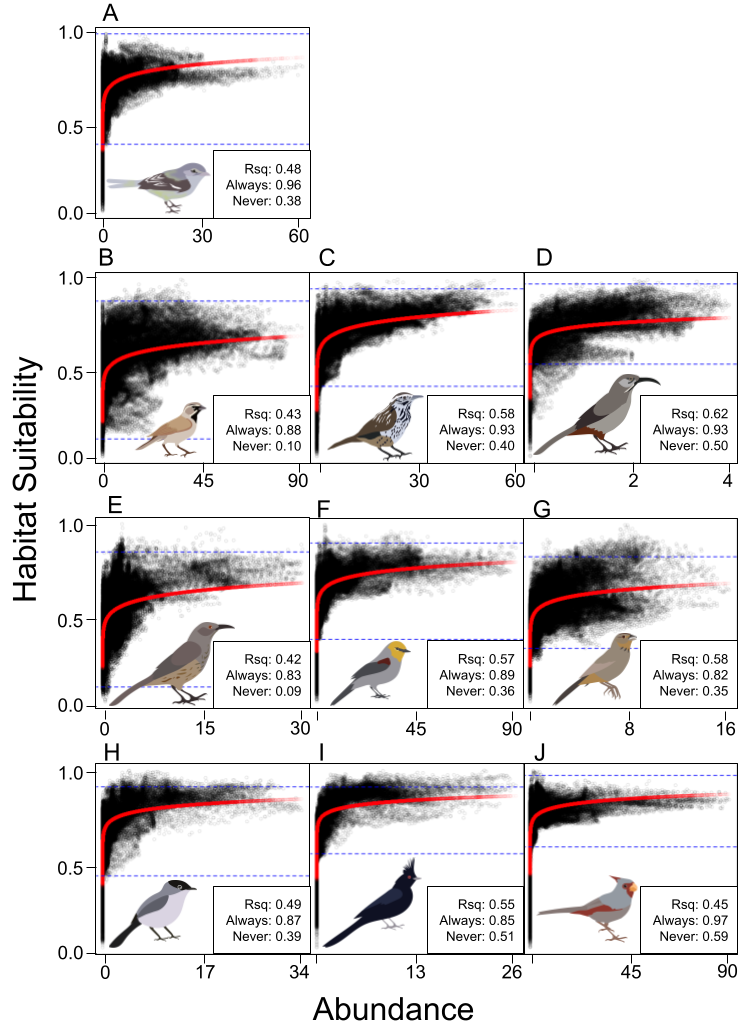

Supplementary Figure 6. Interpolated abundance vs. relative habitat suitability for all species. Red line indicates a line fit to *y=log(x)+b*. Each point represents one raster cell. Points are semi-transparent to show the density of points. Dotted blue lines separate, from top to bottom, habitat suitability above which species is always found, habitat suitability in which the species is sometimes found, and habitat suitability in which the species is never found. R^2^ (“Rsq”) and the suitability values for the “Always” and “Never” dotted blue lines are given in the bottom-right corner of the graph. A) *Vireo bellii*, B) *Amphispiza bilineata*, C) *Campylorhynchus brunneicapillus*, D) *Toxostoma crissale*, E) *Toxostoma curvirostre*, F) *Auriparus flaviceps*, G) *Melozone fusca*, H) *Polioptila melanura*, I) *Phainopepla nitens*, J) *Cardinalis sinuatus*.

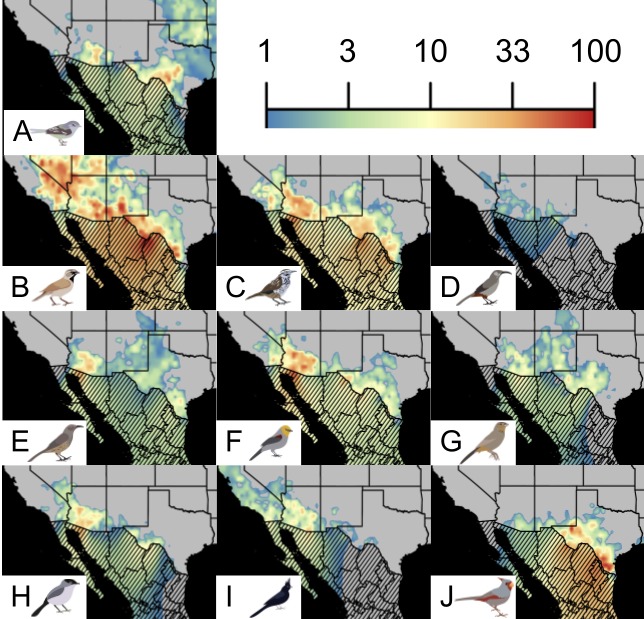

Supplementary Figure 7: estimated abundances across space for species, averaged over all years. Color is proportional to log abundance. Mexico is shaded out due to the lack of BBS points in that country, but the interpolated projections are included. A) *Vireo bellii*, B) *Amphispiza bilineata*, C) *Campylorhynchus brunneicapillus*, D) *Toxostoma crissale*, E) *Toxostoma curvirostre*, F) *Auriparus flaviceps*, G) *Melozone fusca*, H) *Polioptila melanura*, I) *Phainopepla nitens*, J) *Cardinalis sinuatus*.

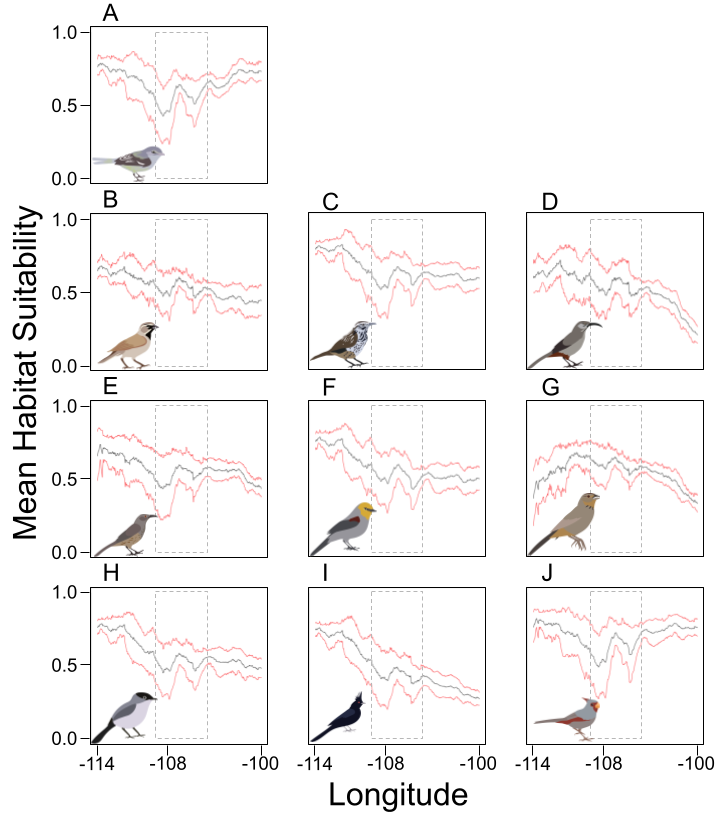

Supplementary Figure 8: Habitat suitability around the Cochise Filter Barrier varies across taxa, but declines within the contact zone. Mean scaled suitability for species across longitude, between -114 and -110 degrees longitude and 30 and 35 degrees latitude. Black lines show mean suitability, while red lines show one standard deviation above and below mean. Dotted gray square shows contact zone (Provost et al., 2021). Species are as follows: A) *Vireo bellii*, B) *Amphispiza bilineata*, C) *Campylorhynchus brunneicapillus*, D) *Toxostoma crissale*, E) *Toxostoma curvirostre*, F) *Auriparus flaviceps*, G) *Melozone fusca*, H) *Polioptila melanura*, I) *Phainopepla nitens*, J) *Cardinalis sinuatus*.

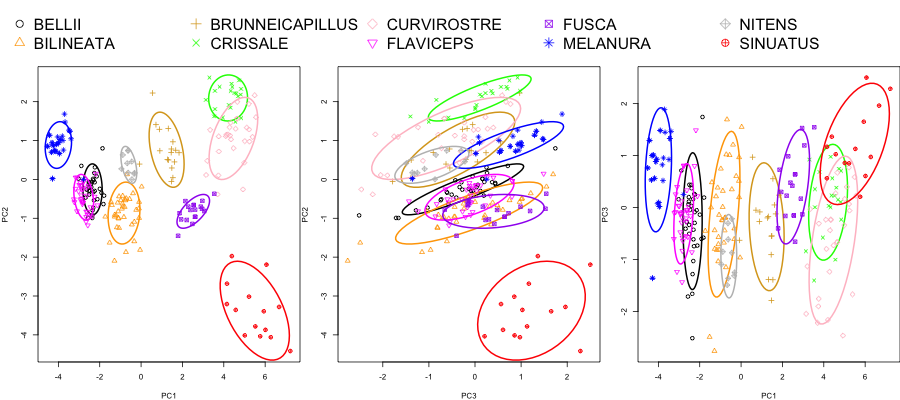

Supplementary Figure 9: Principal components analysis plots for morphological measurements. Colors and shapes correspond to different species. Lines indicate 50% confidence interval area for the mean of each species. Left: PC1 (x-axis) vs. PC2 (y-axis). Center: PC3 (x-axis) vs. PC2 (y-axis.) Right: PC1 (x-axis) vs. PC3 (y-axis.)

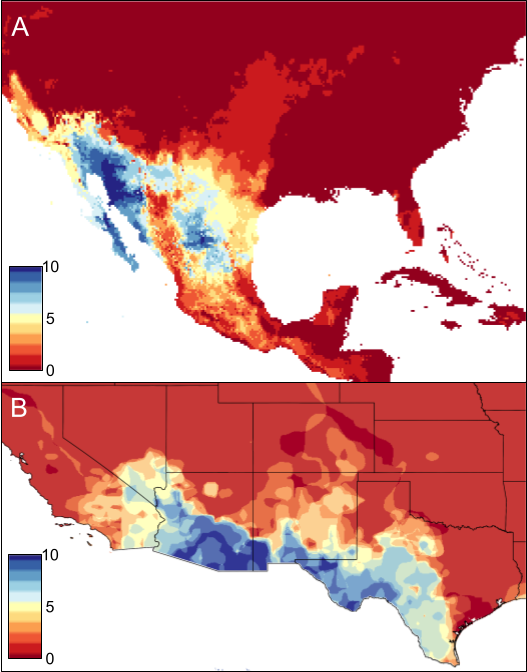

Supplementary Figure 10: Across species, some regions show higher abundance and higher habitat suitability. A) Predicted stability across all species during present and Last Glacial Maximum periods. Warmer colors indicate fewer species show stability in those regions (i.e., have cells where predicted presence>1 across all three time periods). B) Predicted abundance across all species; see Supplementary Figure 7. Warmer colors indicate fewer species are predicted to be abundant (i.e., have cells where predicted abundance>1).

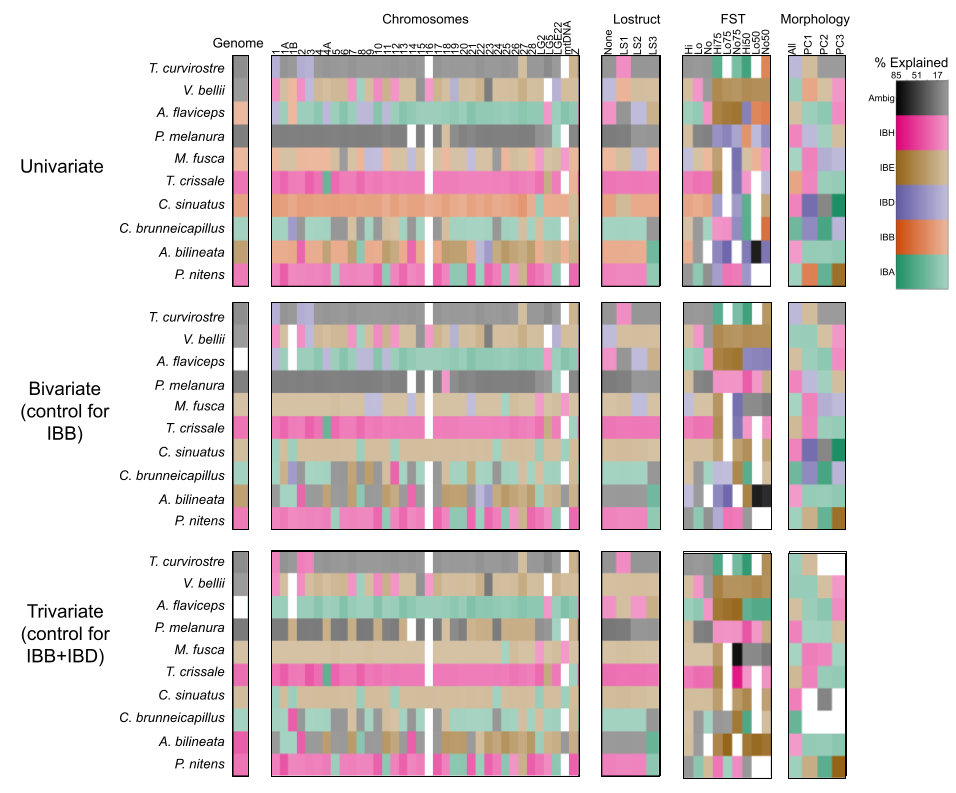

Supplementary Figure 11: GDM model summary for all models. Colors as in Figure 5. Transparency indicates relative percentage explained per model, with more saturated colors having higher percent explained.

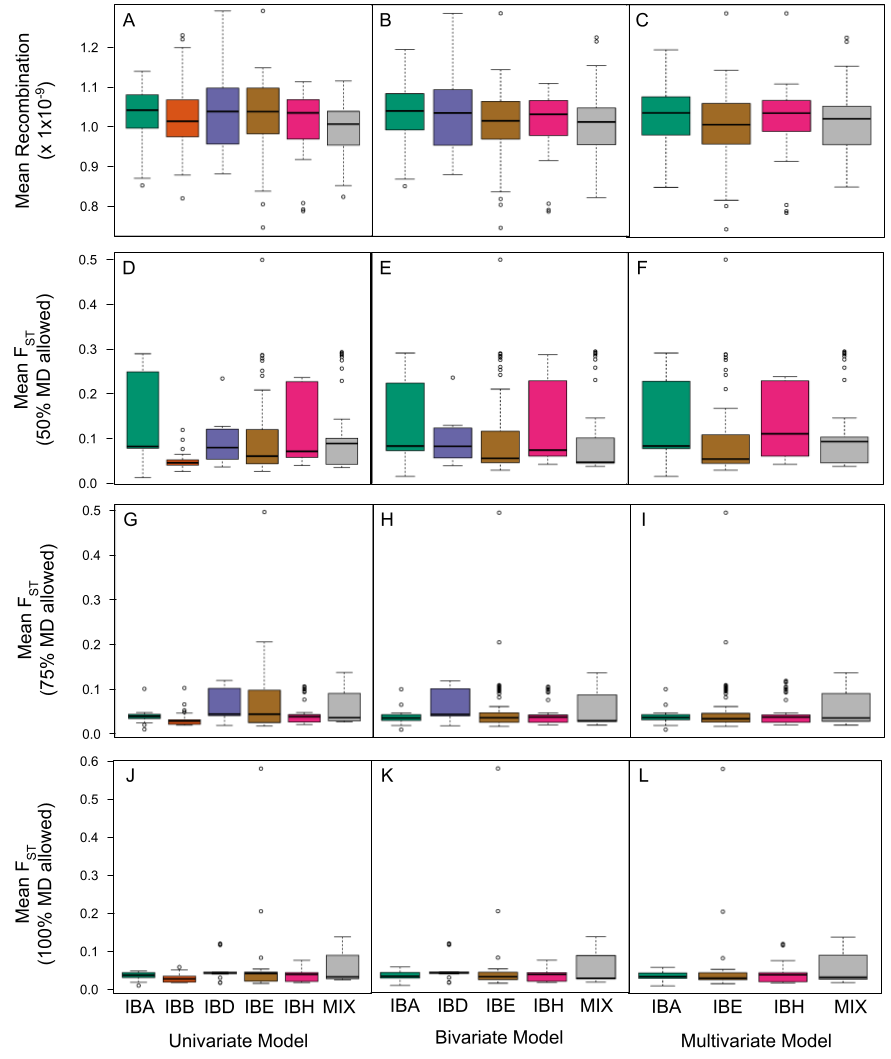

Supplementary Figure 12: D_XY_ and missing data vary across different partitions. Summary statistics along y-axis are as follows: Percent missing data for univariate (A), bivariate (B), and trivariate (C) models. Mean D_XY_ for 50% missing data allowed for univariate (D), bivariate (E), and trivariate (F) models. Mean D_XY_ for 75% missing data allowed for univariate (G), bivariate (H), and trivariate (I) models. Mean D_XY_ for 100% missing data allowed for univariate (J), bivariate (K), and trivariate (L) models. Fill color of box plots correspond to models on the x-axis. Left column x-axis: univariate models.

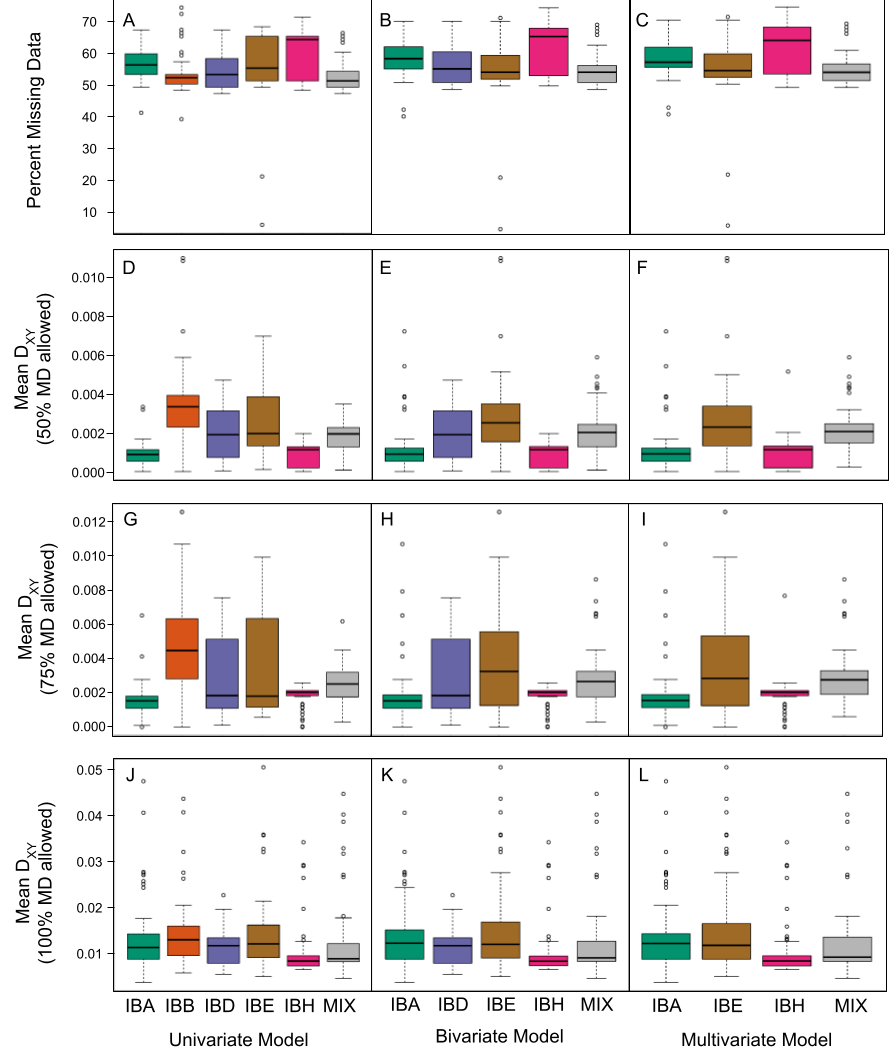

Supplementary Figure 13: F_ST_ and recombination vary across partitions. Summary statistics along y-axis are as follows: Mean recombination rate for univariate (A), bivariate (B), and trivariate (C) models. Mean F_ST_ for 50% missing data allowed for univariate (D), bivariate (E), and trivariate (F) models. Mean F_ST_ for 75% missing data allowed for univariate (G), bivariate (H), and trivariate (I) models. Mean F_ST_ for 100% missing data allowed for univariate (J), bivariate (K), and trivariate (L) models. Fill color of box plots correspond to models on the x-axis.

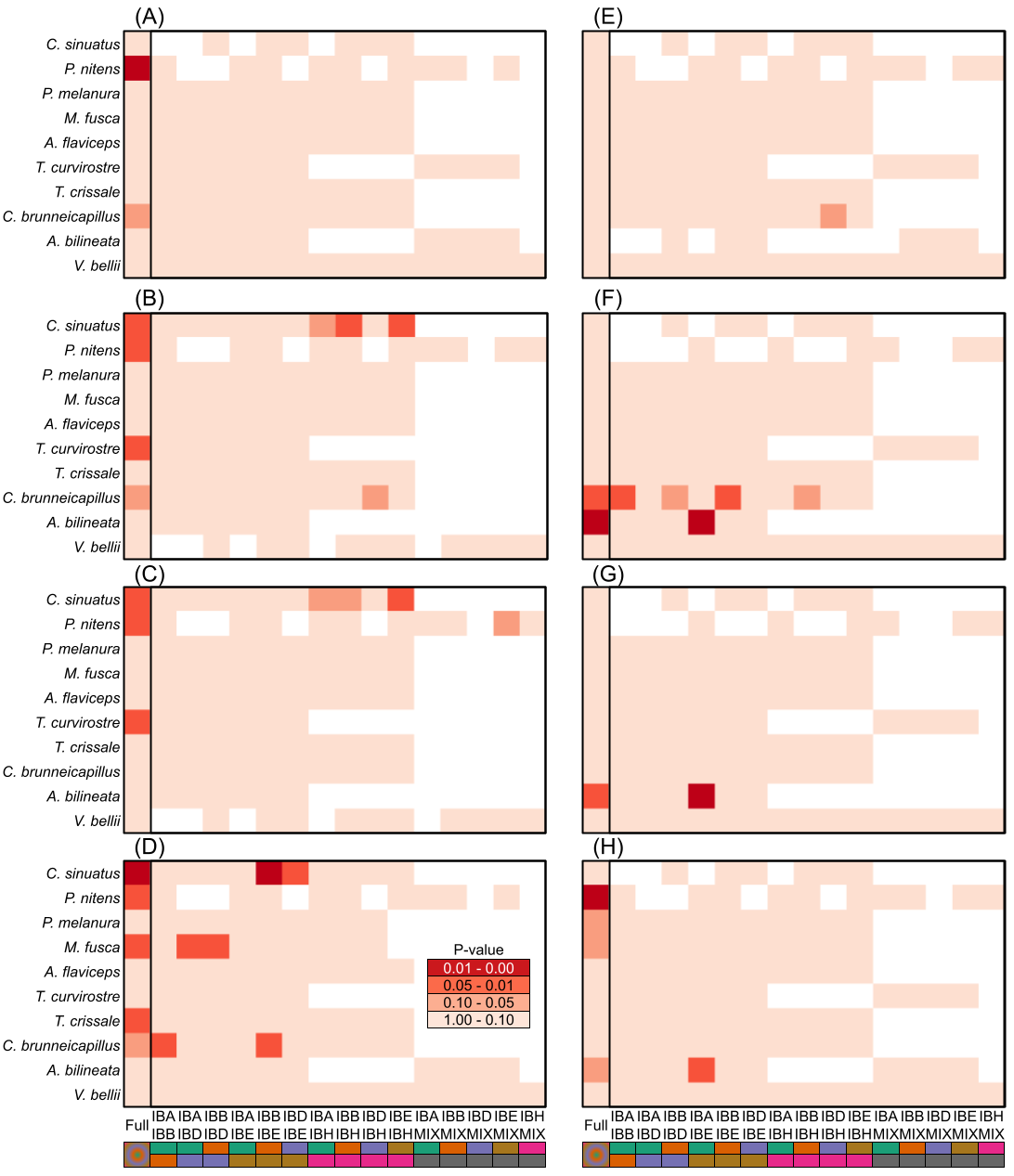

Supplementary Figure 14: Only a few models show significant differences across summary statistics and across species. Analysis of variance tests comparing whether different models of landscape variation vary across different summary statistics for each species. Species are given along the y-axis. Pairwise comparisons of models (IBA, IBB, IBD, IBE, IBH, “MIX” or ambiguous) are shown along the x-axis with colors as in Figure 6. Leftmost model shows the p-value from the overall analysis of variance test (“Full”). Remaining models show p-values from Tukey’s Honest Significant Differences tests on pairwise comparisons between models. Summary statistics are as follows: A) percent missing data; B) D_XY_ for 50% missing data; C) D_XY_ for 75% missing data; D) D_XY_ for 100% missing data; E) recombination rate; F) F_ST_ for 50% missing data; G) F_ST_ for 75% missing data; H) F_ST_ for 100% missing data. Color scale shown in inset of panel D shows darker red values as being more significant than lighter red. White boxes have no associated data due to model types not being present for that species.

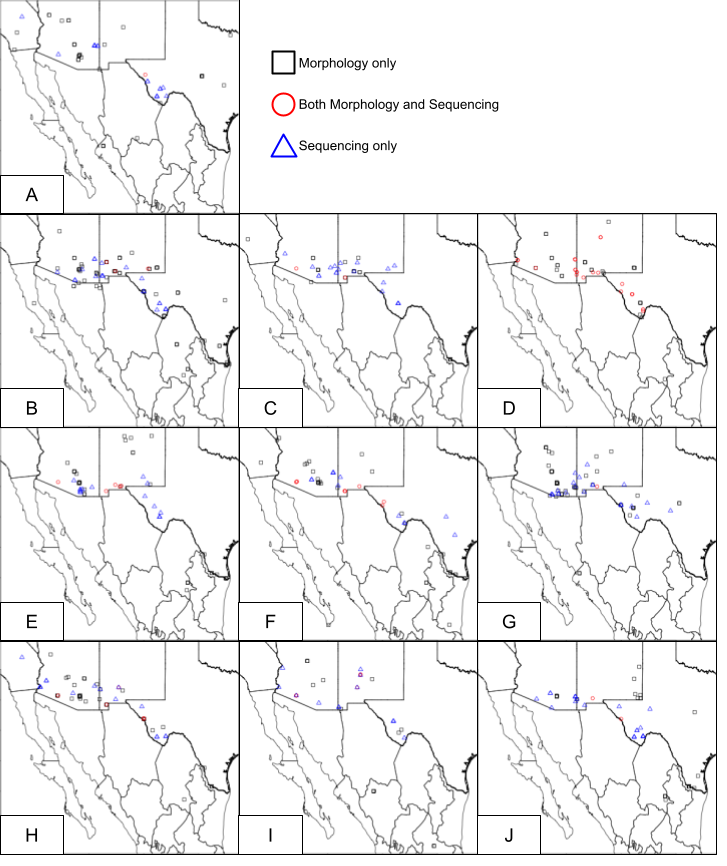

Supplementary Figure 15. Locations and data types of specimens used. Black squares indicate specimens were used for morphological measurements only. Blue triangles indicate specimens were used for sequencing only. Red circles indicate that specimens were used for both morphology and sequencing. Species are as follows: A) *Vireo bellii*, B) *Amphispiza bilineata*, C) *Campylorhynchus brunneicapillus*, D) *Toxostoma crissale,* E) *Toxostoma curvirostre*, F) *Auriparus flaviceps*, G) *Melozone fusca*, H) *Polioptila melanura*, I) *Phainopepla nitens*, J) *Cardinalis sinuatus*.

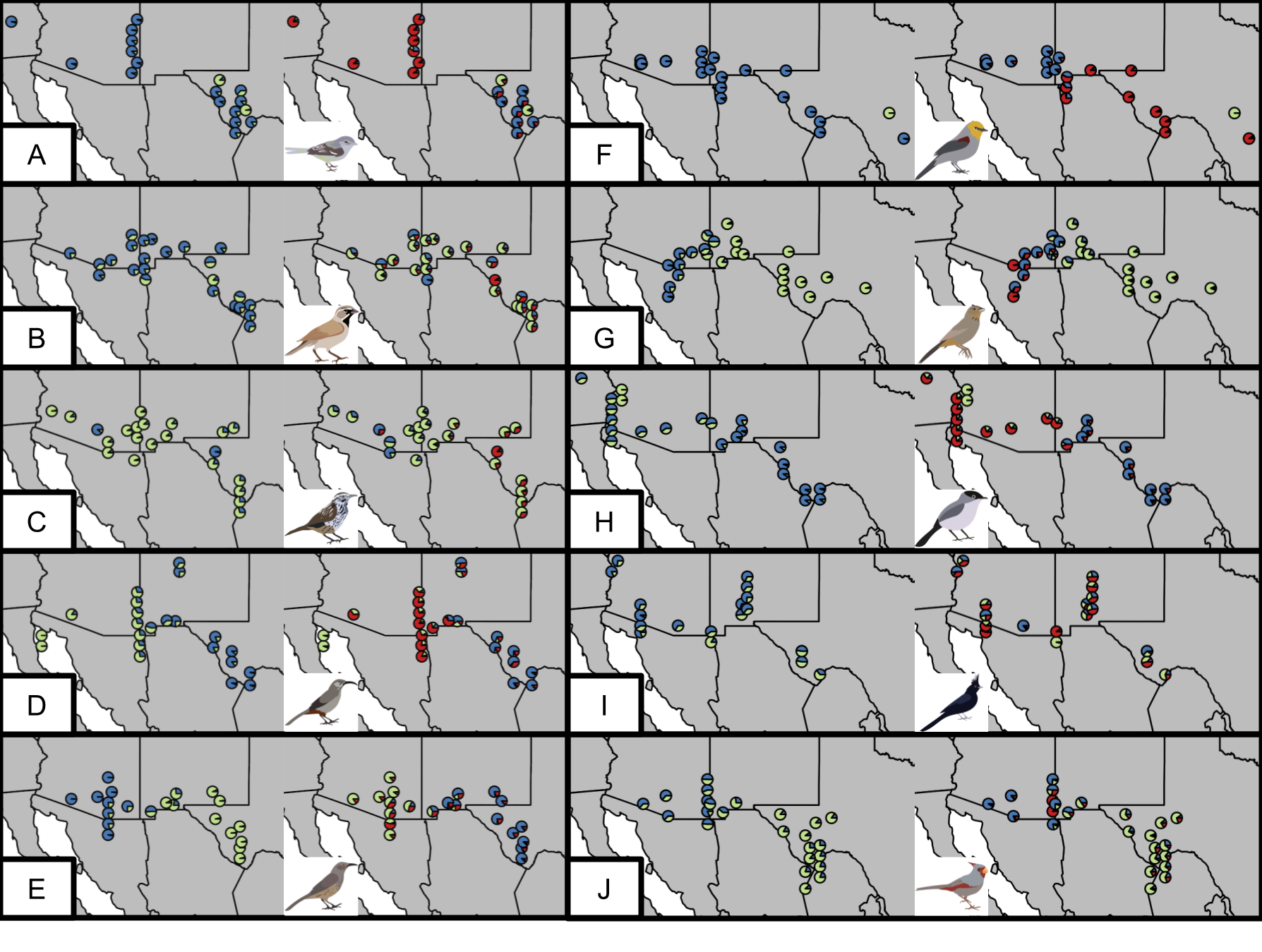

Supplementary Figure 16: PCAngsd results for the 100% missing data allowed partition. Pie charts show individuals with colors giving the proportion of each cluster. Individuals at the same locality are jittered by latitude, though their longitude is kept the same. For each panel, the left side shows K=2 and the right side shows K=3. A) *Vireo bellii*, B) *Amphispiza bilineata*, C) *Campylorhynchus brunneicapillus*, D) *Toxostoma crissale,* E) *Toxostoma curvirostre*, F) *Auriparus flaviceps*, G) *Melozone fusca*, H) *Polioptila melanura*, I) *Phainopepla nitens*, J) *Cardinalis sinuatus*.

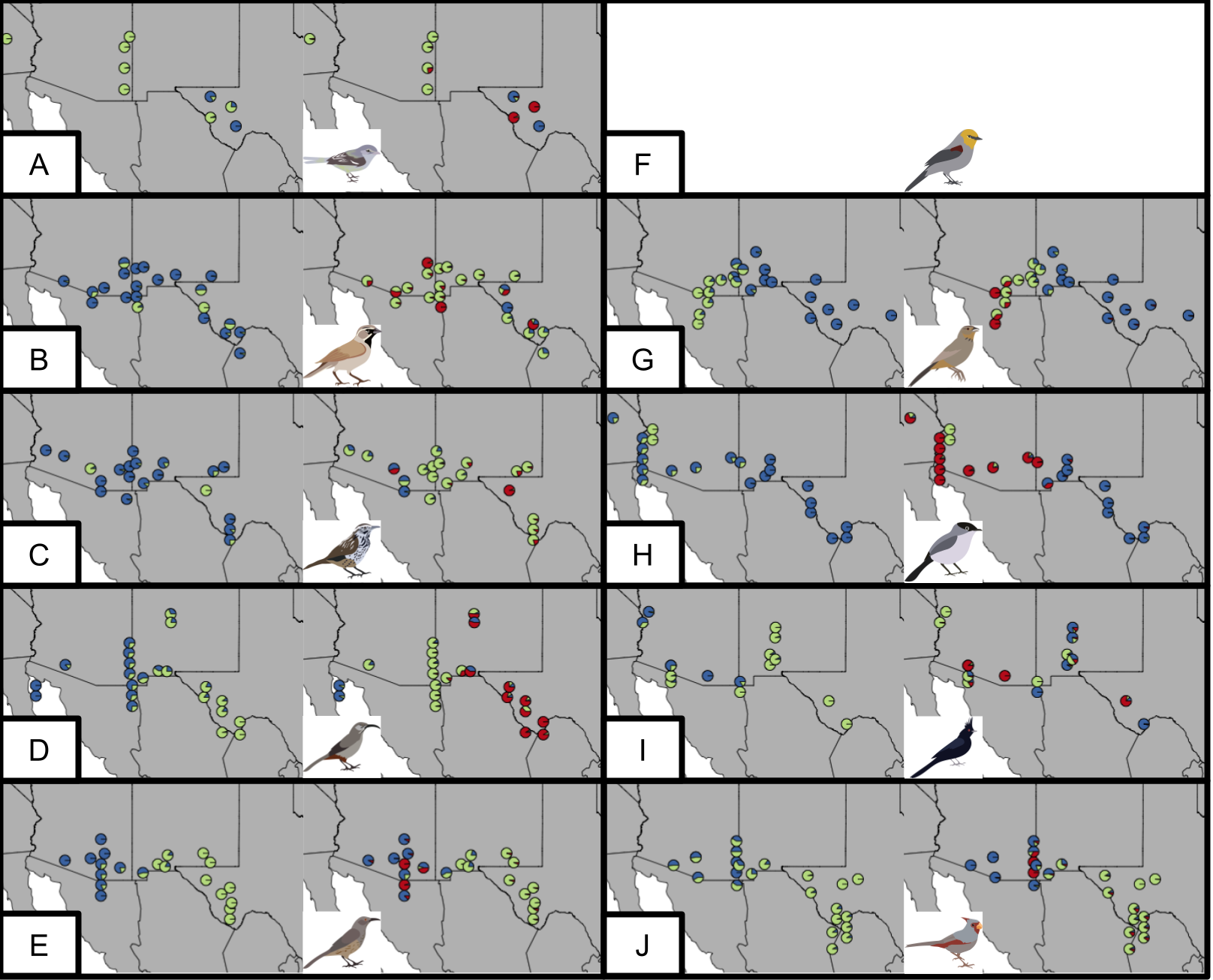

Supplementary Figure 17: PCAngsd results for the 75% missing data allowed partition. Colors and legend as in Supplementary Figure 2. Note that Panel F (*Auriparus flaviceps*) is blank as the analysis failed to converge.

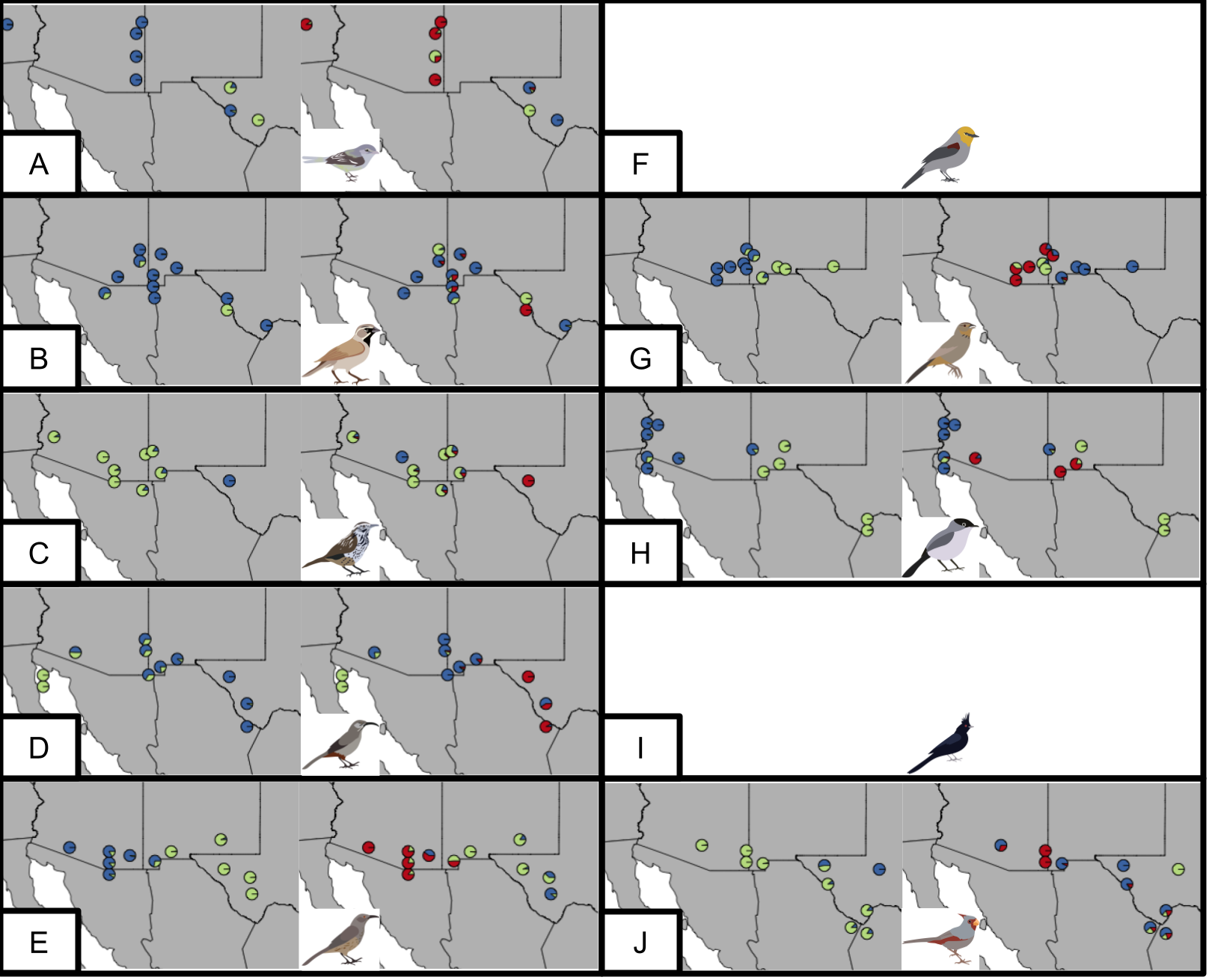

Supplementary Figure 18: PCAngsd results for the 50% missing data allowed partition. Colors and legend as in Supplementary Figure 16 and Supplementary Figure 17. Note that Panels F (*Auriparus flaviceps*) and I (*Phainopepla nitens*) are blank as the analysis failed to converge.

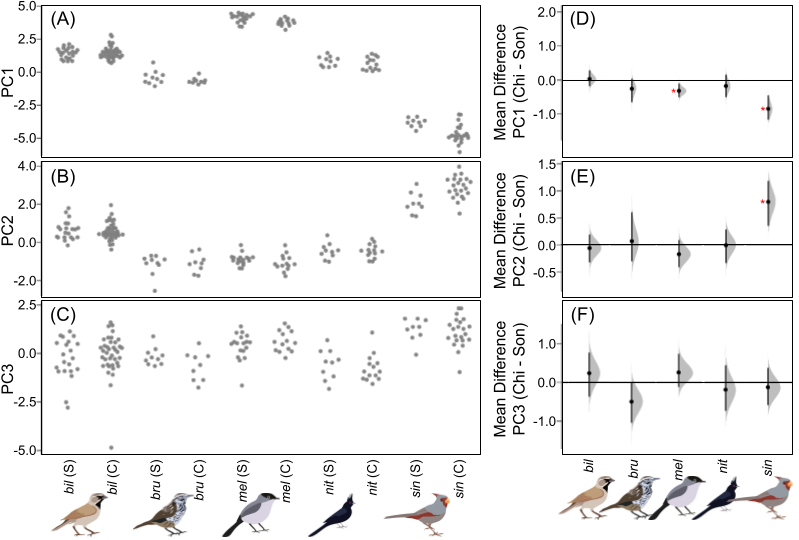

Supplementary Figure 19: Variation across principal component axes of morphology for first five of ten species of birds, with differences across deserts. Left column: raw values for PC1 (A), PC2 (B), and PC3 (C) for each species across each desert. Species abbreviated as follows: “bil”=*Amphispiza bilineata*, “bru”=*Campylorhynchus brunneicapillus*, “mel”=*Polioptila melanura*, “nit”=*Phainopepla nitens*, “sin”=*Cardinalis sinuatus*. Suffix after species designates desert identity: “S”=Sonoran, “C”=Chihuahuan. Points are jittered on the x-axis for visibility. Right column: distribution of unpaired mean differences between Sonoran and Chihuahuan desert individuals for each species from DABEST analysis for PC1 (D), PC2 (E), and PC3 (F). Red asterisk indicates significance.

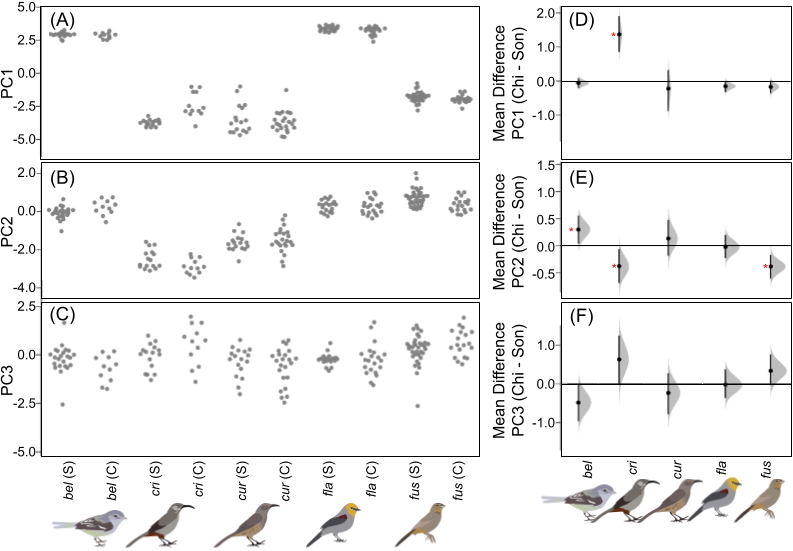

Supplementary Figure 20: Variation across principal component axes of morphology for second five of ten species of birds, with differences across deserts. Left column: raw values for PC1 (A), PC2 (B), and PC3 (C) for each species across each desert. Species abbreviated as follows: “bel”=*Vireo bellii*, “cri”=*Toxostoma crissale*, “cur”=*Toxostoma curvirostre*, “fla”=*Auriparus flaviceps,* “fus”=*Melozone fusca*. Suffix after species designates desert identity: “S”=Sonoran, “C”=Chihuahuan. Points are jittered on the x-axis for visibility. Right column: distribution of unpaired mean differences between Sonoran and Chihuahuan desert individuals for each species from DABEST analysis for PC1 (D), PC2 (E), and PC3 (F). Red asterisk indicates significance.

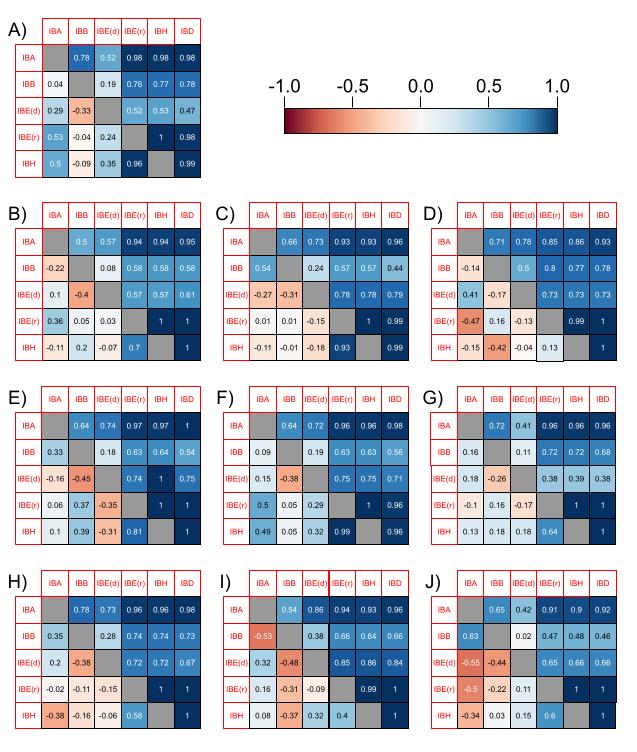

Supplementary Figure 21. Correlation structures within species across matrices used in generalized dissimilarity matrix modeling. Colors indicate correlations and range from -1 (dark red) to 0 (white) to 1 (dark blue). Above the diagonal gives correlations between values before correcting for geographic distances (i.e., IBD). Below the diagonal gives correlations after correcting for geographic distances, as residuals. Note that IBE is decomposed into two measures: IBE(d) which is the environmental distance between points, and IBE(r) which is the environmental resistance between points. Species are as follows: A) *Vireo bellii*, B) *Amphispiza bilineata*, C) *Campylorhynchus brunneicapillus*, D) *Toxostoma crissale,* E) *Toxostoma curvirostre*, F) *Auriparus flaviceps*, G) *Melozone fusca*, H) *Polioptila melanura*, I) *Phainopepla nitens*, J) *Cardinalis sinuatus*.

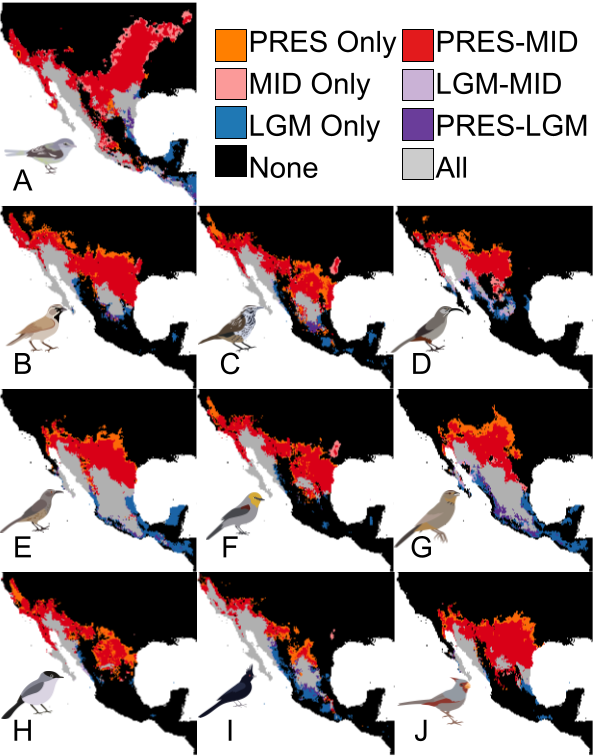

Supplementary Figure 22: Visualization of suitable regions across present (PRES), mid-Holocene (MID), and Last Glacial Maximum (LGM). Colors indicate which regions are suitable at which time periods. A) *Vireo bellii*, B) *Amphispiza bilineata*, C) *Campylorhynchus brunneicapillus*, D) *Toxostoma crissale*, E) *Toxostoma curvirostre*, F) *Auriparus flaviceps*, G) *Melozone fusca*, H) *Polioptila melanura*, I) *Phainopepla nitens*, J) *Cardinalis sinuatus*.

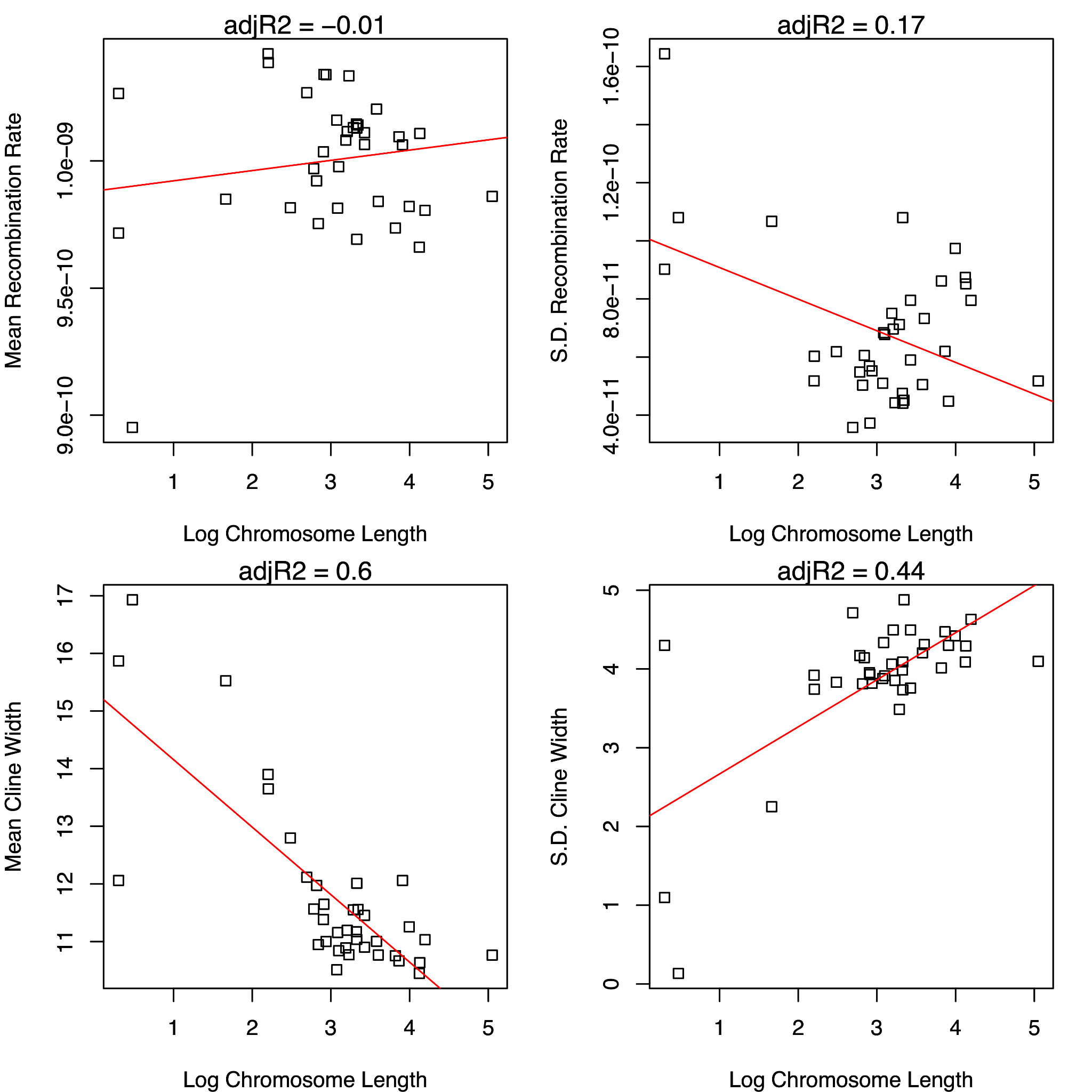

Supplementary Figure 23: Cline metrics (bottom y-axis) and recombination rates (top y-axis) vs log chromosome lengths (x-axis) for each species. Top left: mean recombination rate. Top right: standard deviation (S.D.) recombination rate. Bottom left: mean cline width. Bottom right: S.D.c line width. Line through points show the line of best fit between measurements. Adjusted R^2^ values (adjR2) for lines of best fit are shown above each plot.

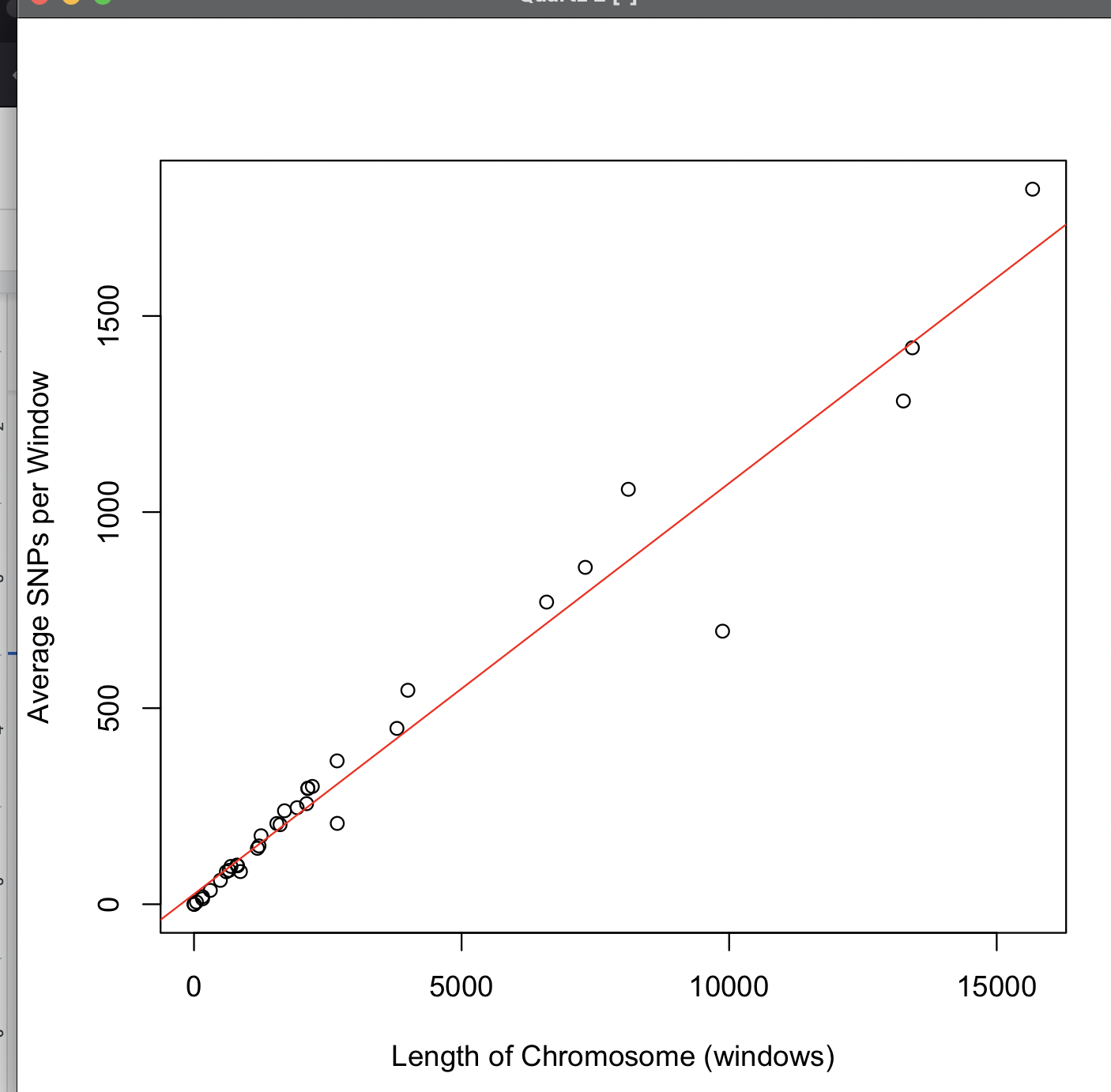

Supplementary Figure 24: Larger chromosomes have more SNPs per window than smaller chromosomes. X-axis shows the length of the chromosome in windows. Y-axis shows the average number of SNPs per 100,000 base pair window. Red line indicates model fit between two measurements (adjusted R^2^ = 0.96, p ~ 0, df = 34).

### Supplementary Tables

Supplementary Table 1: Size-weighted chromosome-wise values for the recombination rate, F_ST_, D_XY_, and proportion of missing data per each species. F_ST_ and D_XY_ are given across the 50% missing, 75% missing, and complete datasets. Values given as mean±standard deviation. These are calculated by weighting chromosomes by number of sites per chromosome. Note that *Campylorhynchus brunneicapillus* and *Phainopepla nitens* have highly inflated F_ST_ values in the 50% dataset due to having only single individuals representing the Chihuahuan desert. Rec”=population recombination rate, or rho.

| **Species** | **Rec (x 10^-10^)** | **F_ST 100_** | **F_ST 75_** | **F_ST 50_** | **D_XY 100_** | **D_XY 75_** | **D_XY 50_** | **Missing Data (%)** |
| --- | --- | --- | --- | --- | --- | --- | --- | --- |
| *Vireo bellii* | 9.0±1.1 | 0.05±0.003 | 0.10±0.004 | 0.13±0.004 | 0.010±0.002 | 0.002±0.002 | 0.002±0.001 | 0.59±0.17 |
| *Amphispiza bilineata* | 11.1±0.04 | 0.02±0.0004 | 0.02±0.0005 | 0.05±0.001 | 0.015±0.004 | 0.007±0.003 | 0.004±0.002 | 0.53±0.02 |
| *Campylorhynchus brunneicapillus* | 10.2±0.3 | 0.03±0.001 | 0.04±0.0009 | 0.29±0.02 | 0.008±0.001 | 0.002±0.0004 | 0.001±0.0002 | 0.54±0.02 |
| *Toxostoma crissale* | 10.5±0.2 | 0.05±0.004 | 0.05±0.006 | 0.07±0.005 | 0.007±0.001 | 0.002±0.0002 | 0.001±0.0001 | 0.50±0.02 |
| *Toxostoma curvirostre* | 10.2±0.4 | 0.12±0.028 | 0.12±0.03 | 0.12±0.02 | 0.009±0.002 | 0.003±0.0005 | 0.002±0.0003 | 0.49±0.02 |
| *Auriparus flaviceps* | 10.1±0.6 | 0.05±0.002 | 0.05±0.002 | 0.09±0.002 | 0.014±0.006 | 0.001±0.001 | 0.001±0.0005 | 0.53±0.02 |
| *Melozone fusca* | 10.2±0.4 | 0.05±0.003 | 0.05±0.003 | 0.06±0.002 | 0.018±0.015 | 0.006±0.006 | 0.004±0.002 | 0.49±0.02 |
| *Polioptila melanura* | 9.4±0.7 | 0.03±0.001 | 0.03±0.0005 | 0.05±0.001 | 0.014±0.009 | 0.002±0.001 | 0.002±0.001 | 0.50±0.03 |
| *Phainopepla nitens* | 9.9±0.5 | 0.02±0.0003 | 0.03±0.0003 | 0.24±0.004 | 0.011±0.005 | 0.002±0.001 | 0.0002±0.001 | 0.65±0.01 |
| *Cardinalis sinuatus* | 10.1±0.5 | 0.029±0.002 | 0.03±0.002 | 0.06±0.004 | 0.012±0.006 | 0.003±0.003 | 0.003±0.002 | 0.52±0.02 |

Supplementary Table 2: Average mean and standard deviation (given as mean±SD) of cline width and cline center location for each species. Species are arranged in order from narrowest to broadest cline width.

| **Species** | **Cline Width** | **Cline Center** |
| --- | --- | --- |
| *Toxostoma curvirostre* | 6.94±1.78 | 7.61±0.59 |
| *Auriparus flaviceps* | 7.51±4.04 | 7.95±1.07 |
| *Melozone fusca* | 7.62±3.12 | 6.55±0.49 |
| *Vireo bellii* | 10.28±3.20 | 9.45±0.96 |
| *Toxostoma crissale* | 10.97±.154 | 7.96±0.90 |
| *Polioptila melanura* | 13.28±1.82 | 3.58±2.81 |
| *Cardinalis sinuatus* | 13.77±1.96 | 7.45±2.72 |
| *Campylorhynchus brunneicapillus* | 15.32±0.52 | 12.70±4.82 |
| *Amphispiza bilineata* | 15.84±0.72 | 11.63±3.93 |
| *Phainopepla nitens* | 15.89±1.20 | 8.91±2.64 |

Supplementary Table 3: Importance of principal components for the morphological analysis.

| **PC** | **Standard Deviation** | **Proportion of Variance** | **Cumulative Proportion** |
| --- | --- | --- | --- |
| 1 | 3.23 | 0.74 | 0.74 |
| 2 | 1.29 | 0.12 | 0.86 |
| 3 | 0.90 | 0.06 | 0.92 |
| 4 | 0.69 | 0.03 | 0.96 |
| 5 | 0.55 | 0.02 | 0.98 |
| 6 | 0.40 | 0.01 | 0.99 |
| 7 | 0.24 | 4.27x10^-3^ | 0.99 |
| 8 | 0.22 | 3.48x10^-3^ | 0.99 |
| 9 | 0.17 | 2.05x10^-3^ | 0.99 |
| 10 | 0.09 | 5.30x10^-4^ | 0.99 |
| 11 | 0.07 | 3.60x10^-4^ | 0.99 |
| 12 | 0.02 | 3.00x10^-5^ | 0.99 |
| 13 | 0.01 | 1.00x10^-5^ | 0.99 |
| 14 | 1.24x10^-10^ | 0 | 1 |

Supplementary Table 4: Loadings of the first three principal components from the raw morphological measurements.

| **Measurement** | **PC1** | **PC2** | **PC3** |
| --- | --- | --- | --- |
| Bill Height | 0.25 | -0.40 | 0.18 |
| Bill Length | 0.25 | 0.41 | -0.19 |
| Bill Width | 0.28 | -0.30 | 0.10 |
| Tarsus Length | 0.27 | 0.26 | -0.10 |
| Primaries Length | 0.29 | 0.10 | -0.18 |
| Secondaries Length | 0.29 | 0.15 | -0.08 |
| Tail Length | 0.28 | 0.21 | -0.03 |
| Kipp’s Index | -0.10 | -0.39 | -0.85 |
| Beak Base Area | 0.24 | -0.42 | 0.26 |
| Beak Volume | 0.29 | -0.03 | 0.09 |
| Beak Lateral Surface Area | 0.29 | 0.19 | -0.06 |
| Beak Full Surface Area | 0.30 | -0.07 | 0.08 |
| Beak Lateral Surface / Volume | -0.26 | 0.24 | 0.15 |
| Beak Full Surface / Volume | -0.28 | 0.05 | 0.21 |

Supplementary Table 5: Specimens used across this study. Catalog numbers are given; for catalog numbers with “*” after them, the catalog number is provisional and is awaiting final assignment. “Seq.” stands for whether specimens were used for sequencing. “Morph” stands for whether specimens were morphologically measured. “Lat.” and “Long.” stand for latitude and longitude, respectively.

| **Catalog** | **Seq.** | **Morph.** | **Species** | **Lat.** | **Long.** |
| --- | --- | --- | --- | --- | --- |
| AMNH 225551* | Yes | No | *Vireo bellii* | 32.90633 | -109.48584 |
| AMNH 225582* | Yes | No | *Vireo bellii* | 32.90633 | -109.48584 |
| AMNH 244216* | Yes | No | *Vireo bellii* | 29.56316 | -103.80576 |
| AMNH 244217* | Yes | No | *Vireo bellii* | 29.56316 | -103.80576 |
| AMNH 244223* | Yes | No | *Vireo bellii* | 29.56316 | -103.80576 |
| AMNH 252976* | Yes | No | *Vireo bellii* | 32.90633 | -109.48584 |
| AMNH 252989* | Yes | No | *Vireo bellii* | 32.888 | -109.49718 |
| AMNH J-0018* | Yes | No | *Vireo bellii* | 32.32576 | -112.7888 |
| AMNH J-0127* | Yes | Yes | *Vireo bellii* | 30.99453 | -104.86922 |
| AMNH SKIN 027978 | No | Yes | *Vireo bellii* | 32.98 | -111.33 |
| AMNH SKIN 027981 | No | Yes | *Vireo bellii* | 32.98 | -111.33 |
| AMNH SKIN 027982 | No | Yes | *Vireo bellii* | 32.98 | -111.33 |
| AMNH SKIN 053342 | No | Yes | *Vireo bellii* | 34.56 | -111.85 |
| AMNH SKIN 378975 | No | Yes | *Vireo bellii* | 30.3 | -97.7 |
| AMNH SKIN 378976 | No | Yes | *Vireo bellii* | 30.9 | -99.7 |
| AMNH SKIN 378977 | No | Yes | *Vireo bellii* | 30.9 | -99.7 |
| AMNH SKIN 378979 | No | Yes | *Vireo bellii* | 30.9 | -99.7 |
| AMNH SKIN 378980 | No | Yes | *Vireo bellii* | 30.9 | -99.7 |
| AMNH SKIN 378981 | No | Yes | *Vireo bellii* | 30.9 | -99.7 |
| AMNH SKIN 378982 | No | Yes | *Vireo bellii* | 30.9 | -99.7 |
| AMNH SKIN 378991 | No | Yes | *Vireo bellii* | 32.26 | -110.87 |
| AMNH SKIN 758926 | No | Yes | *Vireo bellii* | 33.6 | -97.1 |
| AMNH SKIN 758927 | No | Yes | *Vireo bellii* | 30.9 | -99.7 |
| CU 15650 | No | Yes | *Vireo bellii* | 29.13 | -103.51 |
| DMNH 23085 | No | Yes | *Vireo bellii* | 26.82 | -105.61 |
| DMNH 26108 | No | Yes | *Vireo bellii* | 32.14 | -110.69 |
| DMNH 32263 | No | Yes | *Vireo bellii* | 34.92 | -113.63 |
| DMNH 32264 | No | Yes | *Vireo bellii* | 34.92 | -113.63 |
| DMNH 32266 | No | Yes | *Vireo bellii* | 31.35 | -109.05 |
| DMNH 32267 | No | Yes | *Vireo bellii* | 32.32 | -110.83 |
| DMNH 32268 | No | Yes | *Vireo bellii* | 32.26 | -110.87 |
| DMNH 50282 | No | Yes | *Vireo bellii* | 31.97 | -110.96 |
| DMNH 50283 | No | Yes | *Vireo bellii* | 31.97 | -110.96 |
| DMNH 7812 | No | Yes | *Vireo bellii* | 32.64 | -116.78 |
| DMNH 7813 | No | Yes | *Vireo bellii* | 33.77 | -116.56 |
| LSUMZ 3912 | Yes | No | *Vireo bellii* | 34.8 | -116.1 |
| MSB SKIN 14356 | No | Yes | *Vireo bellii* | 32.97 | -108.58 |
| MSB SKIN 2414 | No | Yes | *Vireo bellii* | 34.12 | -106.88 |
| TCWC 16642 H114 | Yes | No | *Vireo bellii* | 30.13548 | -103.239761 |
| TCWC 24033 | Yes | No | *Vireo bellii* | 29.57829 | -102.93075 |
| TCWC 24288 | Yes | No | *Vireo bellii* | 30.547861 | -104.66106 |
| TCWC 24311 | Yes | No | *Vireo bellii* | 30.55033 | -104.66553 |
| TCWC 24506 | Yes | No | *Vireo bellii* | 30.05721 | -103.50431 |
| TCWC 24507 | Yes | No | *Vireo bellii* | 30.05721 | -103.50431 |
| UMMZ SKIN 163424 | No | Yes | *Vireo bellii* | 27.41 | -112.51 |
| UMMZ SKIN 55195 | No | Yes | *Vireo bellii* | 32.22 | -110.93 |
| UMMZ SKIN 60309 | No | Yes | *Vireo bellii* | 34.56 | -111.85 |
| UMMZ SKIN 60310 | No | Yes | *Vireo bellii* | 34.54 | -112.47 |
| UWBM 77850 | Yes | No | *Vireo bellii* | 32.87 | -109.2 |
| UWBM 77855 | Yes | No | *Vireo bellii* | 32.87 | -109.2 |
| UWBM 81294 | No | Yes | *Vireo bellii* | 26.304 | -108.699 |
| UWBM 88987 | No | Yes | *Vireo bellii* | 26.3155 | -108.6989 |
| UWBM 88988 | No | Yes | *Vireo bellii* | 26.3155 | -108.6989 |
| AMNH 225569* | Yes | No | *Amphispiza bilineata* | 32.90515 | -109.49018 |
| AMNH 225589* | Yes | No | *Amphispiza bilineata* | 32.90515 | -109.49018 |
| AMNH 243820* | No | Yes | *Amphispiza bilineata* | 32.99 | -107.2 |
| AMNH 243821* | No | Yes | *Amphispiza bilineata* | 32.99 | -107.2 |
| AMNH 244184* | Yes | No | *Amphispiza bilineata* | 30.775478 | -105.01974 |
| AMNH 244241* | Yes | No | *Amphispiza bilineata* | 29.56316 | -103.80576 |
| AMNH 244306* | Yes | No | *Amphispiza bilineata* | 30.775478 | -105.01974 |
| AMNH J-0010* | No | Yes | *Amphispiza bilineata* | 32.29 | -112.86 |
| AMNH J-0020* | Yes | Yes | *Amphispiza bilineata* | 32.70953 | -108.39813 |
| AMNH J-0021* | No | Yes | *Amphispiza bilineata* | 32.71 | -108.4 |
| AMNH J-0022* | No | Yes | *Amphispiza bilineata* | 32.71 | -108.4 |
| AMNH J-0023* | No | Yes | *Amphispiza bilineata* | 32.71 | -108.4 |
| AMNH J-0025* | No | Yes | *Amphispiza bilineata* | 31.71 | -108.39 |
| AMNH J-0054* | Yes | Yes | *Amphispiza bilineata* | 32.10705 | -107.60438 |
| AMNH J-0067* | No | Yes | *Amphispiza bilineata* | 32.11 | -107.68 |
| AMNH J-0068* | No | Yes | *Amphispiza bilineata* | 32.11 | -107.68 |
| AMNH J-0069* | No | Yes | *Amphispiza bilineata* | 32.11 | -107.6 |
| AMNH J-0071* | No | Yes | *Amphispiza bilineata* | 32.11 | -107.6 |
| AMNH J-0092* | No | Yes | *Amphispiza bilineata* | 31.99 | -107.2 |
| AMNH J-0093* | No | Yes | *Amphispiza bilineata* | 31.99 | -107.2 |
| AMNH J-0101* | No | Yes | *Amphispiza bilineata* | 32.99 | -107.2 |
| AMNH J-0130* | No | Yes | *Amphispiza bilineata* | 30.8 | -105.01 |
| AMNH J-0131* | No | Yes | *Amphispiza bilineata* | 30.8 | -105.01 |
| AMNH J-0140* | No | Yes | *Amphispiza bilineata* | 30.75 | -104.97 |
| AMNH J-0143* | No | Yes | *Amphispiza bilineata* | 30.75 | -104.97 |
| AMNH J-0984* | Yes | No | *Amphispiza bilineata* | 32.64092 | -108.83216 |
| AMNH SKIN 028378 | No | Yes | *Amphispiza bilineata* | 32.98 | -111.33 |
| AMNH SKIN 028385 | No | Yes | *Amphispiza bilineata* | 32.98 | -111.33 |
| AMNH SKIN 028397 | No | Yes | *Amphispiza bilineata* | 32.42 | -110.63 |
| AMNH SKIN 056356 | No | Yes | *Amphispiza bilineata* | 31 | -109 |
| AMNH SKIN 083861 | No | Yes | *Amphispiza bilineata* | 26.3 | -98.8 |
| AMNH SKIN 083862 | No | Yes | *Amphispiza bilineata* | 26.3 | -98.8 |
| AMNH SKIN 083894 | No | Yes | *Amphispiza bilineata* | 30.7 | -101.9 |
| AMNH SKIN 401809 | No | Yes | *Amphispiza bilineata* | 25.95 | -97.51 |
| AMNH SKIN 401811 | No | Yes | *Amphispiza bilineata* | 27.2 | -98.1 |
| AMNH SKIN 401812 | No | Yes | *Amphispiza bilineata* | 27.2 | -98.1 |
| AMNH SKIN 401840 | No | Yes | *Amphispiza bilineata* | 32.2 | -110.87 |
| AMNH SKIN 401865 | No | Yes | *Amphispiza bilineata* | 27.53 | -99.48 |
| AMNH SKIN 518495 | No | Yes | *Amphispiza bilineata* | 30.7 | -101.9 |
| AMNH SKIN 518501 | No | Yes | *Amphispiza bilineata* | 30.7 | -101.9 |
| AMNH SKIN 762584 | No | Yes | *Amphispiza bilineata* | 30.6 | -103.89 |
| AMNH SKIN 762591 | No | Yes | *Amphispiza bilineata* | 27.3 | -102 |
| AMNH SKIN 762598 | No | Yes | *Amphispiza bilineata* | 30 | -115 |
| AMNH SKIN 762607 | No | Yes | *Amphispiza bilineata* | 32.26 | -110.87 |
| AMNH SKIN 767122 | No | Yes | *Amphispiza bilineata* | 30.2 | -97.6 |
| AMNH SKIN 808103 | No | Yes | *Amphispiza bilineata* | 27.3 | -102 |
| AMNH SKIN 808104 | No | Yes | *Amphispiza bilineata* | 27.3 | -102 |
| AMNH SKIN 836849 | No | Yes | *Amphispiza bilineata* | 31.9 | -109.2 |
| DMNH 13222 | No | Yes | *Amphispiza bilineata* | 25.67 | -101 |
| DMNH 17379 | No | Yes | *Amphispiza bilineata* | 32.02 | -109.33 |
| DMNH 17380 | No | Yes | *Amphispiza bilineata* | 31.13 | -109.32 |
| DMNH 23604 | No | Yes | *Amphispiza bilineata* | 25.67 | -101 |
| DMNH 45328 | No | Yes | *Amphispiza bilineata* | 31.34 | -110.93 |
| DMNH 50984 | No | Yes | *Amphispiza bilineata* | 25.27 | -101.37 |
| DMNH 50993 | No | Yes | *Amphispiza bilineata* | 26.13 | -97.4 |
| DMNH 50994 | No | Yes | *Amphispiza bilineata* | 26 | -97.2 |
| DMNH 50996 | No | Yes | *Amphispiza bilineata* | 32.12 | -110.81 |
| DMNH 8790 | No | Yes | *Amphispiza bilineata* | 25.99 | -97.45 |
| DMNH 8796 | No | Yes | *Amphispiza bilineata* | 32.22 | -110.93 |
| DMNH 8797 | No | Yes | *Amphispiza bilineata* | 32.56 | -110.68 |
| DMNH 8798 | No | Yes | *Amphispiza bilineata* | 31.96 | -109.31 |
| LSUMZ 36269 | Yes | No | *Amphispiza bilineata* | 31.6227 | -105.1811 |
| MSB 21396 | Yes | No | *Amphispiza bilineata* | 29.60194444 | -103.0013889 |
| MSB 21438 | Yes | No | *Amphispiza bilineata* | 29.60194444 | -103.0013889 |
| MSB 24372 | Yes | No | *Amphispiza bilineata* | 32.0417302 | -112.8754369 |
| MSB 24591 | Yes | No | *Amphispiza bilineata* | 29.986988 | -103.562886 |
| MSB 24596 | Yes | No | *Amphispiza bilineata* | 29.986988 | -103.562886 |
| MSB 28940 | Yes | No | *Amphispiza bilineata* | 32.34503333 | -106.60295 |
| MSB 29306 | Yes | Yes | *Amphispiza bilineata* | 32.28165 | -104.6047833 |
| MSB SKIN 16241 | No | Yes | *Amphispiza bilineata* | 33.32 | -106.08 |
| MSB SKIN 29308 | No | Yes | *Amphispiza bilineata* | 32.26 | -104.49 |
| MSB SKIN 29316 | No | Yes | *Amphispiza bilineata* | 32.26 | -104.49 |
| MSB SKIN 4574 | No | Yes | *Amphispiza bilineata* | 32.35 | -103.79 |
| MSB SKIN 5186 | No | Yes | *Amphispiza bilineata* | 32.36 | -103.78 |
| MSB SKIN 5187 | No | Yes | *Amphispiza bilineata* | 32.34 | -103.84 |
| UMMZ SKIN 3948 | No | Yes | *Amphispiza bilineata* | 31.13 | -111.38 |
| UMMZ SKIN 4278 | No | Yes | *Amphispiza bilineata* | 31.13 | -111.38 |
| UMMZ SKIN 55404 | No | Yes | *Amphispiza bilineata* | 34.74 | -112.68 |
| UMMZ SKIN 56857 | No | Yes | *Amphispiza bilineata* | 32.22 | -110.93 |
| UMMZ SKIN 57262 | No | Yes | *Amphispiza bilineata* | 31.49 | -110.41 |
| UMMZ SKIN 59879 | No | Yes | *Amphispiza bilineata* | 29.17 | -103.15 |
| UMMZ SKIN 59884 | No | Yes | *Amphispiza bilineata* | 29.17 | -103.15 |
| UMMZ SKIN 59886 | No | Yes | *Amphispiza bilineata* | 29.17 | -103.15 |
| UMMZ SKIN 59888 | No | Yes | *Amphispiza bilineata* | 29.17 | -103.15 |
| UMMZ SKIN 60426 | No | Yes | *Amphispiza bilineata* | 25.9 | -97.5 |
| UWBM 111929 | Yes | No | *Amphispiza bilineata* | 31.76027778 | -109.3769444 |
| UWBM 114894 | Yes | No | *Amphispiza bilineata* | 29.60194444 | -103.0013889 |
| UWBM 77531 | Yes | No | *Amphispiza bilineata* | 31.7972 | -108.737 |
| UWBM 77557 | Yes | No | *Amphispiza bilineata* | 31.5282 | -111.2677 |
| UWBM 77636 | Yes | No | *Amphispiza bilineata* | 31.7972 | -108.804 |
| UWBM 77657 | Yes | No | *Amphispiza bilineata* | 31.5087 | -111.2825 |
| UWBM 77693 | Yes | No | *Amphispiza bilineata* | 31.7972 | -108.804 |
| UWBM 77822 | Yes | No | *Amphispiza bilineata* | 32.22 | -110.62 |
| AMNH 244261* | Yes | No | *Campylorhynchus brunneicapillus* | 30.775478 | -105.01974 |
| AMNH J-0009* | Yes | Yes | *Campylorhynchus brunneicapillus* | 32.29166 | -112.85538 |
| AMNH J-0024* | No | Yes | *Campylorhynchus brunneicapillus* | 32.71 | -108.4 |
| AMNH J-0030* | No | Yes | *Campylorhynchus brunneicapillus* | 31.69 | -108.4 |
| AMNH J-0034* | Yes | Yes | *Campylorhynchus brunneicapillus* | 31.6921 | -108.39689 |
| AMNH J-0053* | No | Yes | *Campylorhynchus brunneicapillus* | 32.11 | -107.6 |
| AMNH J-0060* | No | Yes | *Campylorhynchus brunneicapillus* | 32.11 | -107.60438. |
| AMNH J-0063* | No | Yes | *Campylorhynchus brunneicapillus* | 32.11 | -107.6 |
| AMNH J-0072* | No | Yes | *Campylorhynchus brunneicapillus* | 32.11 | -107.6 |
| AMNH J-0094* | No | Yes | *Campylorhynchus brunneicapillus* | 32.99 | -107.2 |
| AMNH J-0115* | No | Yes | *Campylorhynchus brunneicapillus* | 32.05 | -107.07 |
| AMNH J-0118* | No | Yes | *Campylorhynchus brunneicapillus* | 32.05 | -107.07 |
| AMNH J-0972* | Yes | No | *Campylorhynchus brunneicapillus* | 32.43859 | -109.34566 |
| AMNH J-0987* | Yes | No | *Campylorhynchus brunneicapillus* | 32.64092 | -108.83216 |
| AMNH SKIN 027592 | No | Yes | *Campylorhynchus brunneicapillus* | 32.98 | -111.33 |
| AMNH SKIN 027613 | No | Yes | *Campylorhynchus brunneicapillus* | 32.98 | -111.33 |
| AMNH SKIN 085866 | No | Yes | *Campylorhynchus brunneicapillus* | 26.3 | -98.8 |
| AMNH SKIN 085876 | No | Yes | *Campylorhynchus brunneicapillus* | 26.3 | -98.8 |
| AMNH SKIN 085877 | No | Yes | *Campylorhynchus brunneicapillus* | 26.3 | -98.8 |
| AMNH SKIN 085878 | No | Yes | *Campylorhynchus brunneicapillus* | 26.3 | -98.8 |
| AMNH SKIN 085882 | No | Yes | *Campylorhynchus brunneicapillus* | 26.3 | -98.8 |
| AMNH SKIN 326476 | No | Yes | *Campylorhynchus brunneicapillus* | 31.4 | -111.5 |
| AMNH SKIN 374940 | No | Yes | *Campylorhynchus brunneicapillus* | 32.2 | -110.87 |
| AMNH SKIN 374941 | No | Yes | *Campylorhynchus brunneicapillus* | 32.2 | -110.87 |
| AMNH SKIN 374942 | No | Yes | *Campylorhynchus brunneicapillus* | 32.2 | -110.87 |
| LSUMZ 20006 | Yes | No | *Campylorhynchus brunneicapillus* | 32.385287 | -111.373898 |
| LSUMZ 62493 | Yes | No | *Campylorhynchus brunneicapillus* | 31.348064 | -104.902636 |
| MSB 22000 | Yes | No | *Campylorhynchus brunneicapillus* | 32.6653557 | -107.2903195 |
| MSB 24490 | Yes | No | *Campylorhynchus brunneicapillus* | 29.986988 | -103.562886 |
| MSB 24527 | Yes | No | *Campylorhynchus brunneicapillus* | 29.986988 | -103.562886 |
| MSB 24627 | Yes | No | *Campylorhynchus brunneicapillus* | 29.986988 | -103.562886 |
| MSB 24629 | Yes | No | *Campylorhynchus brunneicapillus* | 29.986988 | -103.562886 |
| MSB 25299 | Yes | No | *Campylorhynchus brunneicapillus* | 32.4648399 | -103.8988304 |
| MSB 26352 | Yes | No | *Campylorhynchus brunneicapillus* | 32.21638333 | -108.9914 |
| MSB 29018 | Yes | No | *Campylorhynchus brunneicapillus* | 33.260584 | -113.883224 |
| MSB 29302 | Yes | No | *Campylorhynchus brunneicapillus* | 32.26156667 | -104.4916 |
| MSB 39823 | Yes | No | *Campylorhynchus brunneicapillus* | 32.08943333 | -107.5912167 |
| UWBM 77526 | Yes | No | *Campylorhynchus brunneicapillus* | 31.7994 | -110.7942 |
| UWBM 77602 | Yes | No | *Campylorhynchus brunneicapillus* | 31.7994 | -110.7942 |
| UWBM 77884 | Yes | No | *Campylorhynchus brunneicapillus* | 32.34 | -109.74 |
| UWBM 77990 | Yes | No | *Campylorhynchus brunneicapillus* | 32 | -109.2 |
| UWBM 77991 | Yes | No | *Campylorhynchus brunneicapillus* | 32 | -109.2 |
| UWBM 90714 | No | Yes | *Campylorhynchus brunneicapillus* | 34.215539 | -117.269592 |
| AMNH 243826 | No | Yes | *Toxostoma crissale* | 32.33 | -112.79 |
| AMNH 243826* | Yes | No | *Toxostoma crissale* | 32.32576 | -112.7888 |
| AMNH DOT-3738 | Yes | No | *Toxostoma crissale* | 31.872954 | -109.06196 |
| AMNH J-0107* | Yes | No | *Toxostoma crissale* | 32.01089 | -107.09655 |
| AMNH J-1015* | Yes | No | *Toxostoma crissale* | 31.69971 | -108.43874 |
| AMNH J-1023* | Yes | No | *Toxostoma crissale* | 32.03875 | -107.56973 |
| AMNH LJM-110* | Yes | No | *Toxostoma crissale* | 30.775478 | -105.01974 |
| AMNH SKIN 053651 | No | Yes | *Toxostoma crissale* | 32.2 | -107.7 |
| AMNH SKIN 053652 | No | Yes | *Toxostoma crissale* | 32.2 | -107.7 |
| AMNH SKIN 053654 | No | Yes | *Toxostoma crissale* | 34.56 | -111.85 |
| AMNH SKIN 053658 | No | Yes | *Toxostoma crissale* | 34.56 | -111.85 |
| AMNH SKIN 078292 | No | Yes | *Toxostoma crissale* | 32.1 | -106.7 |
| AMNH SKIN 078297 | No | Yes | *Toxostoma crissale* | 32 | -106.6 |
| AMNH SKIN 085855 | No | Yes | *Toxostoma crissale* | 32.5 | -110.8 |
| AMNH SKIN 085856 | No | Yes | *Toxostoma crissale* | 32.98 | -111.33 |
| AMNH SKIN 085857 | No | Yes | *Toxostoma crissale* | 32.98 | -111.33 |
| AMNH SKIN 085859 | No | Yes | *Toxostoma crissale* | 32.5 | -110.8 |
| AMNH SKIN 406445 | No | Yes | *Toxostoma crissale* | 31.8 | -110.4 |
| AMNH SKIN 406446 | No | Yes | *Toxostoma crissale* | 31.8 | -110.4 |
| AMNH SKIN 406456 | No | Yes | *Toxostoma crissale* | 32.2 | -110.87 |
| AMNH SKIN 446844 | No | Yes | *Toxostoma crissale* | 32.7 | -108.3 |
| AMNH SKIN 446845 | No | Yes | *Toxostoma crissale* | 32.7 | -108.3 |
| AMNH SKIN 446846 | No | Yes | *Toxostoma crissale* | 32.7 | -108.3 |
| AMNH SKIN 758127 | No | Yes | *Toxostoma crissale* | 32.6 | -114.6 |
| CU 13631 | No | Yes | *Toxostoma crissale* | 30.04 | -103.27 |
| CU 13635 | No | Yes | *Toxostoma crissale* | 30.02 | -103.24 |
| CU 13636 | No | Yes | *Toxostoma crissale* | 30.03 | -103.24 |
| DMNH 49780 | No | Yes | *Toxostoma crissale* | 25.02 | -101.02 |
| LSUMZ 16582 | Yes | No | *Toxostoma crissale* | 32.207 | -109.1802 |
| LSUMZ 16586 16596? | Yes | No | *Toxostoma crissale* | 32.207 | -109.1802 |
| LSUMZ 40753 | Yes | No | *Toxostoma crissale* | 30.6 | -104 |
| LSUMZ 47304 | Yes | No | *Toxostoma crissale* | 30.6 | -104 |
| LSUMZ 47509 | Yes | No | *Toxostoma crissale* | 32.0526078 | -109.03965 |
| LSUMZ 47510 | Yes | No | *Toxostoma crissale* | 32.0526078 | -109.03965 |
| LSUMZ 51732 | Yes | No | *Toxostoma crissale* | 29.5 | -103 |
| LSUMZ 58267 | Yes | No | *Toxostoma crissale* | 30.6 | -104 |
| MSB 20926 | Yes | No | *Toxostoma crissale* | 34.3500633 | -106.8839111 |
| MSB 21308 | Yes | No | *Toxostoma crissale* | 29.60194444 | -103.0013889 |
| MSB 26421 | Yes | No | *Toxostoma crissale* | 34.3420895 | -106.868906 |
| MSB SKIN 21023 | No | Yes | *Toxostoma crissale* | 35.36 | -106.15 |
| MSB SKIN 4564 | No | Yes | *Toxostoma crissale* | 32.33 | -103.77 |
| MSB SKIN 5141 | No | Yes | *Toxostoma crissale* | 32.34 | -103.84 |
| MSB SKIN 5142 | No | Yes | *Toxostoma crissale* | 32.34 | -103.83 |
| TCWC 23990 | Yes | No | *Toxostoma crissale* | 31.27104 | -104.90847 |
| UMMZ SKIN 59771 | No | Yes | *Toxostoma crissale* | 29.25 | -103.28 |
| UMMZ SKIN 86184 | No | Yes | *Toxostoma crissale* | 29.09 | -103.09 |
| UWBM 109226 | Yes | No | *Toxostoma crissale* | 32.8395 | -114.4415 |
| UWBM 115562 | Yes | No | *Toxostoma crissale* | 32.8395 | -114.4415 |
| UWBM 77981 | Yes | No | *Toxostoma crissale* | 32.9 | -109.22 |
| UWBM 77984 | Yes | No | *Toxostoma crissale* | 32.87 | -109.19 |
| AMNH 243824* | Yes | Yes | *Toxostoma curvirostre* | 32.10705 | -107.60438 |
| AMNH J-0003* | Yes | Yes | *Toxostoma curvirostre* | 32.28772 | -112.81215 |
| AMNH J-0050* | Yes | Yes | *Toxostoma curvirostre* | 31.69971 | -108.43874 |
| AMNH J-0089* | Yes | Yes | *Toxostoma curvirostre* | 31.9918 | -107.19848 |
| AMNH J-0109* | No | Yes | *Toxostoma curvirostre* | 32.01 | -107.1 |
| AMNH J-0126* | Yes | Yes | *Toxostoma curvirostre* | 32.03581 | -107.05341 |
| AMNH SKIN 027710 | No | Yes | *Toxostoma curvirostre* | 32.98 | -111.33 |
| AMNH SKIN 027711 | No | Yes | *Toxostoma curvirostre* | 32.98 | -111.33 |
| AMNH SKIN 027721 | No | Yes | *Toxostoma curvirostre* | 32.98 | -111.33 |
| AMNH SKIN 027727 | No | Yes | *Toxostoma curvirostre* | 32.98 | -111.33 |
| AMNH SKIN 085811 | No | Yes | *Toxostoma curvirostre* | 26.3 | -98.8 |
| AMNH SKIN 085812 | No | Yes | *Toxostoma curvirostre* | 26.3 | -98.8 |
| AMNH SKIN 085823 | No | Yes | *Toxostoma curvirostre* | 26.21 | -98.33 |
| AMNH SKIN 085829 | No | Yes | *Toxostoma curvirostre* | 27.53 | -99.48 |
| AMNH SKIN 085847 | No | Yes | *Toxostoma curvirostre* | 32.98 | -111.33 |
| AMNH SKIN 375766 | No | Yes | *Toxostoma curvirostre* | 32.2 | -110.87 |
| AMNH SKIN 375784 | No | Yes | *Toxostoma curvirostre* | 31.8 | -110.4 |
| AMNH SKIN 375786 | No | Yes | *Toxostoma curvirostre* | 25.95 | -97.51 |
| AMNH SKIN 375788 | No | Yes | *Toxostoma curvirostre* | 25.95 | -97.51 |
| AMNH SKIN 375789 | No | Yes | *Toxostoma curvirostre* | 25.95 | -97.51 |
| AMNH SKIN 375790 | No | Yes | *Toxostoma curvirostre* | 25.95 | -97.51 |
| AMNH SKIN 375800 | No | Yes | *Toxostoma curvirostre* | 25.95 | -97.51 |
| AMNH SKIN 758067 | No | Yes | *Toxostoma curvirostre* | 31.49 | -110.41 |
| DMNH 12741 | No | Yes | *Toxostoma curvirostre* | 25.67 | -101 |
| DMNH 12742 | No | Yes | *Toxostoma curvirostre* | 25.67 | -101 |
| DMNH 12743 | No | Yes | *Toxostoma curvirostre* | 25.67 | -101 |
| DMNH 22895 | No | Yes | *Toxostoma curvirostre* | 25.18 | -101.08 |
| DMNH 35977 | No | Yes | *Toxostoma curvirostre* | 33.55 | -111.95 |
| DMNH 46088 | No | Yes | *Toxostoma curvirostre* | 32.22 | -110.93 |
| DMNH 7474 | No | Yes | *Toxostoma curvirostre* | 32.26 | -110.87 |
| DMNH 7475 | No | Yes | *Toxostoma curvirostre* | 32.26 | -110.87 |
| DMNH 7477 | No | Yes | *Toxostoma curvirostre* | 32.26 | -110.87 |
| DMNH 7478 | No | Yes | *Toxostoma curvirostre* | 32.26 | -110.87 |
| LSUMZ 16593 18593? | Yes | No | *Toxostoma curvirostre* | 32.385287 | -111.373898 |
| LSUMZ 18961 | Yes | No | *Toxostoma curvirostre* | 31.7994 | -110.7942 |
| LSUMZ 58304 | Yes | No | *Toxostoma curvirostre* | 31.348064 | -104.902636 |
| LSUMZ 9910 | Yes | No | *Toxostoma curvirostre* | 31.7994 | -110.7942 |
| MSB 21019 | Yes | No | *Toxostoma curvirostre* | 30.266021 | -103.467831 |
| MSB 24262 | Yes | No | *Toxostoma curvirostre* | 32.18956 | -104.6612549 |
| MSB 24488 | Yes | No | *Toxostoma curvirostre* | 29.986988 | -103.562886 |
| MSB 24489 | Yes | No | *Toxostoma curvirostre* | 29.986988 | -103.562886 |
| MSB 24616 | Yes | No | *Toxostoma curvirostre* | 29.986988 | -103.562886 |
| MSB 24783 | Yes | No | *Toxostoma curvirostre* | 31.813625 | -110.5904091 |
| MSB 29272 | Yes | No | *Toxostoma curvirostre* | 30.684167 | -104.03808 |
| MSB SKIN 19093 | No | Yes | *Toxostoma curvirostre* | 32.14 | -104.43 |
| MSB SKIN 23226 | No | Yes | *Toxostoma curvirostre* | 35.04 | -106.67 |
| MSB SKIN 24894 | No | Yes | *Toxostoma curvirostre* | 35.3 | -106.77 |
| MSB SKIN 3149 | No | Yes | *Toxostoma curvirostre* | 34.23 | -105.99 |
| MSB SKIN 3153 | No | Yes | *Toxostoma curvirostre* | 34.22 | -105.99 |
| MSB SKIN 3509 | No | Yes | *Toxostoma curvirostre* | 35.32 | -104.3 |
| MSB SKIN 3510 | No | Yes | *Toxostoma curvirostre* | 35.19 | -106.17 |
| UMMZ SKIN 280 | No | Yes | *Toxostoma curvirostre* | 32.83 | -111.13 |
| UMMZ SKIN 55373 | No | Yes | *Toxostoma curvirostre* | 31.49 | -110.41 |
| UWBM 77610 | Yes | No | *Toxostoma curvirostre* | 32.63 | -105.059 |
| UWBM 90204 | Yes | No | *Toxostoma curvirostre* | 31.940441 | -109.722118 |
| UWBM 95955 | Yes | No | *Toxostoma curvirostre* | 31.71666667 | -110.7833333 |
| UWBM 95956 | Yes | No | *Toxostoma curvirostre* | 31.61666667 | -110.7833333 |
| UWBM 95957 | Yes | No | *Toxostoma curvirostre* | 31.61666667 | -110.7833333 |
| AMNH 252983* | Yes | No | *Auriparus flaviceps* | 32.90633 | -109.48584 |
| AMNH 252988* | Yes | No | *Auriparus flaviceps* | 32.90633 | -109.48584 |
| AMNH J-0007* | Yes | Yes | *Auriparus flaviceps* | 32.29166 | -112.85538 |
| AMNH J-0016* | Yes | Yes | *Auriparus flaviceps* | 32.29166 | -112.85538 |
| AMNH J-0019* | Yes | Yes | *Auriparus flaviceps* | 32.32576 | -112.7888 |
| AMNH J-0028* | Yes | Yes | *Auriparus flaviceps* | 31.6921 | -108.39689 |
| AMNH J-0037* | Yes | Yes | *Auriparus flaviceps* | 31.69971 | -108.43874 |
| AMNH J-0038* | Yes | Yes | *Auriparus flaviceps* | 31.69971 | -108.43874 |
| AMNH J-0108* | Yes | Yes | *Auriparus flaviceps* | 32.01089 | -107.09655 |
| AMNH J-0128* | Yes | Yes | *Auriparus flaviceps* | 30.99453 | -104.86922 |
| AMNH J-0153* | Yes | Yes | *Auriparus flaviceps* | 30.7517 | -105.00391 |
| AMNH J-0965* | Yes | No | *Auriparus flaviceps* | 32.441 | -111.45826 |
| AMNH J-0967* | Yes | No | *Auriparus flaviceps* | 32.441 | -111.45826 |
| AMNH J-0968* | Yes | No | *Auriparus flaviceps* | 32.441 | -111.45826 |
| AMNH J-0995* | Yes | No | *Auriparus flaviceps* | 32.6049 | -108.83727 |
| AMNH KLP-51* | Yes | No | *Auriparus flaviceps* | 32.88479 | -109.50211 |
| AMNH SKIN 027422 | No | Yes | *Auriparus flaviceps* | 32.98 | -111.33 |
| AMNH SKIN 086444 | No | Yes | *Auriparus flaviceps* | 26.37 | -98.82 |
| AMNH SKIN 086447 | No | Yes | *Auriparus flaviceps* | 26.3 | -98.8 |
| AMNH SKIN 086475 | No | Yes | *Auriparus flaviceps* | 26.21 | -98.33 |
| AMNH SKIN 086476 | No | Yes | *Auriparus flaviceps* | 26.21 | -98.33 |
| AMNH SKIN 086492 | No | Yes | *Auriparus flaviceps* | 32.48 | -110.9 |
| AMNH SKIN 086496 | No | Yes | *Auriparus flaviceps* | 22.9 | -98.9 |
| AMNH SKIN 373566 | No | Yes | *Auriparus flaviceps* | 32.2 | -110.8 |
| AMNH SKIN 373569 | No | Yes | *Auriparus flaviceps* | 32.26 | -110.87 |
| AMNH SKIN 373584 | No | Yes | *Auriparus flaviceps* | 25.95 | -97.51 |
| AMNH SKIN 439066 | No | Yes | *Auriparus flaviceps* | 25.95 | -97.51 |
| AMNH SKIN 505626 | No | Yes | *Auriparus flaviceps* | 27.53 | -99.48 |
| AMNH SKIN 708827 | No | Yes | *Auriparus flaviceps* | 32.2 | -110.6 |
| AMNH SKIN 757255 | No | Yes | *Auriparus flaviceps* | 32.26 | -110.87 |
| AMNH SKIN 757263 | No | Yes | *Auriparus flaviceps* | 32.26 | -110.87 |
| AMNH SKIN 768782 | No | Yes | *Auriparus flaviceps* | 32.3 | -110.7 |
| CU 12475 | No | Yes | *Auriparus flaviceps* | 30.21 | -103.24 |
| DMNH 12583 | No | Yes | *Auriparus flaviceps* | 25.67 | -101 |
| DMNH 1424 | No | Yes | *Auriparus flaviceps* | 31.91 | -109.09 |
| DMNH 22761 | No | Yes | *Auriparus flaviceps* | 25.67 | -101 |
| DMNH 29529 | No | Yes | *Auriparus flaviceps* | 33.64 | -111.67 |
| DMNH 29531 | No | Yes | *Auriparus flaviceps* | 35.18 | -114.42 |
| DMNH 29534 | No | Yes | *Auriparus flaviceps* | 33.45 | -112.07 |
| DMNH 29537 | No | Yes | *Auriparus flaviceps* | 35.05 | -114.62 |
| DMNH 46072 | No | Yes | *Auriparus flaviceps* | 33.51 | -111.9 |
| DMNH 51693 | No | Yes | *Auriparus flaviceps* | 28.02 | -100.86 |
| DMNH 7240 | No | Yes | *Auriparus flaviceps* | 32.26 | -110.87 |
| LSUMZ 15579 | Yes | No | *Auriparus flaviceps* | 30 | -99.2 |
| LSUMZ 8436 | Yes | No | *Auriparus flaviceps* | 28.8 | -98.4 |
| MSB 22493 | Yes | No | *Auriparus flaviceps* | 32.0526078 | -109.03965 |
| MSB SKIN 2413 | No | Yes | *Auriparus flaviceps* | 34.12 | -106.88 |
| MSB SKIN 4603 | No | Yes | *Auriparus flaviceps* | 32.98 | -105.93 |
| TCWC 23805 | Yes | No | *Auriparus flaviceps* | 30.057154 | -103.504181 |
| UMMZ SKIN 160181 | No | Yes | *Auriparus flaviceps* | 33.57 | -116.08 |
| UMMZ SKIN 95667 | No | Yes | *Auriparus flaviceps* | 22.88 | -109.92 |
| UWBM 111637 | Yes | No | *Auriparus flaviceps* | 29.59455 | -103.000783 |
| UWBM 112147 | Yes | No | *Auriparus flaviceps* | 29.5945 | -103.000833 |
| UWBM 41248 | No | Yes | *Auriparus flaviceps* | 29.56343 | -102.89763 |
| AMNH 244178* | Yes | No | *Melozone fusca* | 30.775478 | -105.01974 |
| AMNH 244191* | Yes | No | *Melozone fusca* | 30.775478 | -105.01974 |
| AMNH 244195* | Yes | No | *Melozone fusca* | 30.775478 | -105.01974 |
| AMNH 252997* | No | Yes | *Melozone fusca* | 32.11 | -107.6 |
| AMNH 253043* | No | Yes | *Melozone fusca* | 32.99 | -107.2 |
| AMNH J-0035* | No | Yes | *Melozone fusca* | 31.69 | -108.4 |
| AMNH J-0098* | Yes | Yes | *Melozone fusca* | 31.9918 | -107.19848 |
| AMNH J-0971* | Yes | No | *Melozone fusca* | 32.23907 | -109.33829 |
| AMNH J-0992* | Yes | No | *Melozone fusca* | 32.59244 | -108.81808 |
| AMNH J-1005* | Yes | No | *Melozone fusca* | 31.6239 | -108.3727 |
| AMNH SKIN 028207 | No | Yes | *Melozone fusca* | 32.98 | -111.33 |
| AMNH SKIN 028211 | No | Yes | *Melozone fusca* | 32.98 | -111.33 |
| AMNH SKIN 028213 | No | Yes | *Melozone fusca* | 32.98 | -111.33 |
| AMNH SKIN 028215 | No | Yes | *Melozone fusca* | 34.3 | -111.6 |
| AMNH SKIN 028218 | No | Yes | *Melozone fusca* | 32.98 | -111.33 |
| AMNH SKIN 028219 | No | Yes | *Melozone fusca* | 32.98 | -111.33 |
| AMNH SKIN 028223 | No | Yes | *Melozone fusca* | 32.98 | -111.33 |
| AMNH SKIN 028230 | No | Yes | *Melozone fusca* | 32.98 | -111.33 |
| AMNH SKIN 028235 | No | Yes | *Melozone fusca* | 32.98 | -111.33 |
| AMNH SKIN 028549 | No | Yes | *Melozone fusca* | 32.98 | -111.33 |
| AMNH SKIN 041634 | No | Yes | *Melozone fusca* | 32.7 | -108.2 |
| AMNH SKIN 041636 | No | Yes | *Melozone fusca* | 32.26 | -110.87 |
| AMNH SKIN 041637 | No | Yes | *Melozone fusca* | 34.3 | -105.9 |
| AMNH SKIN 053038 | No | Yes | *Melozone fusca* | 34.56 | -111.85 |
| AMNH SKIN 053039 | No | Yes | *Melozone fusca* | 34.56 | -111.85 |
| AMNH SKIN 053042 | No | Yes | *Melozone fusca* | 34.56 | -111.85 |
| AMNH SKIN 053043 | No | Yes | *Melozone fusca* | 34.56 | -111.85 |
| AMNH SKIN 053044 | No | Yes | *Melozone fusca* | 34.2 | -111.3 |
| AMNH SKIN 056313 | No | Yes | *Melozone fusca* | 31.43 | -109.9 |
| AMNH SKIN 056858 | No | Yes | *Melozone fusca* | 32.5 | -117 |
| AMNH SKIN 084206 | No | Yes | *Melozone fusca* | 30.6 | -103.89 |
| AMNH SKIN 084207 | No | Yes | *Melozone fusca* | 30.6 | -103.89 |
| AMNH SKIN 084208 | No | Yes | *Melozone fusca* | 30.6 | -103.89 |
| AMNH SKIN 084209 | No | Yes | *Melozone fusca* | 30.6 | -103.89 |
| AMNH SKIN 084213 | No | Yes | *Melozone fusca* | 30.6 | -103.89 |
| AMNH SKIN 084214 | No | Yes | *Melozone fusca* | 30.6 | -103.89 |
| AMNH SKIN 084217 | No | Yes | *Melozone fusca* | 31.4 | -103.5 |
| AMNH SKIN 084218 | No | Yes | *Melozone fusca* | 30.1 | -104.2 |
| AMNH SKIN 098979 | No | Yes | *Melozone fusca* | 32.2 | -110.87 |
| AMNH SKIN 326500 | No | Yes | *Melozone fusca* | 31.4 | -111.5 |
| AMNH SKIN 368649 | No | Yes | *Melozone fusca* | 31.49 | -110.41 |
| AMNH SKIN 368650 | No | Yes | *Melozone fusca* | 31.49 | -110.41 |
| AMNH SKIN 368666 | No | Yes | *Melozone fusca* | 32.2 | -110.87 |
| AMNH SKIN 368667 | No | Yes | *Melozone fusca* | 32.2 | -110.87 |
| AMNH SKIN 368686 | No | Yes | *Melozone fusca* | 30.6 | -103.89 |
| AMNH SKIN 368687 | No | Yes | *Melozone fusca* | 30.6 | -103.89 |
| AMNH SKIN 368690 | No | Yes | *Melozone fusca* | 30.9 | -99.7 |
| AMNH SKIN 368691 | No | Yes | *Melozone fusca* | 30.9 | -99.7 |
| AMNH SKIN 461707 | No | Yes | *Melozone fusca* | 31.95 | -109.18 |
| AMNH SKIN 761997 | No | Yes | *Melozone fusca* | 31.49 | -110.41 |
| AMNH SKIN 808359 | No | Yes | *Melozone fusca* | 32.2 | -110.8 |
| AMNH SKIN 842071 | No | Yes | *Melozone fusca* | 30.78 | -105.02 |
| AMNH SKIN 842073 | No | Yes | *Melozone fusca* | 30.78 | -105.02 |
| CU 30208 | No | Yes | *Melozone fusca* | 32.26 | -110.88 |
| DMNH 1427 | No | Yes | *Melozone fusca* | 31.91 | -109.09 |
| DMNH 45192 | No | Yes | *Melozone fusca* | 31.34 | -110.93 |
| DMNH 48948 | No | Yes | *Melozone fusca* | 31.71 | -110.07 |
| DMNH 48949 | No | Yes | *Melozone fusca* | 31.72 | -110.74 |
| DMNH 8762 | No | Yes | *Melozone fusca* | 31.96 | -109.31 |
| LSUMZ 16732 | Yes | No | *Melozone fusca* | 30.4 | -100.5 |
| LSUMZ 78587 | Yes | No | *Melozone fusca* | 30.684167 | -104.03808 |
| MSB 24608 | Yes | No | *Melozone fusca* | 29.986988 | -103.562886 |
| MSB 29301 | Yes | No | *Melozone fusca* | 32.1115 | -104.72955 |
| MSB 39814 | Yes | No | *Melozone fusca* | 32.08933333 | -107.5911333 |
| MSB 39815 | Yes | No | *Melozone fusca* | 32.08933333 | -107.5911333 |
| MSB 41397 | Yes | No | *Melozone fusca* | 33.42735 | -108.08727 |
| MSB SKIN 26936 | No | Yes | *Melozone fusca* | 33.97 | -107.25 |
| TCWC 15842 TJH2704 | Yes | No | *Melozone fusca* | 30.9054194 | -102.6154611 |
| UMMZ SKIN 55043 | No | Yes | *Melozone fusca* | 31.93 | -109.22 |
| UMMZ SKIN 55382 | No | Yes | *Melozone fusca* | 31.49 | -110.41 |
| UWBM 48497 | No | Yes | *Melozone fusca* | 33.37 | -111.62 |
| UWBM 77790 | Yes | No | *Melozone fusca* | 31.45 | -111.25 |
| UWBM 77870 | Yes | No | *Melozone fusca* | 31.5275 | -110.71069 |
| UWBM 77873 | Yes | No | *Melozone fusca* | 31.5087484 | -110.7202464 |
| UWBM 77951 | Yes | No | *Melozone fusca* | 31.59 | -110.04 |
| UWBM 77992 | Yes | No | *Melozone fusca* | 32 | -109.2 |
| UWBM 78059 | Yes | No | *Melozone fusca* | 32.87 | -109.19 |
| UWBM 82734 | No | Yes | *Melozone fusca* | 26.275 | -108.795 |
| UWBM 84066 | No | Yes | *Melozone fusca* | 26.31 | -108.81 |
| UWBM 95906 | Yes | No | *Melozone fusca* | 31.47666667 | -111.3355556 |
| UWBM 95907 | Yes | No | *Melozone fusca* | 31.47666667 | -111.3355556 |
| UWBM 95909 | Yes | No | *Melozone fusca* | 31.71666667 | -110.7833333 |
| AMNH 244242* | Yes | No | *Polioptila melanura* | 29.56316 | -103.80576 |
| AMNH 253008* | No | Yes | *Polioptila melanura* | 30.73 | -104.99 |
| AMNH 292985* | Yes | No | *Polioptila melanura* | 32.9094 | -109.49009 |
| AMNH J-0001* | Yes | Yes | *Polioptila melanura* | 32.28772 | -112.81215 |
| AMNH J-0005* | No | Yes | *Polioptila melanura* | 32.29 | -112.86 |
| AMNH J-0006* | No | Yes | *Polioptila melanura* | 32.29 | -112.86 |
| AMNH J-0008* | No | Yes | *Polioptila melanura* | 32.29 | -112.86 |
| AMNH J-0014* | No | Yes | *Polioptila melanura* | 32.29 | -112.86 |
| AMNH J-0026* | Yes | Yes | *Polioptila melanura* | 31.6921 | -108.39689 |
| AMNH J-0031* | No | Yes | *Polioptila melanura* | 31.69 | -108.4 |
| AMNH J-0032* | No | Yes | *Polioptila melanura* | 31.69 | -108.4 |
| AMNH J-0033A* | No | Yes | *Polioptila melanura* | 31.69 | -108.4 |
| AMNH J-0129* | Yes | Yes | *Polioptila melanura* | 30.797 | -105.01138 |
| AMNH J-0139* | Yes | Yes | *Polioptila melanura* | 30.7517 | -105.00391 |
| AMNH J-0144* | No | Yes | *Polioptila melanura* | 30.75 | -104.97 |
| AMNH J-0150* | No | Yes | *Polioptila melanura* | 30.75 | -105 |
| AMNH J-0151* | No | Yes | *Polioptila melanura* | 30.75 | -105 |
| AMNH J-0963* | Yes | No | *Polioptila melanura* | 32.441 | -111.45826 |
| AMNH J-1020* | Yes | No | *Polioptila melanura* | 32.03875 | -107.56973 |
| AMNH SKIN 039349 | No | Yes | *Polioptila melanura* | 32.26 | -110.87 |
| AMNH SKIN 086584 | No | Yes | *Polioptila melanura* | 26.3 | -98.8 |
| AMNH SKIN 086585 | No | Yes | *Polioptila melanura* | 26.3 | -98.8 |
| AMNH SKIN 086586 | No | Yes | *Polioptila melanura* | 26.3 | -98.8 |
| AMNH SKIN 086596 | No | Yes | *Polioptila melanura* | 30 | -104.2 |
| AMNH SKIN 086602 | No | Yes | *Polioptila melanura* | 27.53 | -99.48 |
| AMNH SKIN 086609 | No | Yes | *Polioptila melanura* | 32.98 | -111.33 |
| AMNH SKIN 377584 | No | Yes | *Polioptila melanura* | 31.9 | -110.3 |
| AMNH SKIN 377595 | No | Yes | *Polioptila melanura* | 32.2 | -110.87 |
| AMNH SKIN 808213 | No | Yes | *Polioptila melanura* | 32.3 | -110.8 |
| AMNH WMMIII-385* | Yes | No | *Polioptila melanura* | 29.56316 | -103.80576 |
| CU 13987 | No | Yes | *Polioptila melanura* | 30.21 | -103.24 |
| DMNH 30293 | No | Yes | *Polioptila melanura* | 34.99 | -113.49 |
| DMNH 30298 | No | Yes | *Polioptila melanura* | 32.08 | -110.79 |
| DMNH 30299 | No | Yes | *Polioptila melanura* | 32.22 | -110.79 |
| DMNH 30304 | No | Yes | *Polioptila melanura* | 33.4 | -110.39 |
| DMNH 30305 | No | Yes | *Polioptila melanura* | 33.03 | -109.27 |
| DMNH 30307 | No | Yes | *Polioptila melanura* | 32.08 | -111.56 |
| DMNH 30310 | No | Yes | *Polioptila melanura* | 32.26 | -110.87 |
| DMNH 30313 | No | Yes | *Polioptila melanura* | 33.42 | -110.45 |
| DMNH 46236 | No | Yes | *Polioptila melanura* | 33.51 | -111.9 |
| DMNH 52160 | No | Yes | *Polioptila melanura* | 25.45 | -101.32 |
| DMNH 7666 | No | Yes | *Polioptila melanura* | 33.45 | -112.07 |
| LSUMZ 34177 | Yes | No | *Polioptila melanura* | 31.5625 | -105.1802 |
| LSUMZ 61914 | Yes | No | *Polioptila melanura* | 34.8 | -116.1 |
| MSB 19565 | Yes | No | *Polioptila melanura* | 32.6870174 | -108.9676817 |
| MSB 29256 | Yes | No | *Polioptila melanura* | 32.821172 | -107.29522 |
| MSB 29257 | Yes | Yes | *Polioptila melanura* | 32.821172 | -107.29522 |
| MSB 29366 | Yes | No | *Polioptila melanura* | 33.260584 | -113.883224 |
| MSB 29367 | Yes | No | *Polioptila melanura* | 33.260584 | -113.883224 |
| MSB SKIN 14033 | No | Yes | *Polioptila melanura* | 32.05 | -109.04 |
| MSB SKIN 18796 | No | Yes | *Polioptila melanura* | 32.62 | -106.93 |
| MSB SKIN 19566 | No | Yes | *Polioptila melanura* | 32.69 | -108.97 |
| UWBM 100358 | Yes | No | *Polioptila melanura* | 32.84361111 | -114.4469444 |
| UWBM 108999 | Yes | No | *Polioptila melanura* | 32.84361111 | -114.4469444 |
| UWBM 111997 | Yes | No | *Polioptila melanura* | 32.84361111 | -114.4469444 |
| UWBM 112009 | Yes | No | *Polioptila melanura* | 32.84361111 | -114.4469444 |
| UWBM 112130 | Yes | No | *Polioptila melanura* | 29.60194444 | -103.0013889 |
| UWBM 112138 | Yes | No | *Polioptila melanura* | 29.60194444 | -103.0013889 |
| UWBM 112658 | Yes | No | *Polioptila melanura* | 32.84361111 | -114.4469444 |
| AMNH J-0004* | Yes | Yes | *Phainopepla nitens* | 32.29166 | -112.85538 |
| AMNH J-0011* | Yes | Yes | *Phainopepla nitens* | 32.29166 | -112.85538 |
| AMNH J-0012* | Yes | Yes | *Phainopepla nitens* | 32.29166 | -112.85538 |
| AMNH J-0013* | Yes | No | *Phainopepla nitens* | 32.29166 | -112.85538 |
| AMNH J-0015* | Yes | Yes | *Phainopepla nitens* | 32.29166 | -112.85538 |
| AMNH SKIN 028059 | No | Yes | *Phainopepla nitens* | 32.98 | -111.33 |
| AMNH SKIN 053282 | No | Yes | *Phainopepla nitens* | 34.56 | -111.85 |
| AMNH SKIN 053290 | No | Yes | *Phainopepla nitens* | 34.56 | -111.85 |
| AMNH SKIN 053301 | No | Yes | *Phainopepla nitens* | 33.3 | -110.4 |
| AMNH SKIN 053309 | No | Yes | *Phainopepla nitens* | 34.56 | -111.85 |
| AMNH SKIN 092562 | No | Yes | *Phainopepla nitens* | 26 | -105.8 |
| AMNH SKIN 092565 | No | Yes | *Phainopepla nitens* | 26 | -105.8 |
| AMNH SKIN 092567 | No | Yes | *Phainopepla nitens* | 26 | -105.8 |
| AMNH SKIN 092569 | No | Yes | *Phainopepla nitens* | 26 | -105.8 |
| AMNH SKIN 092570 | No | Yes | *Phainopepla nitens* | 26 | -105.8 |
| AMNH SKIN 706908 | No | Yes | *Phainopepla nitens* | 22.4 | -100.3 |
| AMNH SKIN 706909 | No | Yes | *Phainopepla nitens* | 22.4 | -100.3 |
| AMNH SKIN 706910 | No | Yes | *Phainopepla nitens* | 22.4 | -100.3 |
| CU 14455 | No | Yes | *Phainopepla nitens* | 30.04 | -103.27 |
| LSUMZ 52747 | Yes | No | *Phainopepla nitens* | 30.6 | -104 |
| LSUMZ 64140 | Yes | No | *Phainopepla nitens* | 30.6 | -104 |
| LSUMZ 64141 | Yes | No | *Phainopepla nitens* | 30.6 | -104 |
| LSUMZ 64233 | Yes | No | *Phainopepla nitens* | 29.5 | -103 |
| MSB 29255 | Yes | Yes | *Phainopepla nitens* | 32.82011667 | -107.30325 |
| MSB 29847 29841? | Yes | No | *Phainopepla nitens* | 32.82011667 | -107.30325 |
| MSB 39010 | Yes | No | *Phainopepla nitens* | 34.003226 | -106.990678 |
| MSB 40683 | Yes | No | *Phainopepla nitens* | 33.62378333 | -107.0090167 |
| MSB 44074 | Yes | Yes | *Phainopepla nitens* | 33.632 | -107.0062667 |
| MSB 44075 | Yes | Yes | *Phainopepla nitens* | 33.68205 | -107.0011667 |
| MSB SKIN 15840 | No | Yes | *Phainopepla nitens* | 33.37 | -105.94 |
| MSB SKIN 349 | No | Yes | *Phainopepla nitens* | 31.41 | -108.78 |
| UMMZ SKIN 65240 | No | Yes | *Phainopepla nitens* | 29.85 | -103.51 |
| UWBM 115563 | Yes | No | *Phainopepla nitens* | 34.095333 | -114.0965 |
| UWBM 115608 | Yes | No | *Phainopepla nitens* | 32.8395 | -114.4415 |
| UWBM 77527 | Yes | No | *Phainopepla nitens* | 31.7994 | -110.7942 |
| UWBM 77633 | Yes | No | *Phainopepla nitens* | 31.523 | -108.9787 |
| UWBM 77634 | Yes | No | *Phainopepla nitens* | 31.523 | -108.9787 |
| AMNH 244159* | Yes | No | *Cardinalis sinuatus* | 29.56 | -103.81 |
| AMNH 244218* | Yes | No | *Cardinalis sinuatus* | 29.56 | -103.81 |
| AMNH 244232* | Yes | No | *Cardinalis sinuatus* | 29.56 | -103.81 |
| AMNH DOT-3731 | Yes | No | *Cardinalis sinuatus* | 31.9 | -109.16 |
| AMNH J-0080* | Yes | Yes | *Cardinalis sinuatus* | 32.11 | -107.6 |
| AMNH J-0132* | Yes | Yes | *Cardinalis sinuatus* | 30.75 | -105 |
| AMNH J-0961* | Yes | No | *Cardinalis sinuatus* | 32.44 | -111.46 |
| AMNH J-0962* | Yes | No | *Cardinalis sinuatus* | 32.44 | -111.46 |
| AMNH J-0999* | Yes | No | *Cardinalis sinuatus* | 31.65 | -108.33 |
| AMNH SKIN 084270 | No | Yes | *Cardinalis sinuatus* | 26.3 | -98.8 |
| AMNH SKIN 084271 | No | Yes | *Cardinalis sinuatus* | 26.3 | -98.8 |
| AMNH SKIN 084272 | No | Yes | *Cardinalis sinuatus* | 26.3 | -98.8 |
| AMNH SKIN 084274 | No | Yes | *Cardinalis sinuatus* | 26.3 | -98.8 |
| AMNH SKIN 084311 | No | Yes | *Cardinalis sinuatus* | 26.3 | -98.8 |
| AMNH SKIN 084312 | No | Yes | *Cardinalis sinuatus* | 26.3 | -98.8 |
| AMNH SKIN 084315 | No | Yes | *Cardinalis sinuatus* | 26.3 | -98.8 |
| AMNH SKIN 084323 | No | Yes | *Cardinalis sinuatus* | 31.2 | -98.3 |
| AMNH SKIN 084330 | No | Yes | *Cardinalis sinuatus* | 26.21 | -98.33 |
| AMNH SKIN 760883 | No | Yes | *Cardinalis sinuatus* | 32.2 | -110.87 |
| AMNH SKIN 760885 | No | Yes | *Cardinalis sinuatus* | 32.26 | -110.87 |
| AMNH SKIN 760886 | No | Yes | *Cardinalis sinuatus* | 32.26 | -110.87 |
| AMNH SKIN 760887 | No | Yes | *Cardinalis sinuatus* | 32.26 | -110.87 |
| AMNH SKIN 760888 | No | Yes | *Cardinalis sinuatus* | 32.26 | -110.87 |
| AMNH SKIN 760890 | No | Yes | *Cardinalis sinuatus* | 27 | -98 |
| DMNH 13114 | No | Yes | *Cardinalis sinuatus* | 25.67 | -101 |
| DMNH 23501 | No | Yes | *Cardinalis sinuatus* | 25.67 | -101 |
| DMNH 23503 | No | Yes | *Cardinalis sinuatus* | 25.67 | -101 |
| DMNH 48438 | No | Yes | *Cardinalis sinuatus* | 25.43 | -100.94 |
| LSUMZ 17193 | Yes | No | *Cardinalis sinuatus* | 31.56 | -105.18 |
| MSB 18064 | Yes | No | *Cardinalis sinuatus* | 32.05 | -109.04 |
| MSB 24457 | Yes | No | *Cardinalis sinuatus* | 29.99 | -103.56 |
| MSB 24612 | Yes | No | *Cardinalis sinuatus* | 29.99 | -103.56 |
| MSB 25201 | Yes | No | *Cardinalis sinuatus* | 32.06 | -112.71 |
| MSB SKIN 14262 | No | Yes | *Cardinalis sinuatus* | 34.18 | -103.35 |
| MSB SKIN 21242 | No | Yes | *Cardinalis sinuatus* | 32.33 | -103.24 |
| MSB SKIN 21243 | No | Yes | *Cardinalis sinuatus* | 32.11 | -103.24 |
| MSB SKIN 3707 | No | Yes | *Cardinalis sinuatus* | 32.13 | -103.49 |
| MSB SKIN 5200 | No | Yes | *Cardinalis sinuatus* | 32.38 | -103.72 |
| MSB SKIN 6480 | No | Yes | *Cardinalis sinuatus* | 32.05 | -109.04 |
| MSB SKIN 8810 | No | Yes | *Cardinalis sinuatus* | 32.05 | -109.04 |
| MSB SKIN 8813 | No | Yes | *Cardinalis sinuatus* | 33.39 | -103.81 |
| TCWC 16314 | Yes | No | *Cardinalis sinuatus* | 31.4 | -102.37 |
| TCWC 16576 BDM1900 | Yes | No | *Cardinalis sinuatus* | 30.16 | -103.23 |
| UWBM 100163 | Yes | No | *Cardinalis sinuatus* | 29.6 | -103 |
| UWBM 103346 | Yes | No | *Cardinalis sinuatus* | 31.48 | -111.34 |
| UWBM 105429 | Yes | No | *Cardinalis sinuatus* | 29.6 | -103 |
| UWBM 109277 | Yes | No | *Cardinalis sinuatus* | 29.6 | -103 |
| UWBM 109278 | Yes | No | *Cardinalis sinuatus* | 29.6 | -103 |
| UWBM 77548 | Yes | No | *Cardinalis sinuatus* | 32.21 | -109.18 |
| UWBM 77718 | Yes | No | *Cardinalis sinuatus* | 32.21 | -109.18 |
| UWBM 77780 | Yes | No | *Cardinalis sinuatus* | 32.21 | -109.18 |
| UWBM 77781 | Yes | No | *Cardinalis sinuatus* | 32.21 | -109.18 |

Supplementary Table 6: Table of reference genomes used in this study.

| **Reference Genome** | **Genome Species** | **Citations** | **Species Mapped To** |
| --- | --- | --- | --- |
| GCF_000691975.1 | *Corvus brachyrhynchos* | Zhang et al 2014 | *Vireo bellii* |
| GCF_001522545.2 | *Parus major* | Laine et al 2016 Laine et al 2019 | *Auriparus flaviceps* |
| GCF_000385455.1 | *Zonotrichia albicollis* | Romanov et al 2011 | *Amphispiza bilineata Melozone fusca* |
| GCF_000277835.1 | *Geospiza fortis* | Zhang et al 2014 | *Cardinalis sinuatus* |
| GCF_001447265.1 | *Sturnus vulgaris* | Starling Genome Consortium, 2015 | *Campylorhynchus brunneicapillus Toxostoma crissale*  *Toxostoma curvirostre Polioptila melanura*  *Phainopepla nitens* |

Supplementary Table 7: Best models selected to generate the ecological niche models for each species. AUC=area under curve. Train=training dataset. Test=average values across all test datasets. OR=average test 10% omission rate. Reg=regularization parameter. LQ=linear and quadratic.

| **Species** | **Features** | **Reg** | **Train  AUC** | **Test  AUC** | **OR** | **Parameters** |
| --- | --- | --- | --- | --- | --- | --- |
| *Toxostoma crissale* | LQ | 1 | 0.97 | 0.96 | 0.07 | 18 |
| *Amphispiza bilineata* | LQ | 0.5 | 0.91 | 0.90 | 0.14 | 25 |
| *Toxostoma curvirostre* | LQ | 2 | 0.90 | 0.87 | 0.13 | 16 |
| *Auriparus flaviceps* | LQ | 0.5 | 0.94 | 0.92 | 0.20 | 25 |
| *Vireo bellii* | LQ | 0.5 | 0.87 | 0.83 | 0.14 | 28 |
| *Campylorhynchus brunneicapillus* | LQ | 0.5 | 0.93 | 0.91 | 0.22 | 23 |
| *Cardinalis sinuatus* | LQ | 1.5 | 0.93 | 0.91 | 0.23 | 16 |
| *Melozone fusca* | LQ | 0.5 | 0.93 | 0.89 | 0.26 | 27 |
| *Polioptila melanura* | LQ | 0.5 | 0.96 | 0.95 | 0.24 | 23 |
| *Phainopepla nitens* | LQ | 1.5 | 0.96 | 0.95 | 0.23 | 17 |
